# On Long-Term Species Coexistence in Five-Species Evolutionary Spatial Cyclic Games with Ablated and Non-Ablated Dominance Networks

**DOI:** 10.1101/2024.09.27.615336

**Authors:** Dave Cliff

**Affiliations:** School of Engineering Mathematics & Technology, University of Bristol, BS8 1UB, U.K.

**Keywords:** Biodiversity, Cyclic Competition, Asymmetric Interaction, Species Coexistence, Evolutionary Spatial Games, Rock-Paper-Scissors, Replication

## Abstract

I present a replication and, to some extent, a refutation of key results published by Zhong, Zhang, Li, Dai, & Yang in their 2022 paper “Species coexistence in spatial cyclic game of five species” (*Chaos, Solitons and Fractals*, 156: 111806), where ecosystem species coexistence was explored via simulation studies of the evolutionary spatial cyclic game (Escg) Rock-Paper-Scissors-Lizard-Spock (Rpsls) with certain predator-prey relationships removed from the game’s “interaction structure”, i.e. with specific arcs ab-lated in the Escg’s dominance network, and with the Escg run for 10^5^ Monte Carlo Steps (mcs) to identify its asymptotic behaviors. I replicate the results presented by Zhong et al. for interaction structures with one, two, three, and four arcs ablated from the dominance network. I then empiri-cally demonstrate that the dynamics of the Rpsls Escg have sufficiently long time constants that the true asymptotic outcomes can often only be identified after running the ablated Escg for 10^7^mcs or longer, and that the true long-term outcomes can be markedly less diverse than those reported by Zhong et al. as asymptotic. Finally I demonstrate that, when run for sufficiently many mcs, the original unablated Rpsls system exhibits essentially the same asymptotic outcomes as the ablated Rpsls systems, and in this sense the only causal effect of the ablations is to alter the time required for the system to converge to the long-term asymptotic states that the unablated system eventually settles to anyhow.

**Graphical Abstract:** 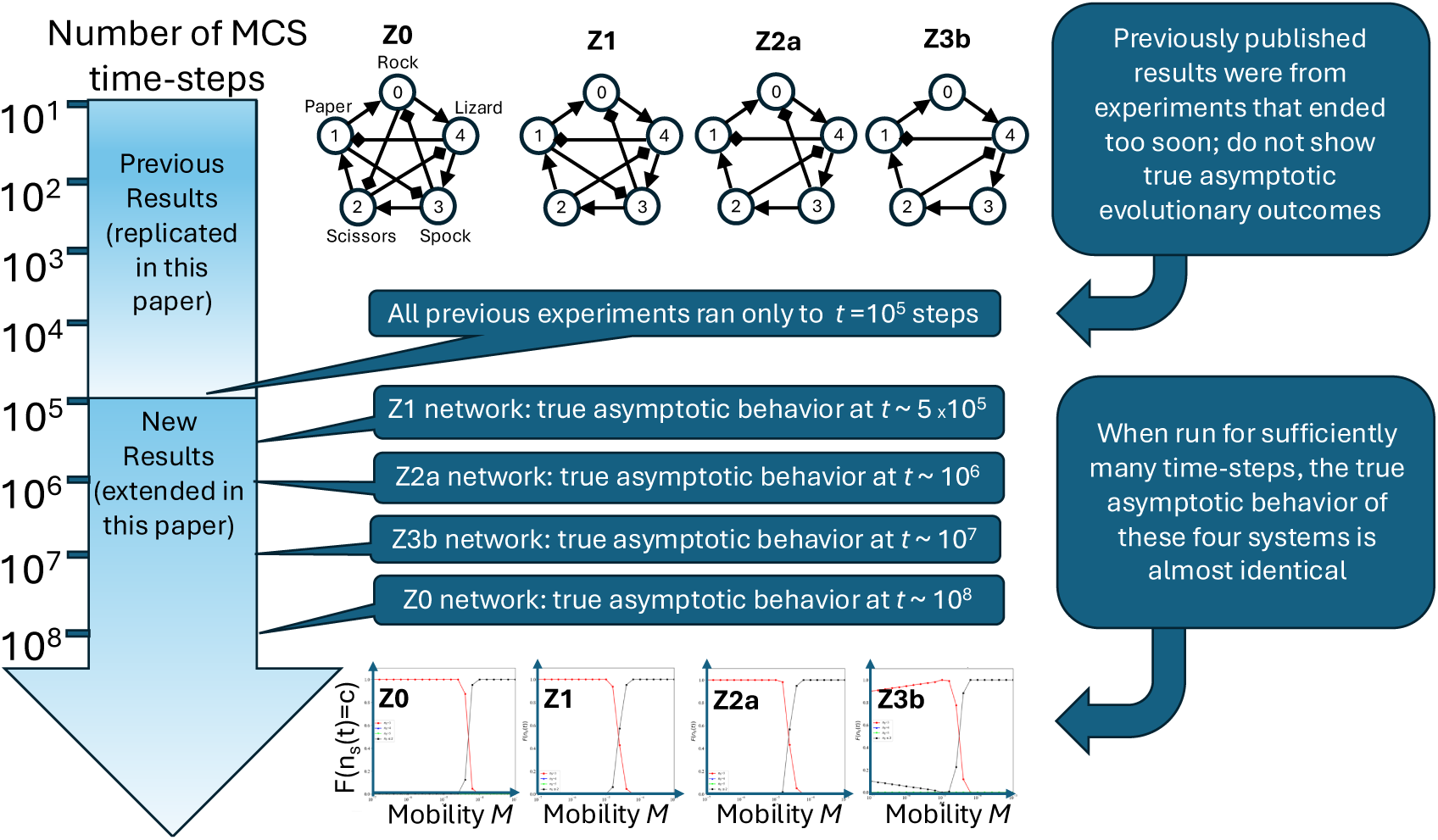

**Highlights:**

- I replicate key results from Zhong et al. (2022) where biodiversity was explored via the game Rock-Paper-Scissors-Lizard-Spock (Rpsls).
- Zhong et al. reported results from Rpsls games where specific predatorprey interactions were ablated from the game’s dominance network.
- My replication reveals problems in Zhong et al.’s design of experiments.
- Zhong et al. did not run their simulations for sufficiently long to reveal the true asymptotic behavior of the ablated Rpsls systems.
- Zhong et al. did not present control outcomes from the unablated Rp-sls system, so there is no baseline data for comparison to the treatment outcomes.
- I present results from simulations that are run for 100 to 1000 times longer than the experiments reported by Zhong et al., thereby revealing the true asymptotic behaviors of the system.
- The asymptotic outcomes are remarkably uniform – practically indistinguishable – in the cases where one, two, or three arcs are ablated from the Rpsls dominance network.
- My asymptotic results for the baseline original unablated system are also very similar to those for the one-two- and three-ablation systems.
- My results question whether the ablations have any effect other than speeding the system’s convergence to its eventual asymptotic state.
- Results from Zhong et al.’s four-ablation system do not fit so well with the lower-ablation-count systems: potential reasons for this, and avenues for further research on it, are discussed.

## 1. Introduction

In 2022 a research paper by Zhong, Zhang, Li, Dai, & Yang titled “Species coexistence in spatial cyclic game of five species” was published in the journal *Chaos, Solitons and Fractals* (2022, 156: 111806) [1]. In their paper, Zhong et al. explored the frequency distribution of outcomes from simulations studies of a minimal model of ecosystem species coexistence, based on the evolutionary spatial cyclic game (Escg) Rock-Paper-Scissors-Lizard-Spock (Rpsls) with certain predator-prey relationships removed from the game’s “interaction structure”, i.e. with specific arcs ablated in the Escg’s dominance network.^1^ Zhong et al. identified the asymptotic behaviors of the ablated Escg by running simulations for 10^5^ Monte Carlo Steps (mcs: a major time-step in the simulation process, explained further in Section 2, below). In this paper I replicate the results presented by Zhong et al. for interaction structures with one, two, three, and four arcs ablated from the dominance network. I then empirically demonstrate that the dynamics of the ablated Rpsls Escg have sufficiently long time constants that the true asymptotic outcomes can often only be identified after simulations running for 10^7^mcs or longer (i.e., 100× the durations used by Zhong et al.), and that the true long-term outcomes can be markedly different from those reported by Zhong et al. as asymptotic. Following this, I demonstrate that, when run for sufficiently many mcs, the original *unablated* Rpsls system exhibits almost exactly the same asymptotic outcomes as the ablated Rpsls systems, and hence in this sense the only causal effect of the dominance-network arc ablations is to reduce the time required for the system to converge on the long-term asymptotic states that the unablated system eventually settles to anyhow.

Section 2 gives further details of the background to this work, the text of that section being reproduced essentially verbatim from [4]. Section 3 then gives a detailed summary of the model and simulation methods as used by Zhong et al. in [1]. After that, Section 4 shows visualization and analysis of results from my own simulation experiments which accurately replicate Zhong et al.’s results presented in [1]. The next two sections then present a detailed critique of those results: Section 5 argues that the observations made and conclusions drawn by Zhong et al. are based on data from experiments that had not been run for sufficiently many time-steps, and that some features in the graphs plotted by Zhong et al., features they highlighted as noteworthy, are in fact nothing more than non-asymptotic artefacts from decaying transients in the system dynamics which appear because the experiments have not been run for long enough; Section 6 then goes on to present results from new simulation experiments, not reported by Zhong et al., where the original unablated Rpsls dominance network is used, to give a proper baseline reference that the results from the ablated networks can be compared to: somewhat surprisingly, the original unablated Rpsls system exhibits what are, qualitatively, essentially identical asymptotic behaviors to those of the ablated one-, two, and three-ablation Rpsls systems reported by Zhong et al. Finally, in Section 7, results from the four-ablation system explored by Zhong et al. are discussed separately, because they seem not to show the same long transients as the one-, two-, and three-ablation cases, and in that sense the four-ablation results remain something of a mystery. All of these results are collectively discussed further in Section 8, and conclusions are drawn in Section 9.

For completeness, the appendices show numerous plots of data that more fully illustrate the points made in my critique but which did not fit naturally into the narrative of Sections 5 and 6. Python source-code developed for the simulation experiments reported here is being made freely available via the MIT Open-Source License, for download from Github.^2^ The data generated for this paper, in uncompressed csv-format files, occupies approximately two terabytes of storage: a small amount of illustrative sample data is being made available on the GitHub source-code repository.

## 2. Background

There is a well-established body of peer-reviewed research literature which explores issues in ecosystems stability, biodiversity, and co-evolutionary dynamics via computationally intensive simulations of minimally simple models of multiple interacting biological species. Landmark papers in this field were published in 2007–08 by Reichenbach, Mobilia, and Frey [5, 6, 7], who extended the previous non-spatial model of May and Leonard [8] by modeling each species as a time-varying number of discrete individuals, where at any one time each individual occupies a particular cell in a regular rectangular lattice or grid of cells, and can *move* from cell to cell over time — that is, the individuals are *spatially located* and *mobile*. Individuals can also, under the right circumstances, *reproduce* (asexually, cloning a fresh individual of the same species into an adjacent empty cell on the lattice); and they can also *compete* with individuals in neighbouring cells. Different authors use different phrasings to explain the inter-species competition, but it is common to talk in terms of predator-prey dynamics: that is, each species is predator to (i.e., *dominates*) some specified set of other species, and is in turn also prey to (i.e., is *dominated* by) some set of other species.

The population dynamics are determined to a large extent by the model’s *dominance network*, a directed graph (digraph) where each node in the network represents one of the species in the model, and a directed edge (i.e., an arc) from the node for species *S_i_* to the node for species *S_j_*_:*j*_*_/_*_=*i*_ denotes that *S_i_* dominates *S_j_*. To exhibit interesting long-term dynamics, the dominance network must contain at least one cycle (i.e., a path from some species *S_i_* to some *S_j_* traced by traversing edges in the directions of the arrows, potentially passing through some number of intermediate species’ nodes, where at *S_j_* there then exists an as-yet-untraversed edge back to *S_i_*). Under the constraint that no species can be both predator and prey to another species at the same time, the smallest dominance network of interest is a minimal three-node cycle, which represents the intransitive dominance hierarchy of the simple hand-gesture game Rock-Paper-Scissors (RPS), as illustrated in Figure 1. In RPS-based models, the three species are *R* (rock), *P* (paper), and *S* (scissors) and when two neighboring individuals compete the rules are as follows: if they are both the same species, the competition is a draw and nothing else happens; but otherwise *R* kills *S*, *S* kills *P*, and *P* kills *R*, with the cell where the killed individual was located being set to empty, denoted by . Because the individuals in these models are spatially located on a lattice, and because the inter-species competition is determined by having pairs of individuals play RPS-like games with cyclic dominance digraphs, this class of co-evolutionary population dynamics models is often referred to as *evolutionary spatial cyclic games* (Escgs).

**Figure 1:**
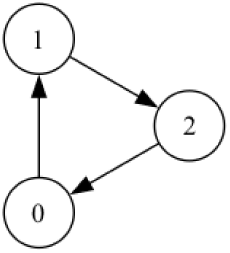
Dominance network, a directed graph or *digraph*, for the three-species Rock-Paper-Scissors (RPS) game. Species *S_i_* are denoted by nodes numbered by index *i* 1, 2, 3, with directed edges running from the dominator (“predator”) species to the dominated (“prey”’) species. There are multiple *labelings* of this graph (e.g.: (*S*_1_=*R, S*_2_=*P, S*_3_=*S*); (*S*_1_=*P, S*_2_=*S, S*_3_=*R*); *. . .*) but if one graph can be turned into another purely by rearranging the node labels then those two graphs are topologically equivalent or *isomorphic*. All possible labelings of the RPS digraph are isomorphic with each other, so there is only one isomorphically unique RPS digraph.

Escgs are inherently stochastic and to generate rigorous results it is often necessary to simulate Escg systems many times, aggregating results over many independent and identically distributed (IID) repetitions of the system evolving over time. At the core of these simulations is the *Elementary Step* (ES), in which one or two agents are chosen at random to either compete to the death, or to reproduce, or to move location. Escg studies typically involve executing trillions of ESs and hence the computational efficiency of the core ES algorithm is a key concern: for further discussion of this, see [4].

Almost all simulations of co-evolutionary population dynamics via Escgs are simple discrete-time systems that are technically unchallenging to write a program for, and are strongly reminiscent of – but not identical to – cellular automata (see e.g. [9]). The lattice/grid needs to first be set up, i.e. its dimensions and initial conditions at the first time-step need to be specified. Each cell in the grid is either empty, or contains exactly one individual organism, and each individual is a member of exactly one of the model’s set of species. If the number of species in the model is denoted by *N_s_*, one common style of initialisation is to assign one individual to every cell in the grid, with that individual’s species being an equiprobable choice from the set of available species (i.e., choose species *S_i_* with probability 1*/N_s_*; *i*). The modeller also needs to specify the dimensionality of the lattice, and its *length* (i.e., number of cells) along each dimension. In almost all of the published work in this field, the lattice is two-dimensional and square, so its extent is defined by a single system hyperparameter: the side-length (conventionally denoted by *L*). The total number of cells in the lattice (conventionally denoted by *N*) is hence *N* = *L*^2^. Working with 2D lattices has the advantage that the global state of the system can be readily visualised as a snapshot at time *t* as a color-coded or gray-shaded 2D image, with each species in the model assigned its own specific color or gray-scale value, and animations can easily be produced visualising the change in the system state over time.

In the literature on Escgs, authors often make the distinction between two scales of time-step in the simulation. At the very core of the simulation process is a loop that iterates over a number of *elementary steps* (ESs), the finest grain of time-step; and then some large number of consecutive ESs is counted as what is conventionally referred to as a *Monte Carlo Step* (MCS).

In a single ES, one individual cell (denoted *c_i_*) is chosen at random, and then one of its immediately neighboring cells (denoted *c_n_*) is also chosen at random: in almost all of the literature on 2D lattice model Escgs, the set of neighbours is defined as the 4-connected von Neumann neighborhood rather than the 8-connected Moore neighborhood commonly used in cellular automata research (although e.g. [10, 11, 12] used the Moore neighborhood in their lattice models), and the work reported here uses von Nuemann. There seems to be no firm convention on whether to use periodic boundary conditions (also known as toroidal wrap-around) or “walled garden” no-flux boundary conditions (such that cells at the edges and corners of the lattice have a correspondingly reduced neighbour-count) – some authors use periodic, others no-flux. The results presented in this paper come from simulations with no-flux boundary conditions.

In each ES one of three possible actions occurs: *competition*, *reproduction*, or *movement*, and the probabilities of each of these three actions occurring per ES is set by system parameters *µ, σ,* and *ɛ*, respectively (this is explained in more precise detail later, in Section 3). *Competition* involves the individuals at *c_i_* and *c_n_* interacting according to the rules of the cyclic game, resulting in either a draw or one of the individuals losing, in which case it is deleted from its cell, replaced by Ø; *reproduction* occurs when one of *c_i_* or *c_n_* holds Ø, the empty cell being filled by a new individual of the same species as the nonempty neighbor; and *movement* simply swaps the contents of *c_i_* and *c_n_*.

Because, in the original formulation, each ES involves only one of the three possible actions (competition, reproduction, or movement) occurring for a single cell, a Monte Carlo Step (mcs) is conventionally defined as a sequence of *N* consecutive ESs, the rationale being that, on the average, each cell in the lattice will be randomly chosen once per mcs, and hence that, again on the average, every cell in the grid has the potential to change once between any two successive mcss. Most published research on this type of model uses mcs as the unit of time when plotting time-series graphs illustrating the temporal evolution of the system, and I follow that convention here. Some authors (e.g. [13]) don’t refer to mcs but instead talk of each sequence of *N* consecutive ESs in their Escg as one new *generation*.

In their seminal papers, Reichenbach et al. [5, 6, 7] studied 2D lattice systems where interspecies competition was via *N_s_*=3 RPS games, with *µ*=*σ*=1.0, and where *L* ranged from 100 to 500, and they showed and explained how the overall system dynamics result in emergence of one or more temporally and spatially coherent interlocked *spiral waves*. The specific nature of the wave-patterning, i.e. the size and number of spiral waves seen in the system-snapshot images, depended on a *mobility* measure *M* =*ɛ/*2*N*, which is proportional to the expected area of lattice explored by a single agent per mcs.

In the years since publication of [5, 6, 7], many papers have been published that explore the dynamics of such co-evolutionary spatial RPS models. For examples of recent publications exploring a range of issues in the three-species RPS Escg, see: [14, 15, 16, 17, 18, 19, 20, 21, 22, 23]; and [24].

More recently, various authors have reported experiments with a closely related system where *N_s_*=5: this game is known as Rock-Paper-Scissors-Lizard-Spock (Rpsls), an extension of RPS introduced by [25] and subsequently featured in a 2012 episode of the popular US TV show *Big Bang Theory*. The dominance network for the Rpsls game is illustrated in Figure 2 and explained in the caption to that figure. This (and other five-species Escgs) was first explored in the theoretical biology literature by [10, 11, 12]; and RPS-like Escgs with *N_S_* 5 were explored by [26].

The rest of this paper focuses on one recently-published study of the evolutionary dynamics of the Rpsls system when one or more of the arcs in the dominance network are ablated, reducing the number of inter-species interactions: this paper is Zhong et al.’s 2022 paper [1], which is explained in detail in the next section.

## 3. Summary of Zhong et al. (2022)

### 3.1. *The Evolutionary Spatial Cyclic Game (*Escg)

Figure 3 shows the set of ablated Rpsls dominance networks explored by Zhong et al. in their paper: I have labelled each network with an identifier that includes the number of ablated arcs in that network; and for consistency I refer to the original unablated Rpsls dominance network as ‘Z0’.

The lattice is an *L×L* square Cartesian grid of cells. The contents of any cell in the lattice can be either the empty cell, denoted here by, or a natural number *i* representing that an individual agent of species-type *s_i_* ∈ {0, 1*, . . ., N_S_* − 1} occupies that cell. Let C*_NS_* denote the set of all possible cell values in an Escg with *N_S_* species: here, for the five-species Rpsls game, C_5_ = {0, 1, 2, 3, 4, Ø}.

In its simplest form, on each elementary step (ES), an individual cell *c_i_* is chosen at random and then one of its immediately neighboring cells *c_n_* is chosen, also at random: for instance, [23, p.2] state “At each time step, we choose a pair of neighboring sites randomly”. Below, I give an initial definition of the Escg on a 2D square lattice as Algorithm 1, which uses this simple random selection of two neighboring cells; then later, in Section 4, I’ll discuss an improvement to this approach which is given in Algorithm 4.

Let the contents of cell *c_i_* and the contents of cell *c_n_* be denoted by a pair of cell-content values, each in _5_. The update rules for the Escg using the Rpsls dominance network *Z*0 (shown in Figure 2) can then be expressed as follows: cell-to-cell *competition*, which occurs with frequency determined by the parameter *σ*, is defined by Equations 1 to 5:

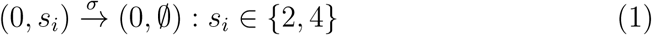

**Figure 2:**
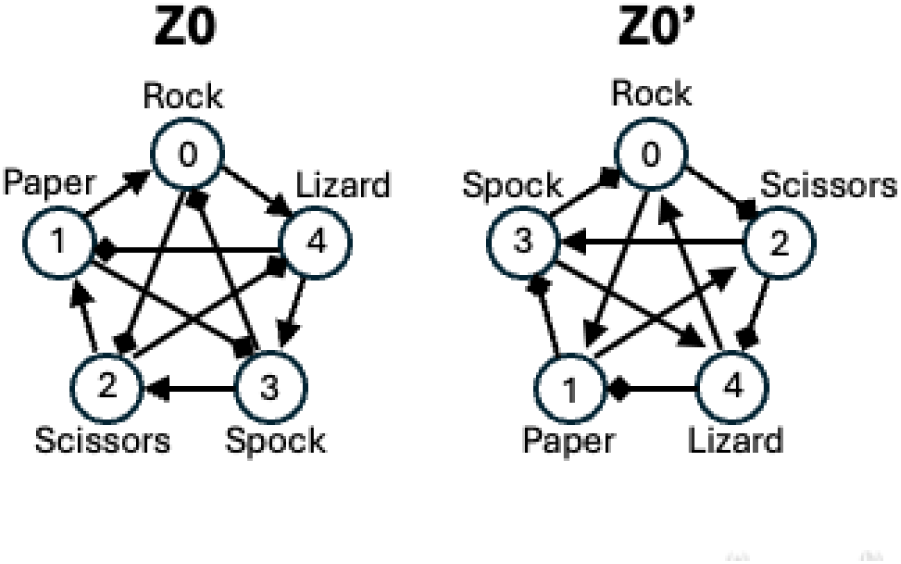
Network Z0 (left) is the Rpsls dominance network as presented in Figure 1(a) of Zhong et al. (2022): the outer pentagon subnetwork formed by triangle-headed arrows is referred to by Zhong et al. as the *spontaneous competition* dominance interactions while the inner pentagram subnetwork formed by diamond-headed arrows is referred to by Zhong et al. as the *alternative competition* dominance interactions. Network Z0’ (right) is topologically equivalent to Z0, despite appearing superficially different. The rules of this game are: scissors cut paper; paper covers rock; rock blunts scissors; scissors decapitates lizard; lizard eats paper; paper disproves Spock; Spock vaporizes rock; rock crushes lizard; lizard poisons Spock; and Spock smashes scissors.

**Figure 3:**
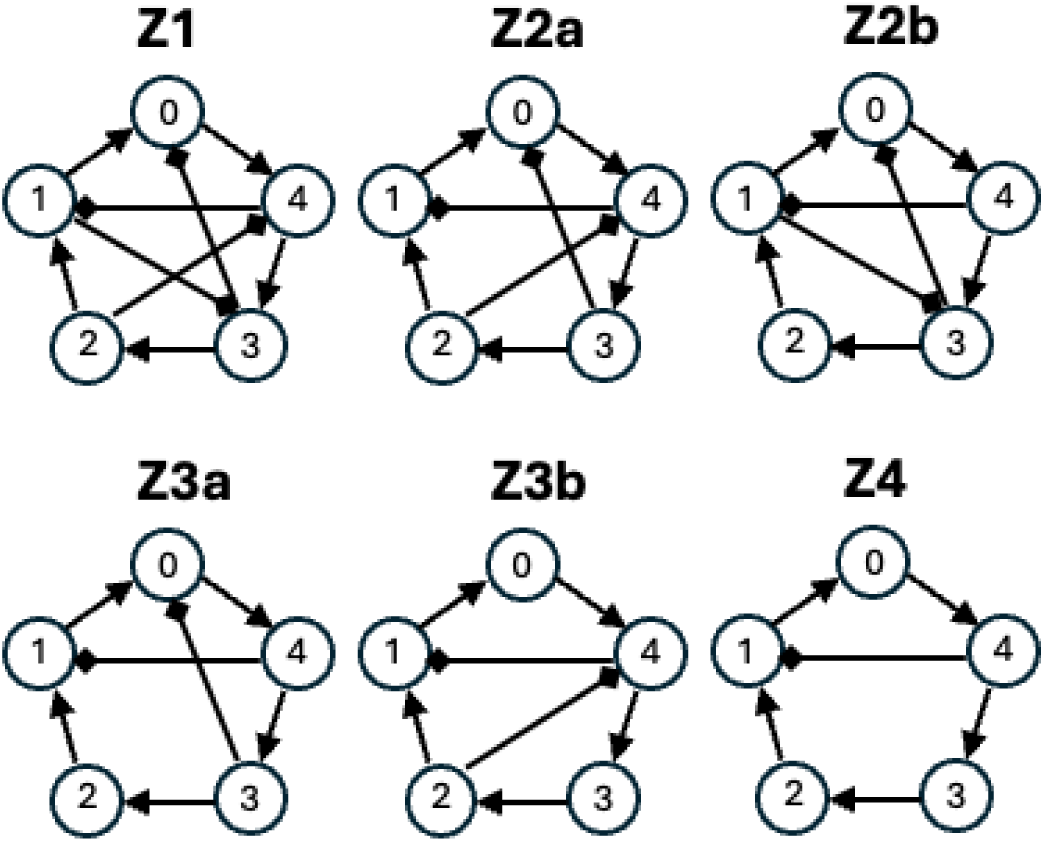
The ablated dominance networks explored in Zhong et al. (2022). Network Z1 is Zhong et al.’s Figure 1(b); Network Z2a is Zhong et al.’s Figure 1(c)-upper; Network Z2b is Zhong et al.’s Figure 1(c)-lower; Network Z3a is Zhong et al.’s Figure 1(d)-upper; Network Z3b is Zhong et al.’s Figure 1(d)-lower; and Network Z4 is Zhong et al.’s Figure 1(e).

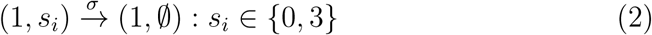

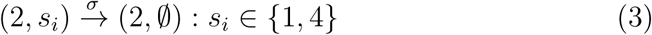

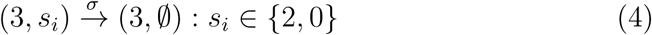

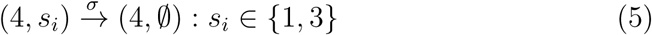

*Reproduction*, where the individual of species *s_i_*in cell *c_i_* clones a copy of itself into an adjacent empty cell, an event which occurs with frequency determined by the parameter *µ*, is as stated in Equation 6:

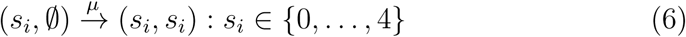

And finally *movement*, where an individual of species *s_i_*in cell *c_i_* swaps location with whatever contents are in its neighbour *c_n_*, an event which occurs with frequency determined by the parameter *ɛ*, is expressed in Equation 7:

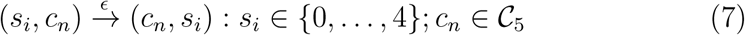

Zhong et al. state that on each elementary step, only one of the three possible types of action occurs, and they also state that each of the three actions “. . . occur between the two selected nodes at the probabilities *σ, µ,* and *ɛ*, respectively”, but that is not strictly correct. Rather, they are each *hyperparameters* (i.e., exogenously imposed parameter-values that are constant during the course of an experiment) which do happen to take values in the range [0.0, 1.0] but (as is explained in more depth in [4]) the three are then combined in a “normalization” process, not explicitly described in [1] but clearly explained by [27], in which each of the three hyperparameters are divided by the sum of the three, so for example let *µ^j^* denote the normalized *µ*, then *µ^j^* = *µ/*(*σ* + *µ* + *ɛ*); *µ* [0.0, 1.0], Hence *µ* is *not* the probability of movement being the action chosen on any one ES, because the actual probability of choosing Move as the next action is given by *µ^j^* which depends not only on *µ* but also on *σ* and *ɛ*. As is common in the literature for this field of study, in all of the experiments reported by Zhong et al., and also in all of the experiments reported here in this paper, *µ* = *σ* = 1.0, without loss of generality; and *ɛ* is a function of *M*, the Escg’s *mobility* hyperparameter, introduced in Section 2.

For the discussions in the remainder of this paper, it is useful to introduce a few extra items of notation additional to those employed by Zhong et al. in their paper: here, let *N_a_* denote the number of arcs ablated from the complete tournament RPSLS dominance network; let *n_s_*(*t*) denote the number of surviving (non-extinct) species in the system at time *t*; let *t*_max_ denote the duration of a simulation experiment, i.e. how many mcs the experiment runs for before terminating, and note that in all of Zhong et al.’s experiments, *t*_max_=10^5^; finally, let *N*_iid_ denote the number of independent and identically distributed (iid) repetitions of a specific simulation experiment with a given set of (hyper-)parameter values – in all of the experiments reported in Zhong et al. (2022), *N*_iid_=500.

In Zhong et al. (2022) the primary mode of analysis and summary of experiment outcomes aggregated over some suitably large number of iid simulation runs was plots of what Zhong et al. referred to as *F*, the frequency of outcome of number of species surviving at the end of the simulation, as a function of *M*, the mobility value used in the simulation: in my notation, this is more specifically denoted by *F* (*n_s_*(*t*_max_)=*c*) vs. *M*, for *c* 1*, . . .,* 5, but for brevity I’ll refer to these simply as “*FvsM* ” plots.

### 3.2. *The* Escg *algorithm*

Let *l* be the 2D lattice of cells such that an individual cell with coordinates (*x, y*) in the lattice is denoted by *l*(*x, y*), and where the individual cell is the *position* of an individual agent *i* in the model, which I’ll denote by *p→_i_*. Let U[*n_lo_, n_hi_*] denote a new random draw from a uniform distribution over the range [*n_lo_, n_hi_*] ⊂ R and similarly let U{*m*_0_*, m*_1_*, . . .*} represent a new uniform (equiprobable) random choice of member from the set {*m*_0_*, m*_1_*, . . .*}.

Assume here the existence of a function RndNeighbour(*l, p→_i_,* N, B) which returns the coordinate pair *p→_n_* = (*x_n_, y_n_*) for a randomly chosen member from the neighbourhood of *p→_i_* = (*x_i_, y_i_*) where specifies the neighborhood function to use (e.g. von Neumann or Moore, etc), and with boundary conditions specified by as either periodic or no-flux.

Assume also that the three possible actions introduced above in Equations 1 to 7 are encoded as three simple functions, each of which take as arguments: the lattice *l*; the lattice position *p→_i_* of the randomly chosen individual cell *c_i_*; and the lattice coordinates *p→_n_* of *c_i_*’s randomly chosen neighboring cell, denoted *c_n_*. Additionally, Compete requires a specification of the dominance network, denoted by D:

- Compete(*l, p→_i_, p→_n_,* D) implements Equations 1 to 5.
- Reproduce(*l, p→_i_, p→_n_*) implements Equation 6.
- Move(*l, p→_i_, p→_n_*) implements Equation 7.

Each of these functions returns the updated lattice *l*. These three functions will be called from within the procedure for a single elementary step, ElStep which takes as arguments the lattice, the position vectors *p→_i_* and *p→_n_* of *c_i_* and *c_n_* respectively, the three hyperparameters *µ, σ,* and *ɛ*, and the dominance network, and returns the updated lattice. The entire algorithm for an instance of the evolutionary spatial cyclic game (Escg) on a square 2D lattice is as shown in Algorithm 1, which calls the PopulateLattice algorithm listed in Algorithm 2, and the Original Elementary Step (OES) algorithm ElStep, listed in Algorithm 3.

As is discussed in more detail in [4], the ElStep algorithm as shown in Algorithm 3 is inefficient in space and in time, because the contents of the randomly-chosen cells at *p→_i_* and *p→_n_* may not be compatible with the randomly chosen action selected within ElStep. For instance, if either *l*(*p→_i_*)=Ø or *l*(*p→_n_*)= and then in ElStep the randomly-chosen action is Compete, no change to the lattice will occur on that elementary step, and so that step becomes a “no-op”, i.e. an operation that does nothing.^3^ The elementary step also becomes a no-op if both *l*(*p→_i_*) Ø and *l*(*p→_n_*) Ø and ElStep randomly selects Reproduce as the action for that step; and also if *l*(*p→_i_*)=Ø and *l*(*p→_n_*)=Ø then ElStep is again a no-op, for all three possible actions.

**Algorithm 1.**
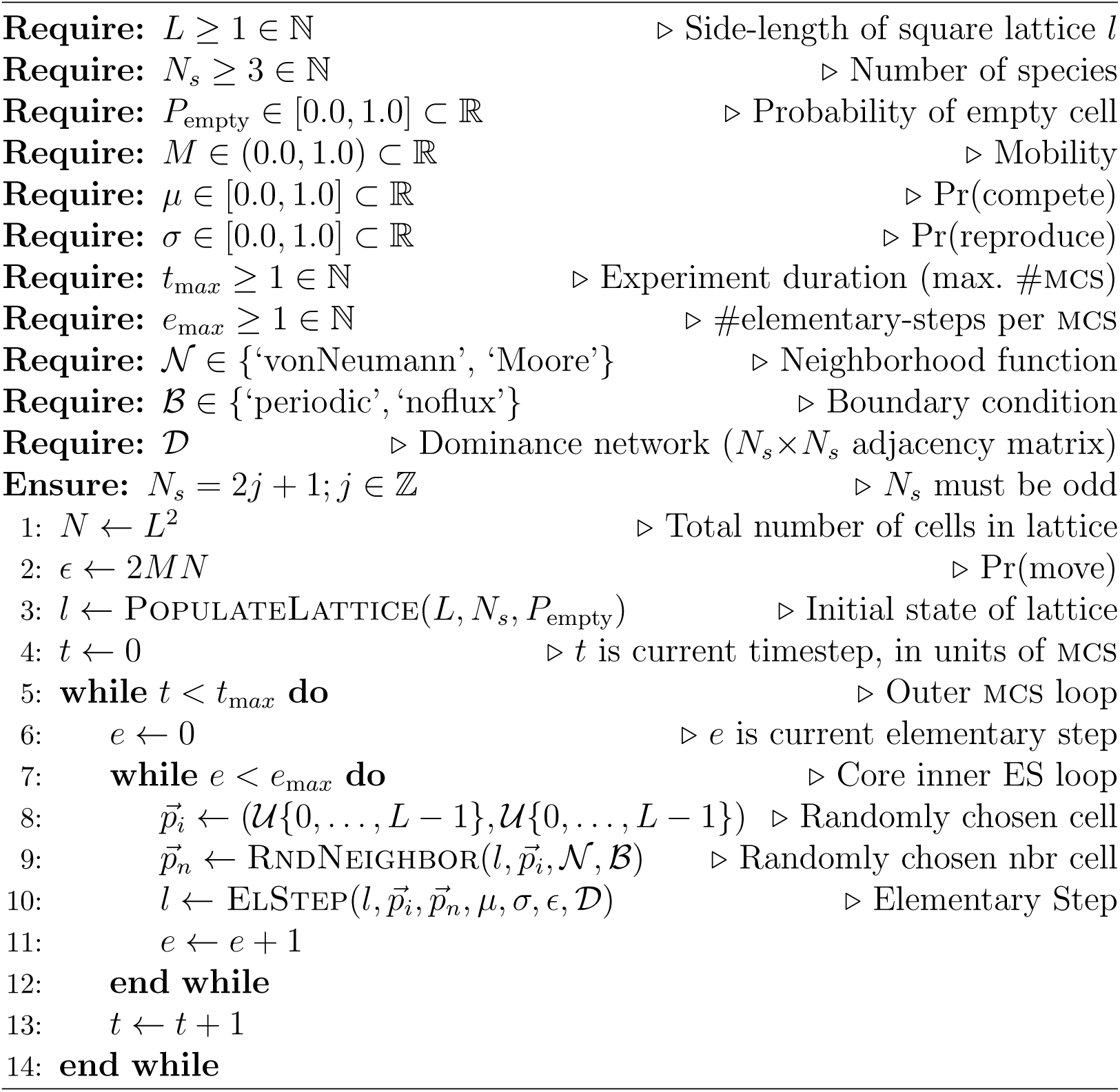
Evolutionary Spatial Cyclic Game (Escg) – 2D Square

**Algorithm 2.**
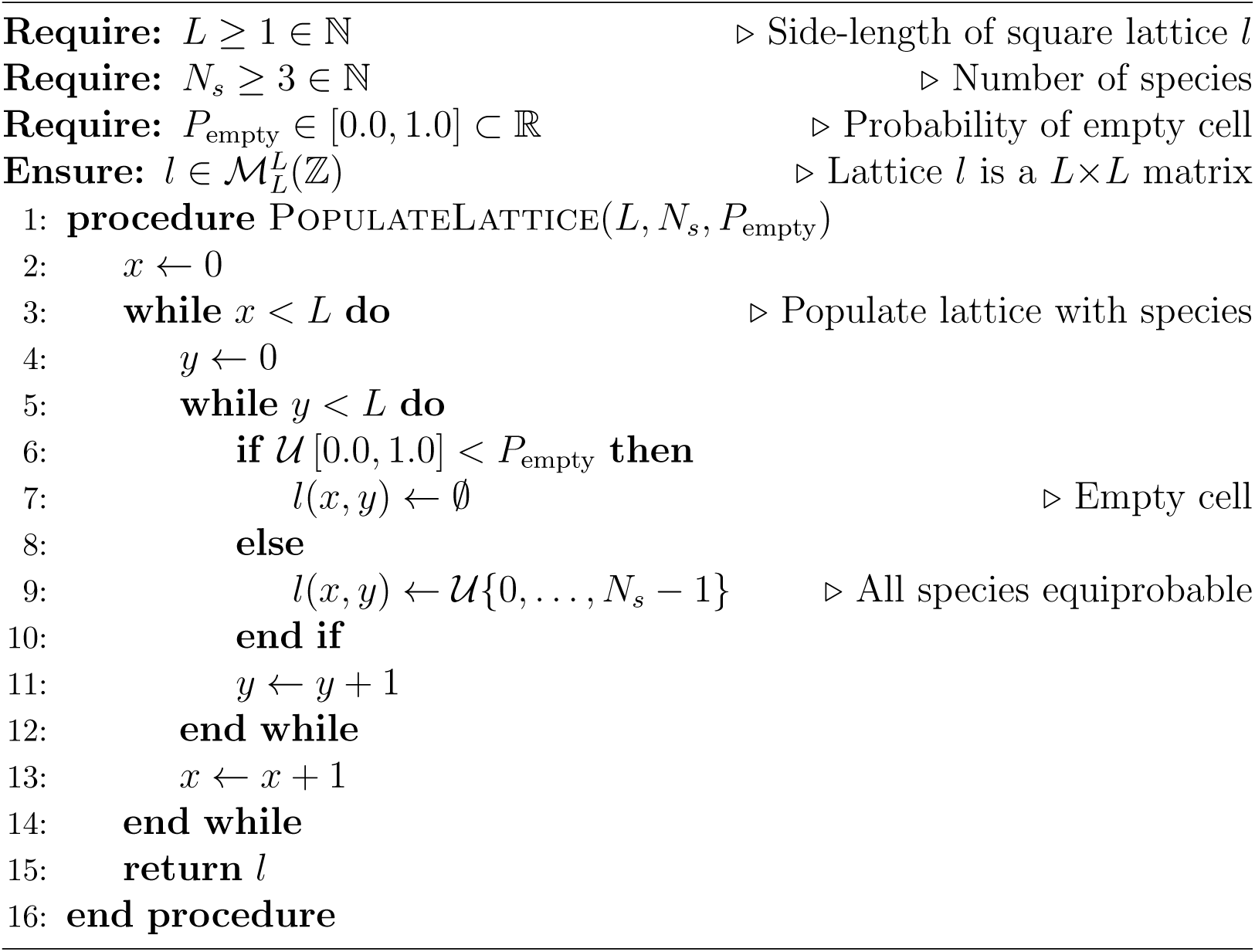
Populating the square lattice

**Algorithm 3.**
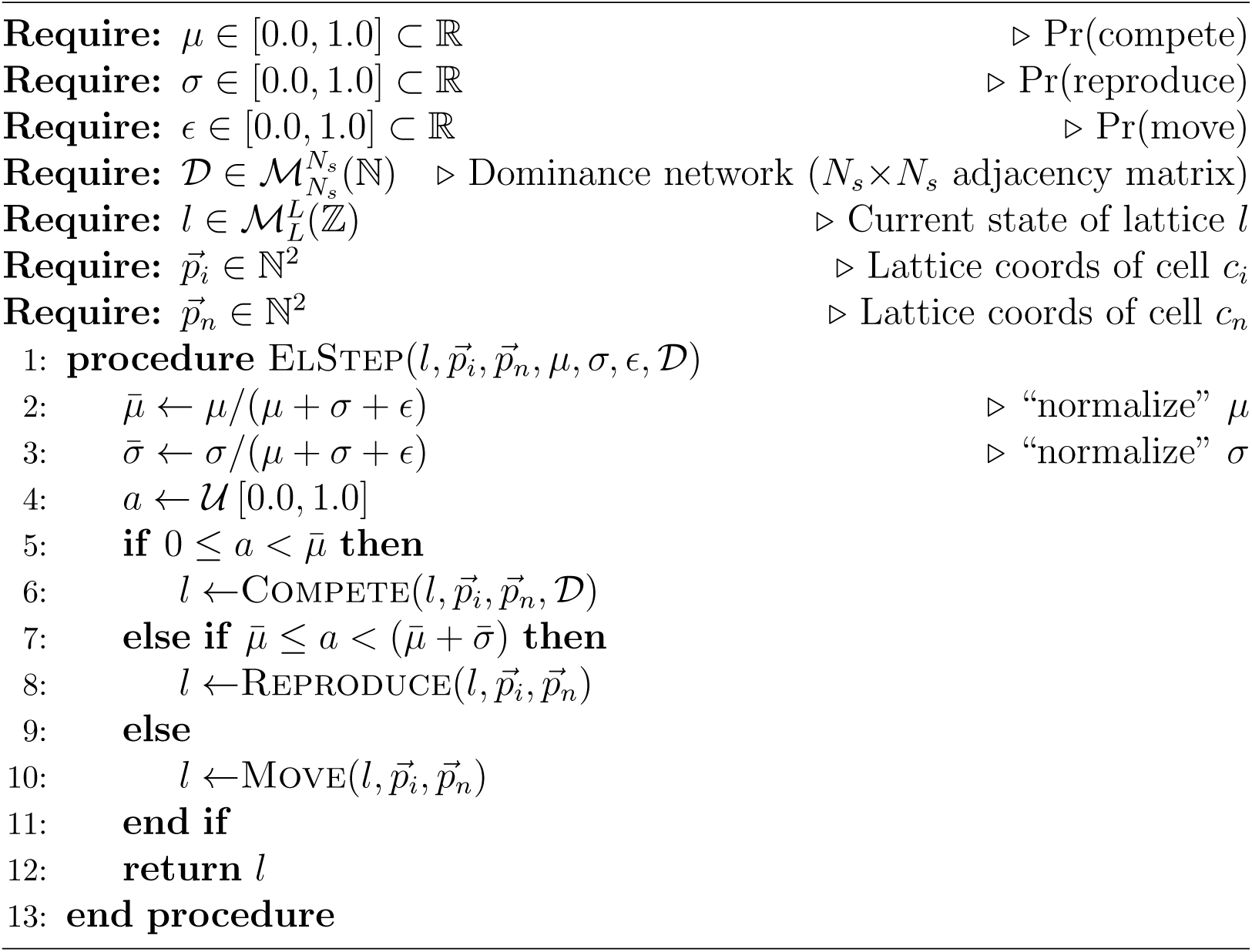
Original Elementary Step (OES)

In each individual Escg experiment, the density *ρ_i_*(*t*) of each species *S_i_* (i.e., what proportion of the lattice cells are occupied by agents of species type *S_i_*) was recorded after each mcs: illustrative time-series of the *ρ_i_* values from several experiments were given in [4]. For the purposes of this paper the primary variable of interest is *n_s_*(*t*) the number of non-extinct species at time *t* recorded over a set of *N_iid_* iid simulation experiments for any given set of hyperparameter values.

## 4. Replicating the results of Zhong et al. (2022)

The upper graph in each of Figures 4, 5, 6, and 7 re-prints^4^ the *FvsM* results from the *N_a_*=1, *N_a_*=2, *N_a_*=3, and *N_a_*=4 experiments presented in Figures 3, 5, 6, and 7 of Zhong et al. (2022), respectively, and the lower graph in each of Figures 4, 5, 6, and 7 shows corresponding *FvsM* results from my replication of each of those respective sets of experiments. For ease of comparison, my graphs use the same data-point markers and line-colors as Zhong et al., which (for reasons not stated by Zhong et al.) vary from graph to graph.

**Figure 4:**
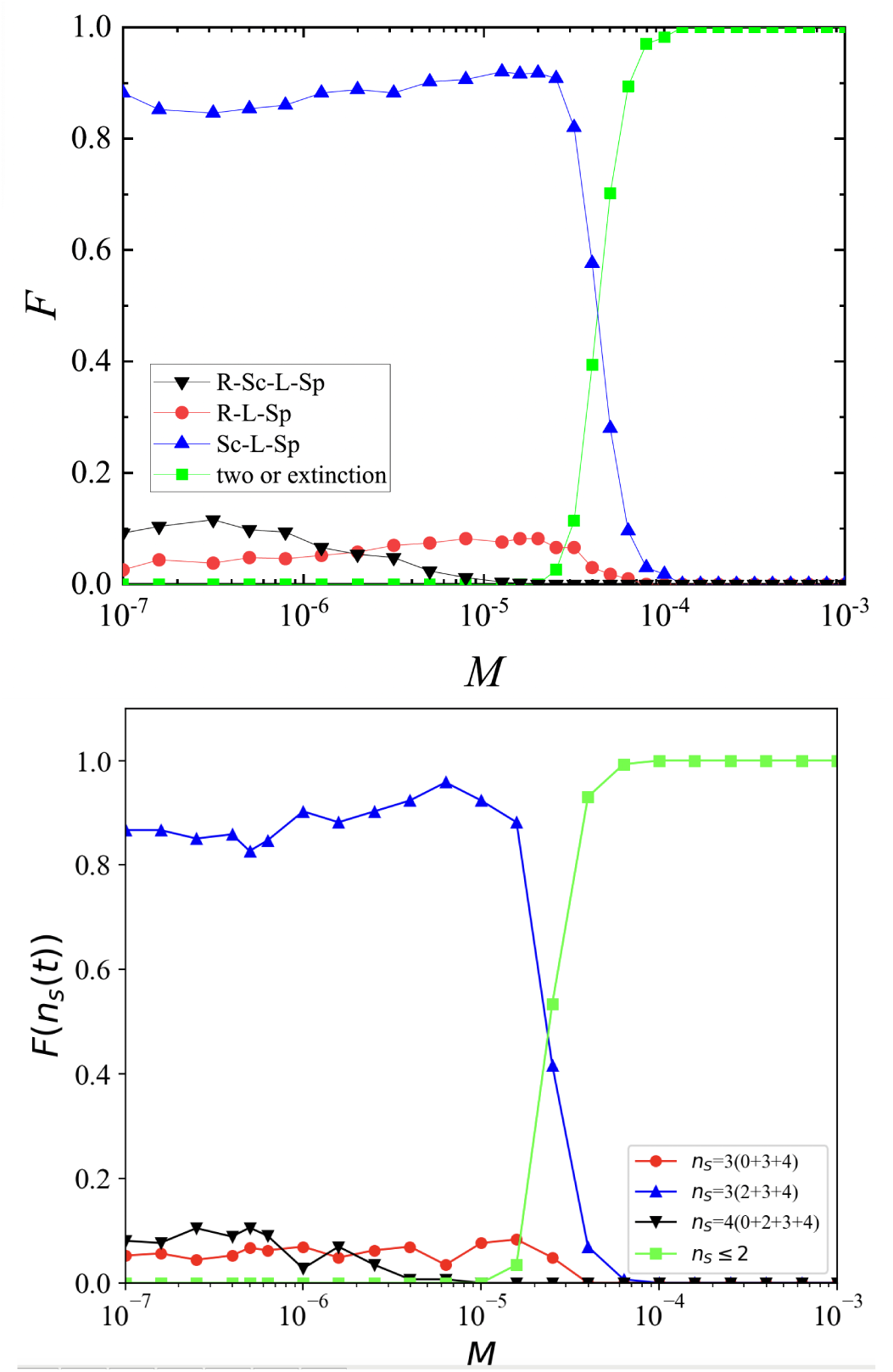
Graphs of *F* (*n_s_*(*t*_max_)=*c*) for *c* 1*, . . .,* 5 (i.e., frequency of outcome of number of species surviving at the end of the experiment) vs. *M* (mobility) – referred to as *FvsM* plots – in my replication of the *N_a_*=1 (dominance network Z1) experiments reported by Zhong et al.: upper graph is Zhong et al.’s Figure 3(a) *FvsM* results; lower graph shows corresponding *FvsM* results from my replication of the same experiments. The legend in Zhong et al.’s Figure 3(a) uses the abbreviations *R* for ‘Rock’, *Sc* for ‘Scissors’, *L* for ‘Lizard’, and *Sp* for ‘Spock’. Note that Zhong et al. combine the results for *F* (*n_s_*(*t*_max_)=2) and *F* (*n_s_*(*t*_max_)=1) into a single class of outcome labelled “two or extinction”. The legend in the lower graph of outcomes from my replication uses the numeric node-labels introduced in Figure 2: each row of the legend shows the color of the line and marker for the given value of *n_s_* followed by, for *F* (*n_s_*(*t*_max_)*>*2), in parentheses and separated by ‘+’ symbols, the node-numbers of that set of *n_s_* surviving species.

As can be seen, qualitatively there is extremely good agreement between the original results and my replication, but my results are not in exact quantitative agreement with the originals. Specifically, for all four of my *FvsM* graphs, I found that using *t*_max_=1.7 × 10^5^ (in contrast to the *t*_max_=10^5^ used by Zhong et al.) gave the best alignment between the two sets of results; and even with that adjustment to *t*_max_, the *M* -value at the crossover point where *F* (*n_s_*(*t*_max_) *>* 2) = *F* (*n_s_*(*t*_max_) ≤ 2) is slightly – but noticeably – lower in my results than in Zhong et al.’s: for example, in Figure 4 the crossover point in Zhong et al.’s *FvsM* graph is at *M* 4.1 10*^−^*^5^ whereas in my replication the crossover is at *M* 2.2 10*^−^*^5^.

These minor points of quantitative difference indicate that, in all likelihood, Zhong et al.’s actual implementation of their Rpsls Escg, their program code, does not exactly implement the Escg laid out in Algorithm 1. For instance, if Algorithm 1 is altered so that at line 9, where originally a cell is chosen at random from anywhere in the lattice, instead a random *non-empty* cell is chosen, per the amended algorithm snippet shown in Algorithm 4, then *every* call to ElStep will offer the possibility of either a competition, a reproduction, or a movement occurring, because we now guarantee that at least one of the two cells selected on each ES contains an individual, and hence the chances of a no-op on each ES are significantly reduced. This reduction in the number of no-ops per mcs would mean that the dynamics of the Escg would unfold at a quicker pace, because a greater number of substantive changes to the lattice would occur on each mcs, and so the value of *t*_max_ needed to get a good match with Zhong et al.’s results could be significantly reduced from the *t*_max_=1.7×10^5^ used here.

**Algorithm 4.**
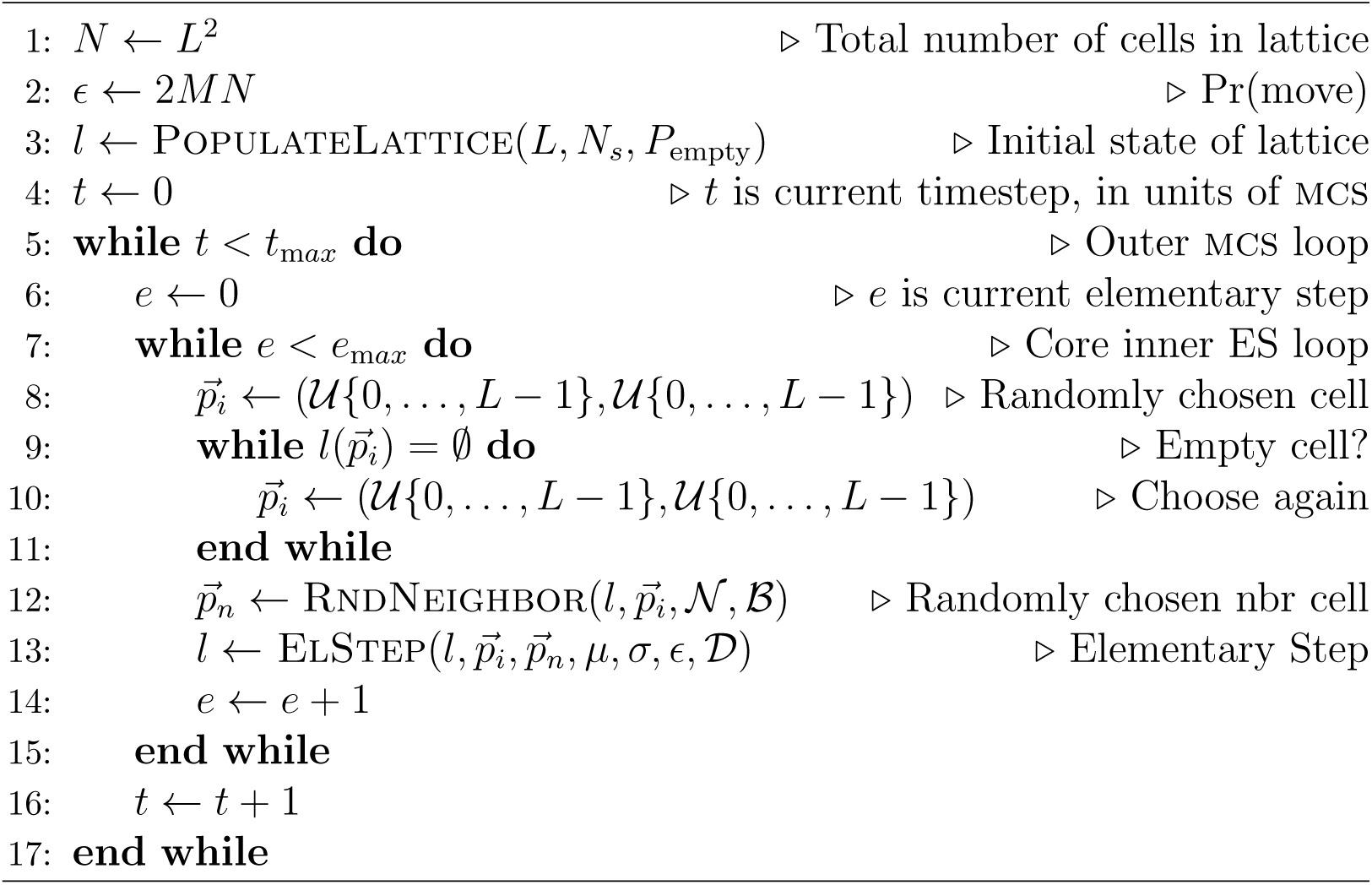
Amended Escg: start with non-empty cell

In principle, I could make a series of alterations to my implementation of Algorithm 1 such as the one just discussed, and then via a trial-and-error process eventually arrive at a better quantitative match between my replication results and those of Zhong et al., but to do so would not be a productive use of time because the arguments that follow in Sections 5 and 6 would be neither strengthened nor weakened by having the replication results fitting quantitatively closer to the original results: that is, the replication results as presented here are already sufficiently close to the originals to support the critique that I offer in the next two sections of this paper.

## 5. Asymptote or Transient?

In the abstract of their paper, in the captions to their figures 2 to 7, and repeatedly in the text of their paper, Zhong et al. state that they are presenting results showing asymptotic behaviors of the system, as they explain thus:

**Figure 5:**
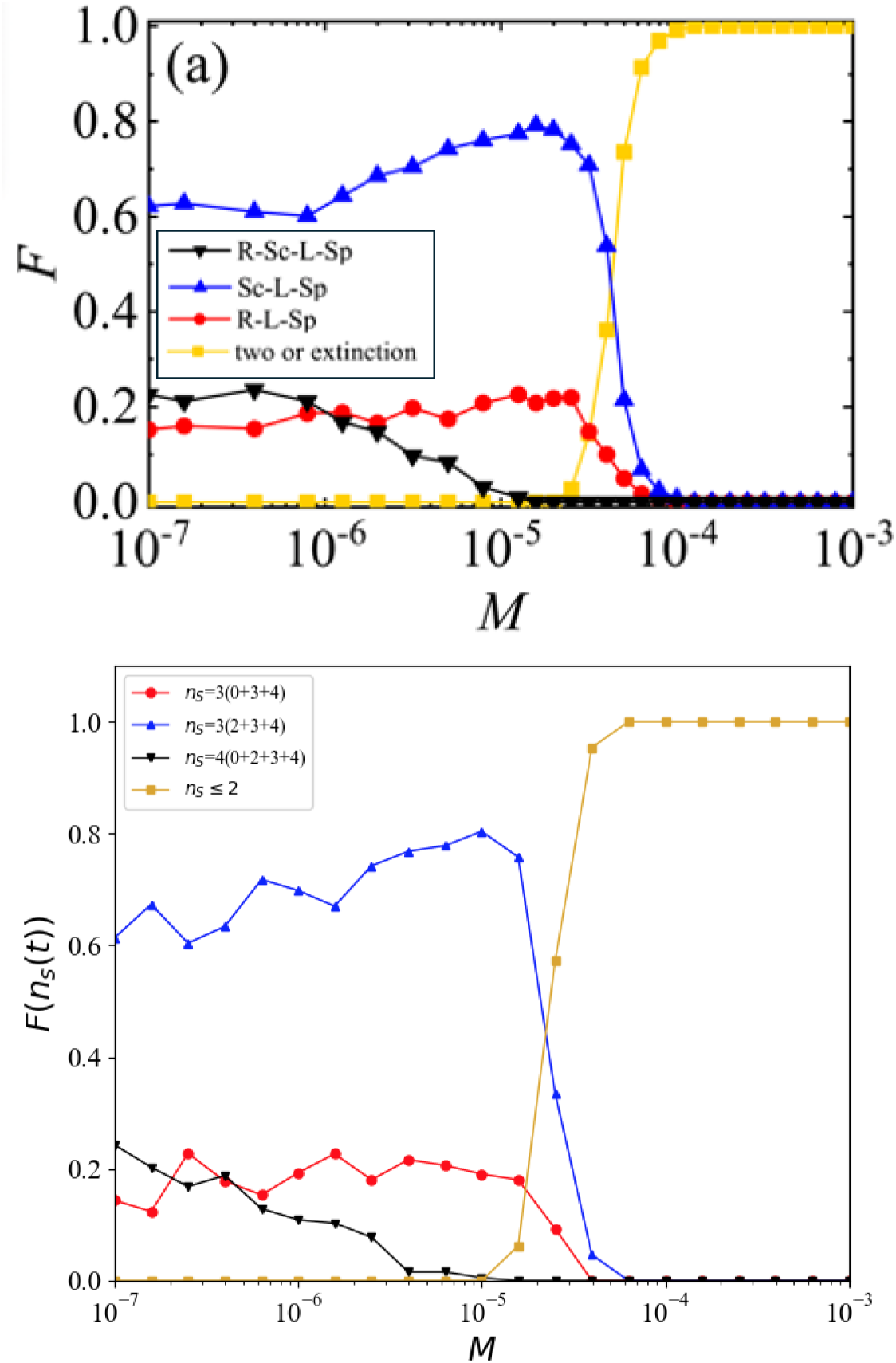
Replication of one of the two *N_a_*=2 (dominance network Z2a) experiments results reported by Zhong et al.: Upper graph is Zhong et al.’s Figure 5a (edited to include the legend); *N*_iid_=500; lower graph shows corresponding results from my replication of the same experiments; *N*_iid_=200. Format and legend labels as for Figure 4.

**Figure 6:**
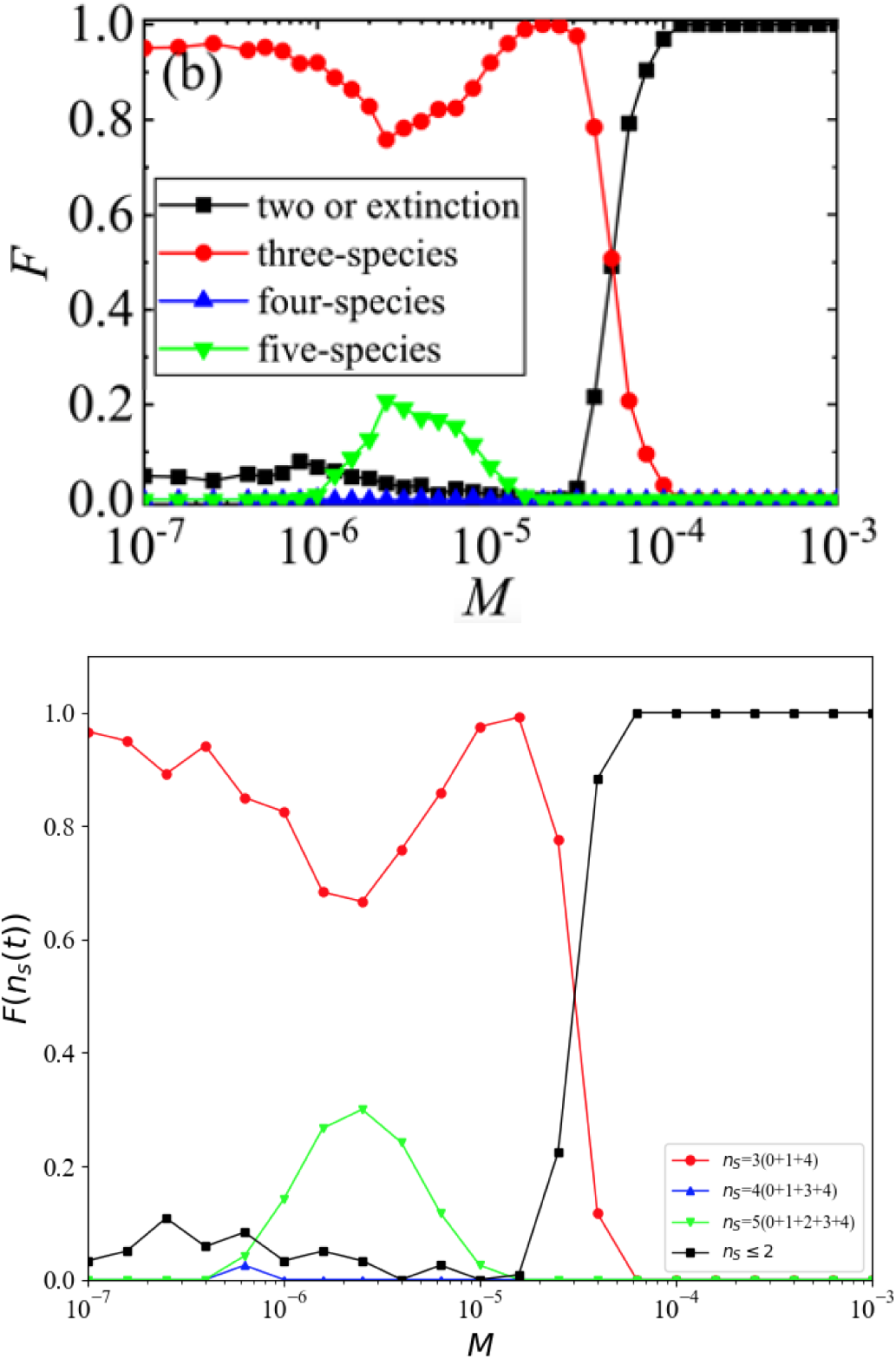
Replication of one of the two *N_a_*=3 (dominance network Z3b) experiments results reported by Zhong et al.: Upper graph is Zhong et al.’s Figure 6b (edited to include the legend); *N*_iid_=500; lower graph shows corresponding results from my replication of the same experiments; *N*_iid_=200. In Zhong et al’s Figure 6 and subsequent Figure 7 (shown here in Figure 7) the results are no longer explicitly presented for each distinct *n_s_*(*t*_max_)=*c* outcome such that, for example, the possible multiple different *n_s_*(*t*_max_)=3 outcomes illustrated in Figures 4 and 5 are instead presented in Zhong et al.’s Figures 6 and 7 as a single aggregate class of *n_s_*(*t*_max_)=3 outcome: for ease of comparison, I have followed that style of presentation here.

**Figure 7:**
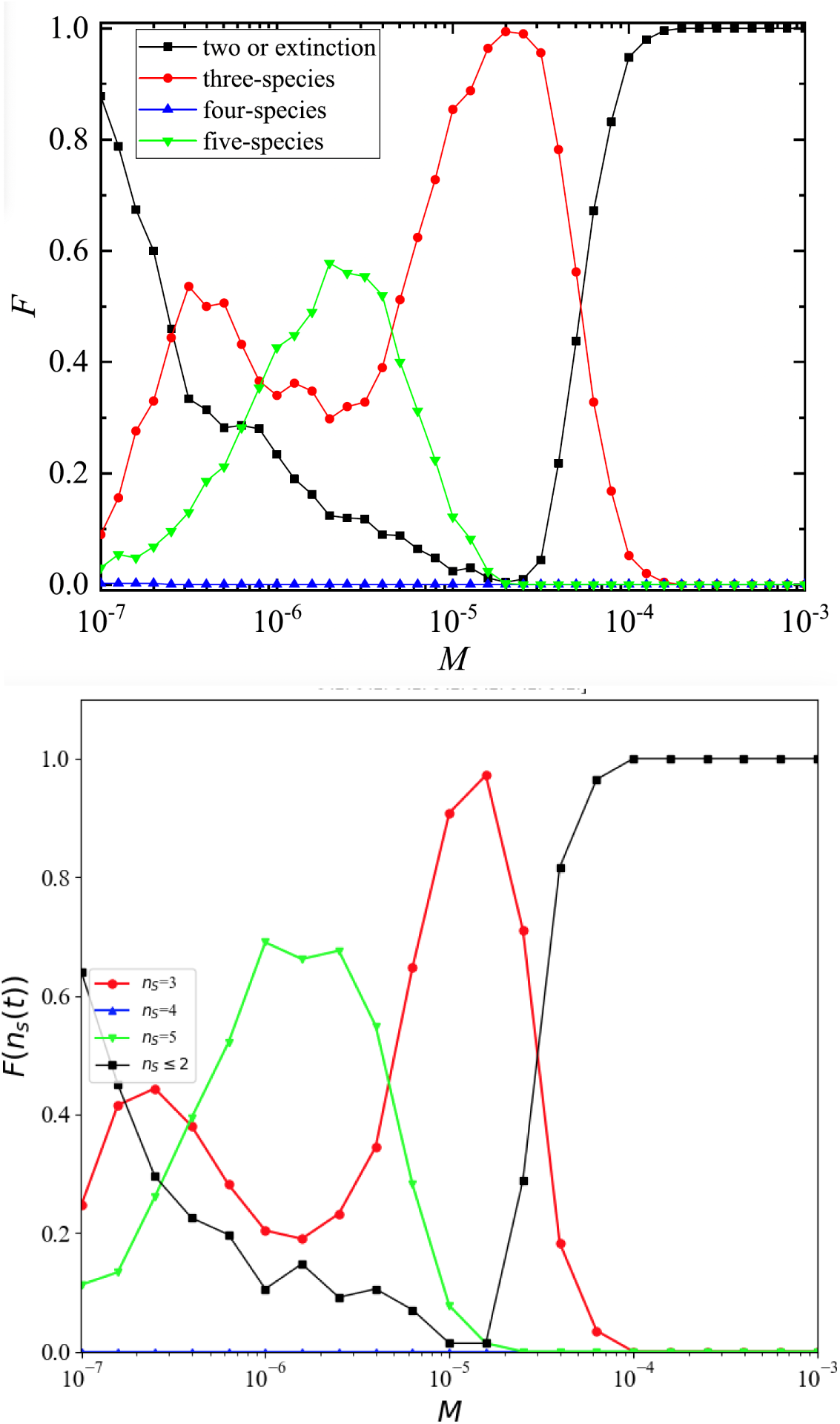
Replication of the *N_a_*=4 (dominance network Z4) experiments results reported by Zhong et al.: upper graph is Zhong et al.’s Figure 7, *N*_iid_=500; lower graph shows corresponding results from my replication of the same experiments; *N*_iid_=200. Format and legend labels as for Figure 4.

“To get an overview of the asymptotic behaviors against the mobility, we explore the occurrence of different asymptotic behaviors. For this aim, we run 500 realizations independently with equally and randomly distributed initial conditions for 10^5^ mcss and calculate the abundance of different asymptotic behaviors for each mobility.” [1, Section 3.1]

In common technical usage, “asymptotic behavior” refers to how some response or dependent variable behaves as the input or independent variable tends to infinity. Here, the independent variable is time (measured in mcs) and the dependent variable is the frequency of occurrence of the number of species coexisting in the modified Rpsls Escg system: for example, Zhong et al.’s caption to their Figure 2 states that two possible asymptotic behaviors are four-species coexistence and three-species coexistence, in experiments that ran for 10^5^mcs. Using the terminology introduced above in Section 3.1, Zhong et al. had *t*_max_=10^5^, and for each of their individual Escg simulation experiments the frequencies of observed outcome of each possible number of coexisting species at time *t*_max_, (denoted here by *n_s_*(10^5^)) was claimed to be the asymptotic behavior of that experiment. Implicit in this is the assumption that there is no need to run the simulations for longer than 10^5^mcs because there would be no further changes; that is, Zhong et al. assumed 10^5^mcs to be such a very large number of mcs as to be, for practical purposes, infinite. In this section I show that this assumption is incorrect, and that it can be necessary to run the simulations for *t*_max_ = 10^7^ or longer before the true asymptotic behaviors of the system can be convincingly identified. A consequence of this is that some of the features in their *FvsM* graphs that were highlighted for discussion by Zhong et al. are in fact only artefacts of slowly decaying transients in the system’s dynamics: when the Escg is run for sufficiently long, the features highlighted by Zhong et al. as asymptotic and worthy of discussion simply decay to nothing, and the diversity of observed outcomes reduces to near-homogeneity.

Figure 8 shows an exemplar set of results that illustrate this point: it shows FvsM plots for the Z2a ablated Rpsls dominance network at four different values of *t*_max_: 170kmcs (which gives a close match to Zhong et al.’s results for *t*_max_=10^5^ as was shown previously in Figure 5), 350kmcs, 500kmcs, and 10^6^mcs. Looking for instance at the *F* values at *M* =10*^−^*^7^, as the system is run for longer, the frequency of occurrence of four-species coexistence (i.e., *F* (*n_s_*(*t*_max_)=4)) falls from roughly 20% at *t*_max_=170k to absolute zero at *t*_max_=10^6^, and the *t*_max_=10^6^ results show conclusively that four-species co-existence is in fact not an asymptotic behavior of this system at all: the presence of four-species outcomes in Zhong et al.’s Z2a results is merely an artefact of their experiments having not been run for long enough.

**Figure 8:**
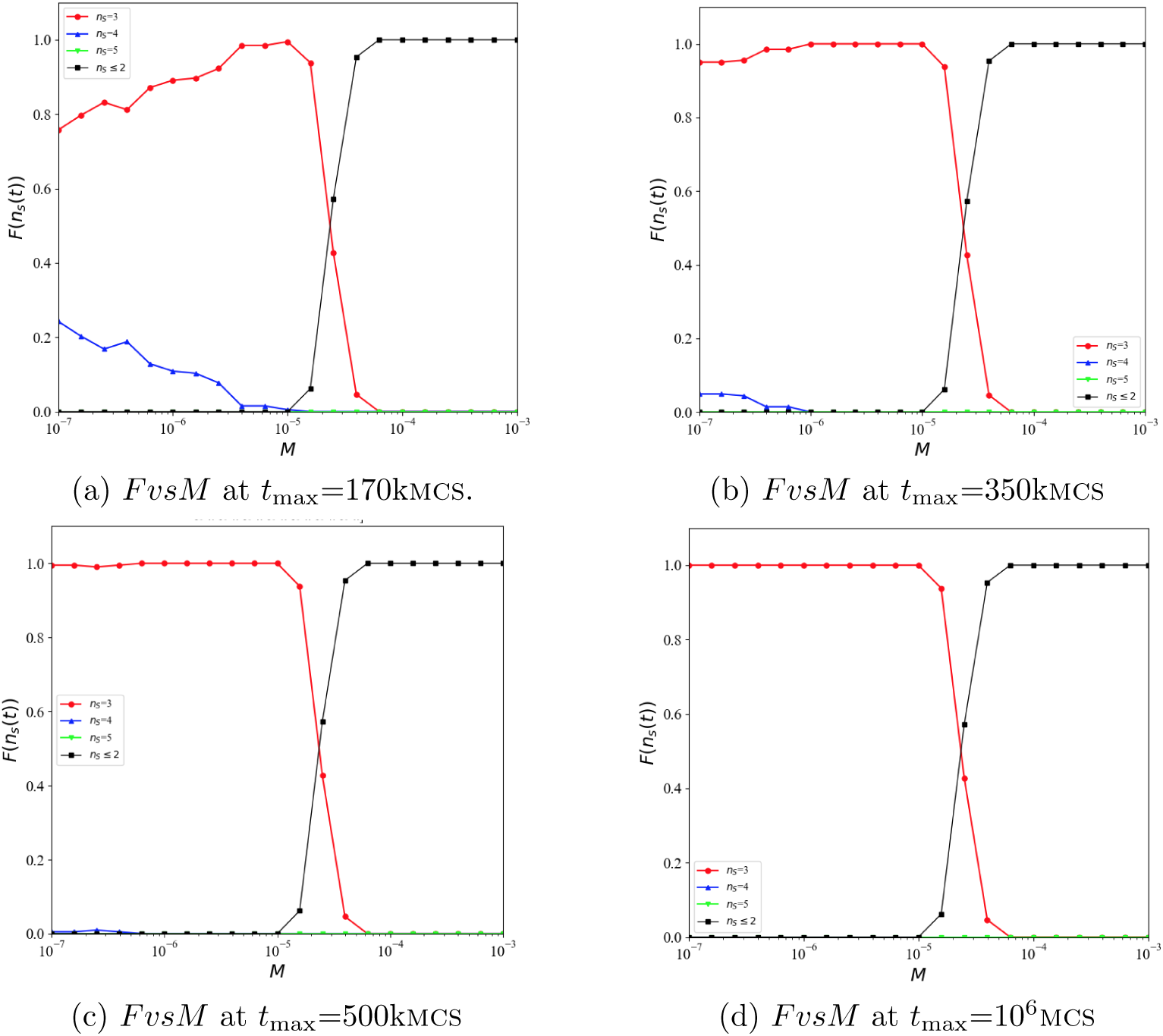
Plots of *FvsM* for the *N_a_*=2 Rpsls dominance network Z2a, *N* =200×200, results from which were shown in Zhong et al.’s Figure 5a; for four different experiment durations: (a) *t*_max_=1.7×10^5^mcs (which gives the best match to Zhong et al.’s Figure 5a); (a) *t*_max_=3.5×10^5^mcs; (c) *t*_max_=5.0×10^5^mcs; (d) *t*_max_=10^6^mcs. Format and legend labels as for Figure 6. In each graph, at each value of *M*, results from *N*_iid_=200 simulations are shown. As can be seen, as *t*_max_ is increased, F(*n_s_*(*t*_max_)=4)→0, and the true asymptotic frequency distribution of outcomes is shown in (d): for *M* below some threshold value of ≈2.5×10*^−^*^4^ the outcome is always *n_s_*(*t*)=3, and once *M* is above the threshold the outcome is always *n_s_*(*t*)≤2.

To get a better sense of the nature of these transients, data such as that shown in the four graphs in Figure 8 (which plotted *F* (*n_s_*(*t*_max_)) vs. *M* at different specific values of *t*_max_) can be visualised instead along the orthogonal axis of projection, i.e. plotting *F* (*n_s_*(*t*)) vs. *t* at different specific values of *M* : for brevity, I will refer to these as *FvsT* plots. Figure 9 shows one such graph of *FvsT* from *N*_iid_=500 simulation experiments with the *N_a_*=2 Rpsls dominance network Z2a, for *t*_max_=10^6^ and with *M* =10*^−^*^7^. As would be expected, initially all experiments have five species co-existing, so *F* (*n_s_*(*t*)=5)=1.0 and *F* (*n_s_*(*t*)=*c*)=0.0 for all other values of *c*; then at *t*≈150mcs, the simula-tions start to show their first extinction, and *F* (*n_s_*(*t*)=5) drops rapidly to zero, matched by the consequent rise in *F* (*n_s_*(*t*)=4) which reaches 1.0 at *t* 1kmcs; then, commencing at *t* 1kmcs, we see a growing number of experiments undergoing a second extinction, leaving three co-existing species, and hence *F* (*n_s_*(*t*)=3) starts to rise toward 1.0 while *F* (*n_s_*(*t*)=4) attenuates, eventually falling to zero at *t*≈700kmcs.

**Figure 9:**
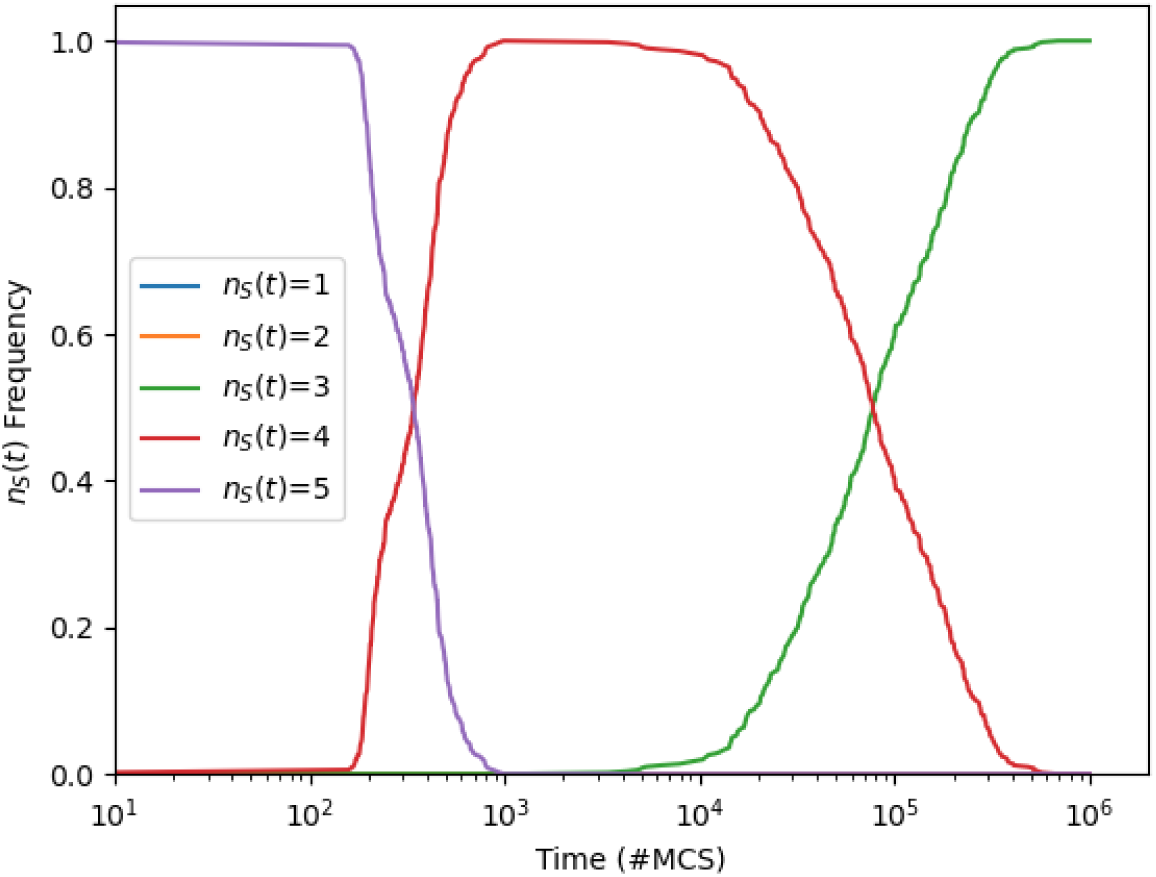
Time series of *F* (*n_s_*(*t*)=*c*) for *c* ∈ {1*, . . .,* 5} (i.e., frequency of occurrence of the number of co-existing species at time *t*) from *N*_iid_=500 simulation experiments using the *N_a_*=2 Rpsls dominance network Z2a, with *t*_max_=10^6^ and with *M* =10*^−^*^7^. For brevity, plots such as these are referred to as *FvsT* plots. Horizontal axis is time measured in mcs; vertical axis is *F* (*n_s_*(*t*)=*c*).

The fact that the true asymptotic state for this system is reached at *t* ≈ 700kmcs demonstrates that Zhong et al.’s assertion that running simulations of this system for *t*_max_=10^5^mcs is sufficient to identify its asymptotic behaviors is simply incorrect, and simulations substantially longer than 100kmcs are necessary.

Thus far the discussion here has concentrated on results from the Z2a ablated dominance network, but the same issue with slowly decaying transients being mis-identified as asymptotic outcomes affects the results presented by Zhong et al. for networks Z1 and Z3b: Figure 10 shows for comparison the *FvsM* plots for Z1 at *t*_max_ = 170*k* and *t*_max_=10^6^; Figure 11 shows *FvsM* for Z3b at the same two values of *t*_max_; and Figure 12 shows example *FvsT* plots for networks Z1 and Z3b, illustrating the slowly decaying transients.

**Figure 10:**
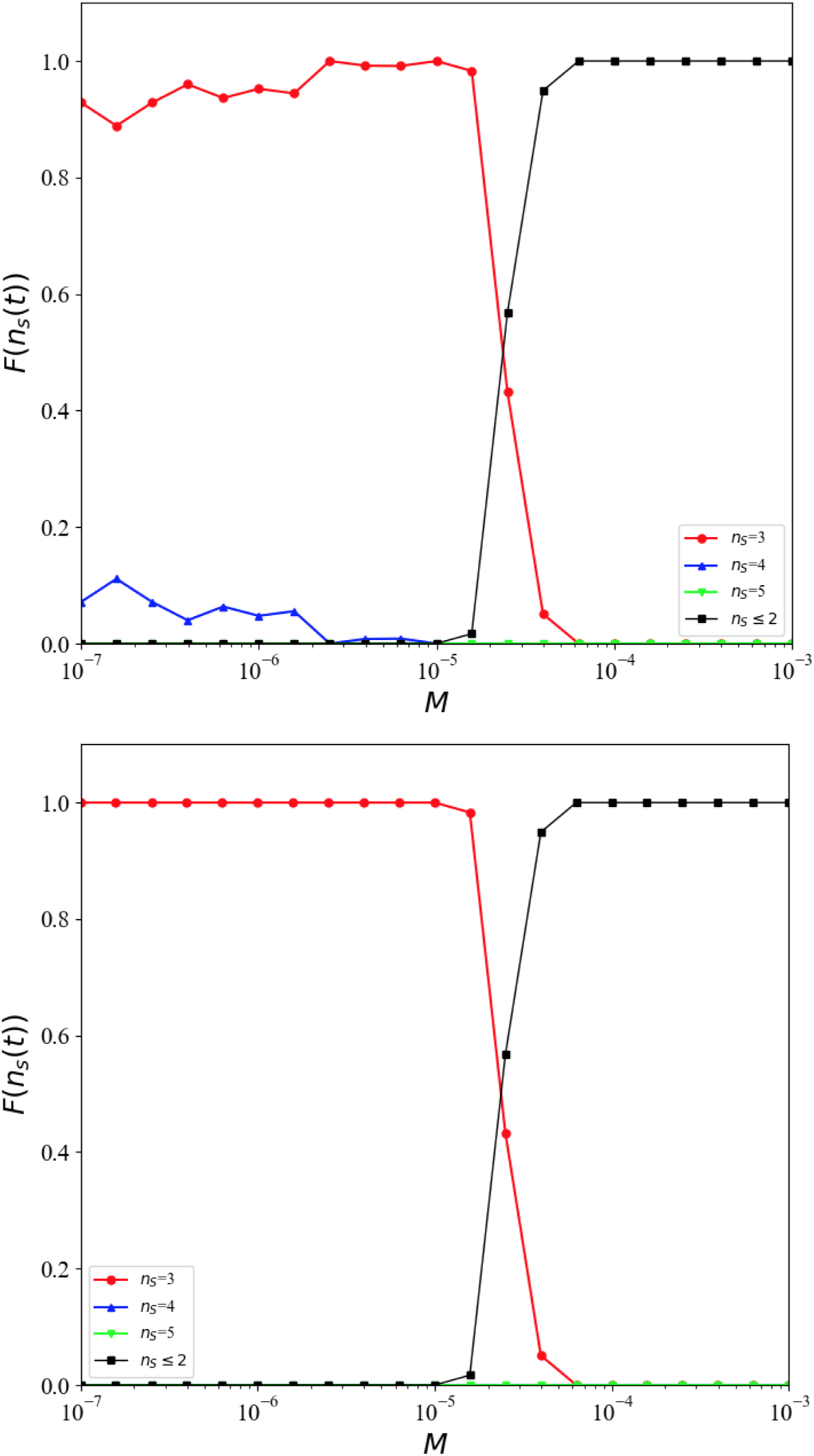
Plots of *FvsM* for the Z1 ablated dominance network at (upper graph) *t*_max_=170*k*mcs and (lower graph) *t*_max_=10^6^mcs. Format and legend labels as for Figure 6.

**Figure 11:**
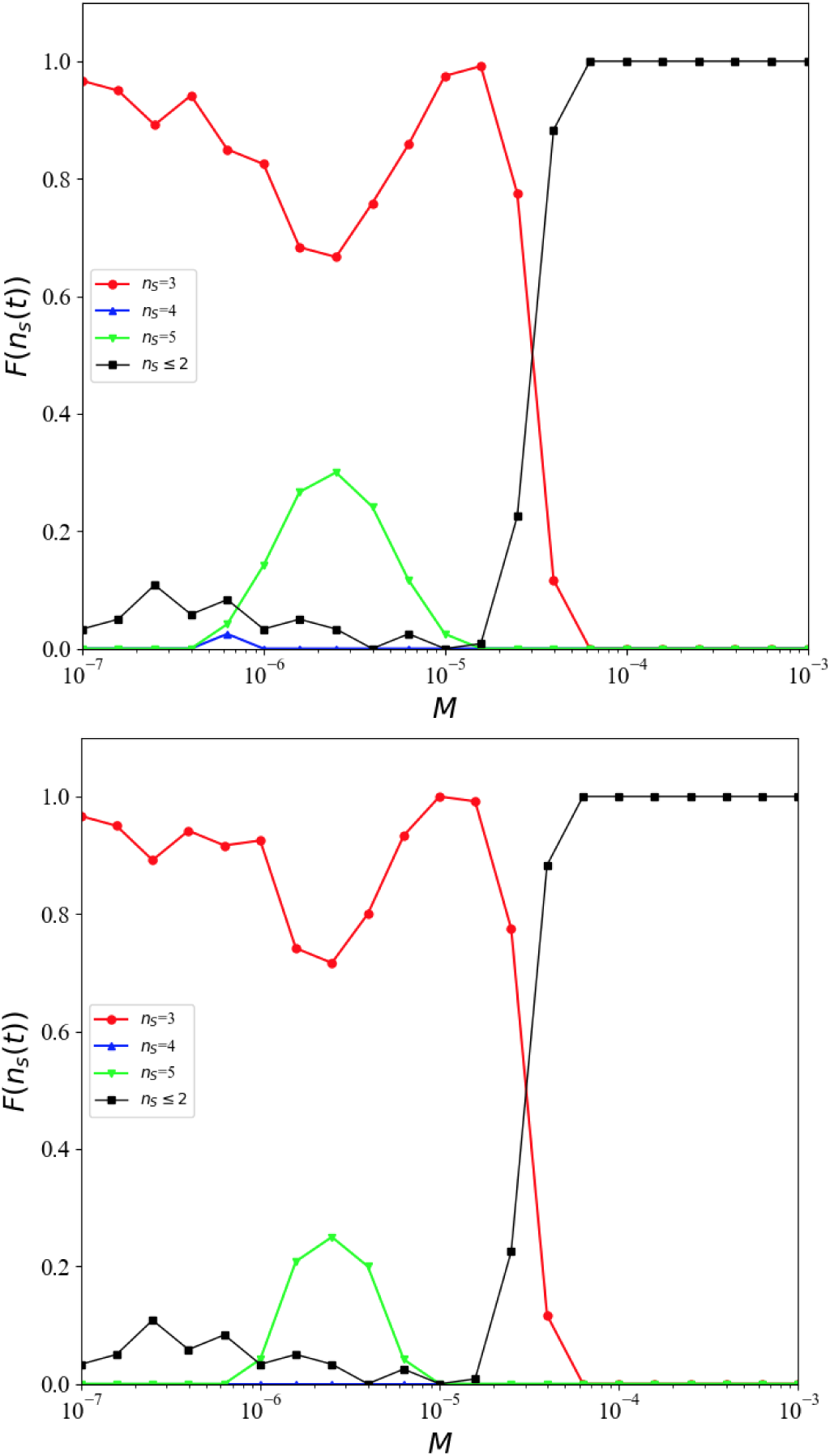
Plots of *FvsM* for the *N_a_*=3 Rpsls dominance network Z3b, results from which were shown in Zhong et al.’s Figure 6b; *N* =200×200, for two different durations of experiment. Upper graph is from experiment duration *t*_max_=1.7 10^5^mcs, which gives the best match to the Z3b results published by Zhong et al. (2022); lower graph is from experiment duration *t*_max_=10^6^mcs. Format and legend labels as for Figure 4.

**Figure 12:**
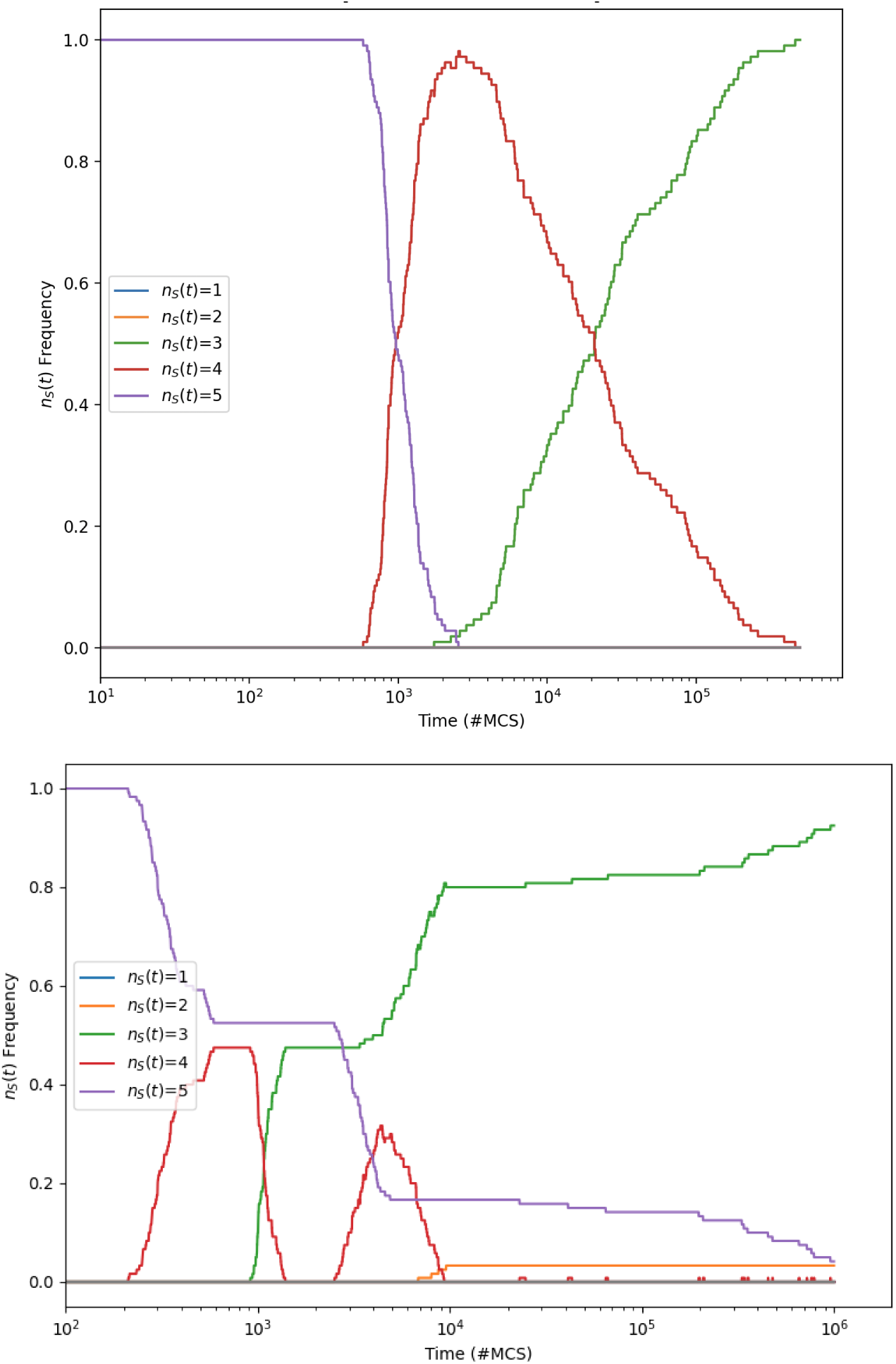
Illustrative *FvsT* plots at *t*_max_=10^6^ for the Z1 ablated dominance network at *M* =10*^−^*^7^ (upper graph) and for the Z3b ablated dominance network at *M* =10*^−^*^6^ (lower graph); format as for Figure 9. For Z1, the longest-persisting transient decays to zero by *t* 5 10^5^mcs. For Z3b, even at *t*_max_=10^6^, the *F* (*n_s_*(*t*)=5) transient is clearly decaying but has not yet reached zero; further *FvsT* plots for Z3b are presented in Appendix B.

The *FvsM* plots for ablated dominance network Z1 in Figure 10 and the *FvsT* plot for Z1 in Figure 12 demonstrate conclusively that four-species coexistence is not an asymptotic behavior for this system. However the *FvsM* plots for ablated dominance network Z3b in Figure 11 and the *FvsT* plot for Z3b in Figure 12 are not so immediately conclusive. Writing about their Z3b *FvsM* plot, Zhong et al. observe:

“At low mobility, the state of Scissors-Lizard-Spock dominates the population, its occurrence stays at high value except for a dip at around *M* =3 × 10*^−^*^6^. Different from other interaction structures in modified Rpsls, it may allow for five-species coexistence since the two three-species-cyclic interactions only share one species, Lizard. Once the coexistence of Rock-Lizard-Spock becomes possible, the coexistence of five species becomes possible. Actually, Fig. 6(b) [*reproduced here as the upper graph in* Figure 6] shows that the occurrence of asymptotic five-species state displays a bump at around *M* =3×10*^−^*^6^.” [1, Section 3.3]

In Figure 11 the ‘bump’ in frequency of five-species coexistence visible in the low-*t*_max_ *FvsM* data for *M* ∈ [≈4×10*^−^*^7^, ≈2×10*^−^*^5^] is still present in the high-*t*_max_ *FvsM* data, although the longer run-time has resulted in minor reductions in the bump’s height and width. Clearly some further simulation work is required to make a convincing case for what are the true asymptotic behaviors of the system with the Z3b dominance network.

To that end, Figure 13 shows an *FvsM* plot of outcomes from a set of Z3b experiments where *t*_max_=10^7^, ten times longer than the *t*_max_=10^6^ Z3b results shown in Figure 11, and 100 times longer than the *t*_max_=10^5^ used by Zhong et al. in their experiments. As can be seen, when run to *t*_max_=10^7^, the ‘bump’ of five-species coexistence is almost completely flattened to zero. The slowly decaying transient of five-species coexistence, which causes that bump in the *FvsM* plot when the experiments are not run for long enough, is further illustrated in the *FvsT* plots of Figure 14. However, for the bump to be completely eliminated, and the true asymptotic behavior of the system to be visible in the *FvsM* plot, clearly the simulation would need to be run even longer than *t*_max_=10^7^: looking at the rate of decay for *F* (*n_s_*(*t*)=5) in the lower graph of Figure 14, it seems likely to hit zero by *t*_max_≈10^8^. Given that the Z3b *F* (*n_s_*(*t*)=5) decays to zero for all other values of *M*, I think it intuitively reasonable to conjecture – to make a projection or prediction – that the true asymptotic behavior Z3b at *M* 2.5 10*^−^*^6^ is *F* (*n_s_*(*t*)=5)=0, which gives the *FvsM* plot for the true asymptotic behavior of the Z3b system that is shown in Figure 15.

**Figure 13:**
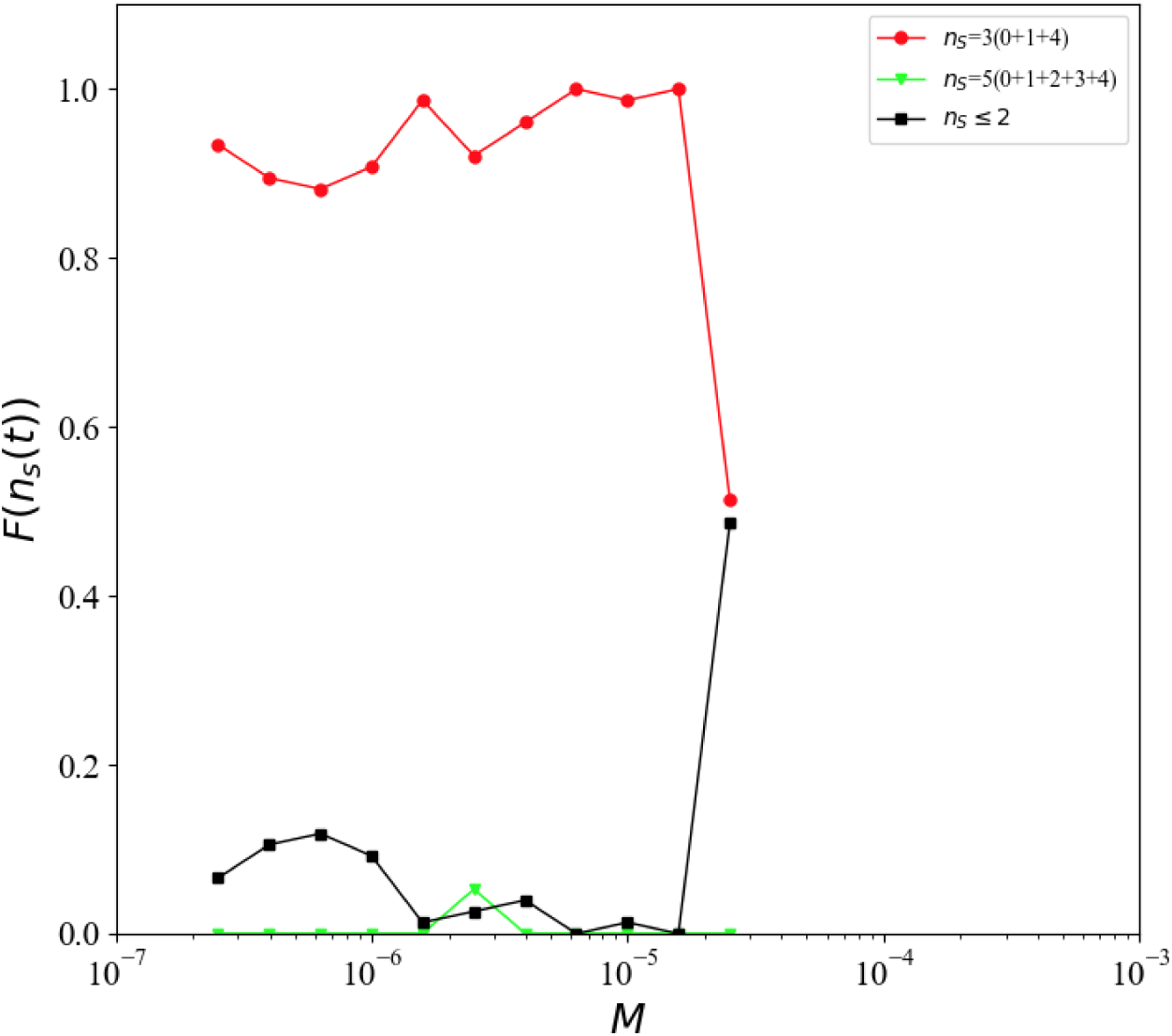
Plot of *FvsM* for the Z3b ablated dominance network run to *t*_max_=10^7^, one hundred times longer than that used by Zhong et al., data generated only over the range *M* ∈ [2.51 10*^−^*^7^, 2.51 10*^−^*^5^,], the values for which the *FvsM* plots from shorter-running experiments showed a ‘bump’ of five-species co-existence. As can be seen, the bump has almost entirely disappeared as a consequence of increasing the run-time to *t*_max_=10^7^. Format and legend labels as for Figure 4.

**Figure 14:**
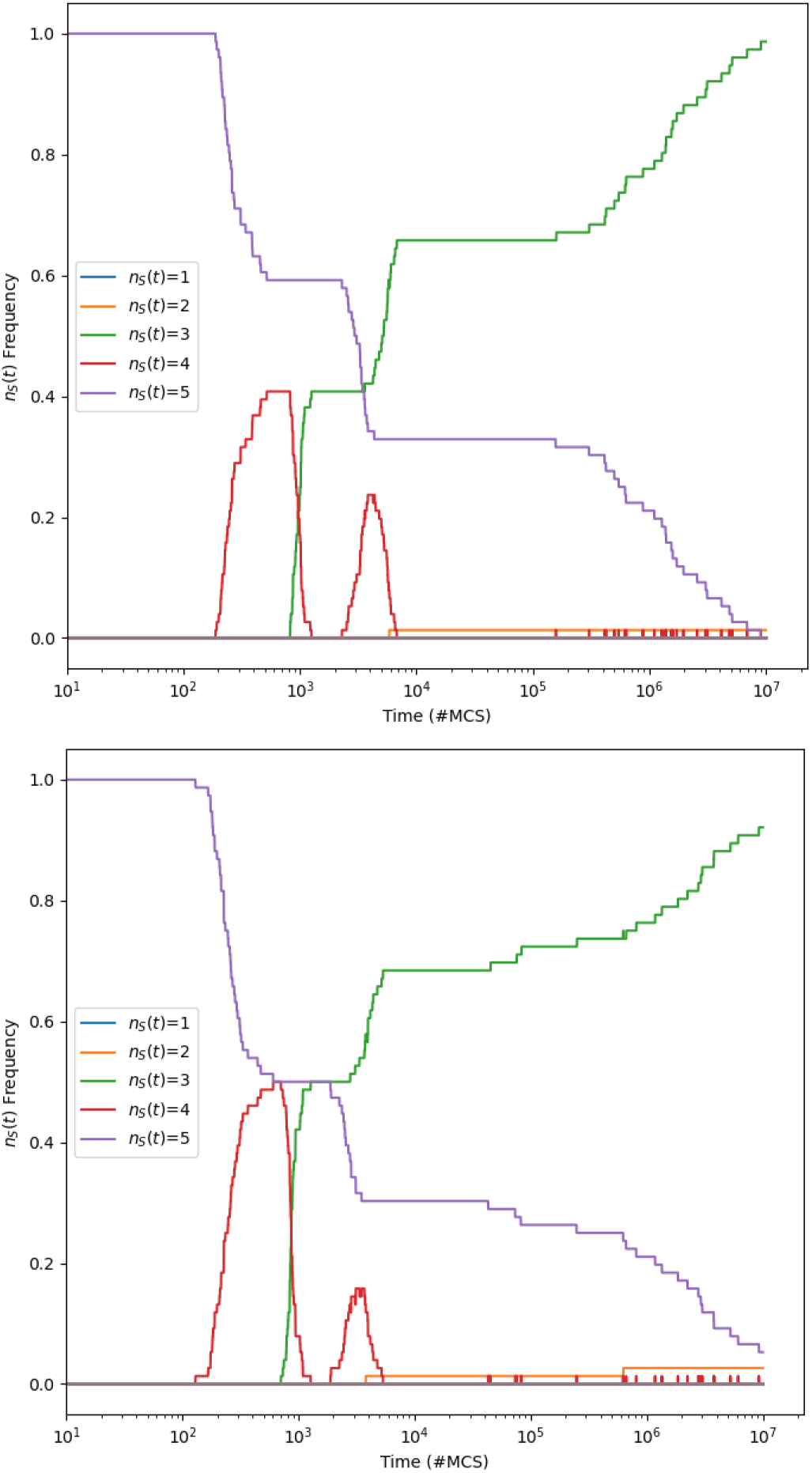
Plots of *FvsT* for the Z3b ablated dominance network run to *t*_max_=10^7^, one hundred times longer than that used by Zhong et al.: upper graph is for *M* =1.58 10*^−^*^6^; lower graph is for *M* =2.51 10*^−^*^6^. In both graphs, at *t*=10^5^mcs (the value of *t*_max_ used by Zhong et al. to determine the “asymptotic behavior” of the system) the frequency of five-species coexistence *F* (*n_s_*(*t*)=5)*>*20% but when the experiment is continued to *t*_max_=10^7^, the value of *F* (*n_s_*(*t*)=5) steadily decays toward zero, and it seems reasonable to conjecture that zero is its true asymptotic value. Format as for Figure 9.

**Figure 15:**
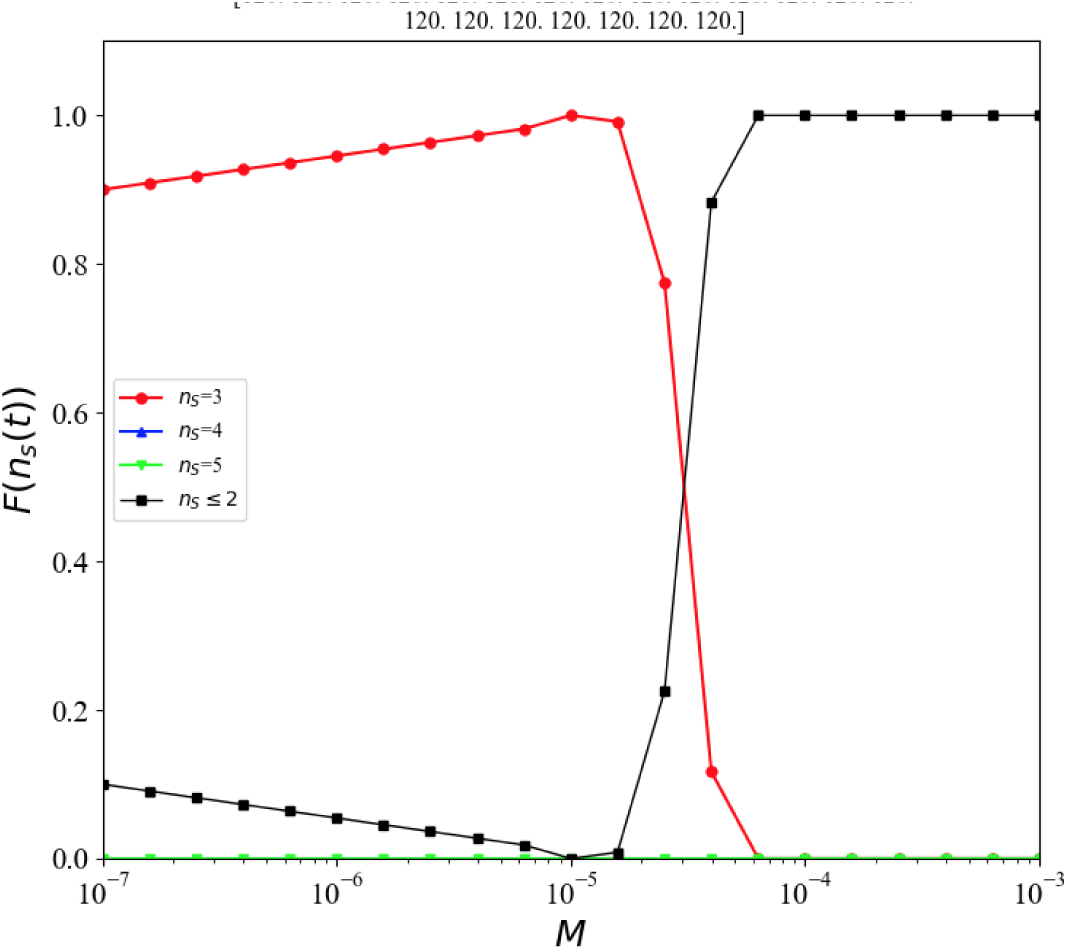
Plot of predicted/projected *FvsM* for the true asymptotic behavior of the Z3b ablated dominance network, based on the data previously shown in Figures 11 and 13, and with *F* (*n_s_*(*t*)=5) predicted to be zero at *M* =2.51 10*^−^*^6^ for the reasons given in the text. The *F* values shown for *M* ≥ 10*^−^*^5^ are results from simulations lasting *t*_max_=10^6^ (from Figure 11); the *F* values for 10*^−^*^7^ ≤ *M<*10*^−^*^5^ are synthetic data showing outcomes predicted on the basis of Figure 13 and the arguments given in the text.

To summarise the critique thus far: the true asymptotic *FvsM* plots for Z1 (shown in Figure 10), Z2a (Figure 8d), and Z3b (Figure 15), are all visualizations of essentially the same outcome, and are also equally similar to the *FvsM* results for ablated dominance networks Z2b and Z3a shown in Zhong et al.’s Figures 5b and 6a. That is, for each of the five ablated Rpsls systems Z1 to Z3b studied by Zhong et al., when the system is run with a sufficiently large value of *t*_max_, the true asymptotic behavior is essentially identical across all five cases. The only thing that varies is how many time-steps the simulations of the differently ablated networks have to be run for, for the system to converge on its asymptotic behavior.

In the next section, I go on to show that the true asymptotic behavior of the original Rpsls system with the *unablated* dominance network Z0 is *also* essentially identical to the asymptotic behaviors of the five ablated Rpsls systems Z1 to Z3b.

## 6. Baseline: dynamics of unablated Rpsls at *N* =200*×*200

Figure 16 shows *FvsM* plots for the unablated (*N_a_*=0) original Rpsls system with dominance network Z0 (illustrated in Figure 2) at *t*_max_=10^5^ and *t*_max_=10^6^. As can be seen, the results from *t*_max_=10^5^ are far from asymptotic, given the degree of change brought about by extending the run-time to *t*_max_=10^6^. The Z0 *FvsM* plot for *t*_max_=10^6^ is similar to those for Z1, Z2a, and Z3 (shown in Figures 10, 8d, and 15, respectively), except that in the range *M* ∈[10*^−^*^4^, 10*^−^*^3^] there is another ‘bump”, this time in the *F* (*n_s_*(*t*)=3) values, similar to the bump in *F* (*n_s_*(*t*)=5) values highlighted by Zhong et al. in their Z3b results, as discussed in the previous section. And, as with the Z3b results, the presence of this bump in the Z0 *FvsM* plot is an indication that running experiments to *t*_max_=10^6^ is too short for the true asymptotic behavior of this system to be shown.

**Figure 16:**
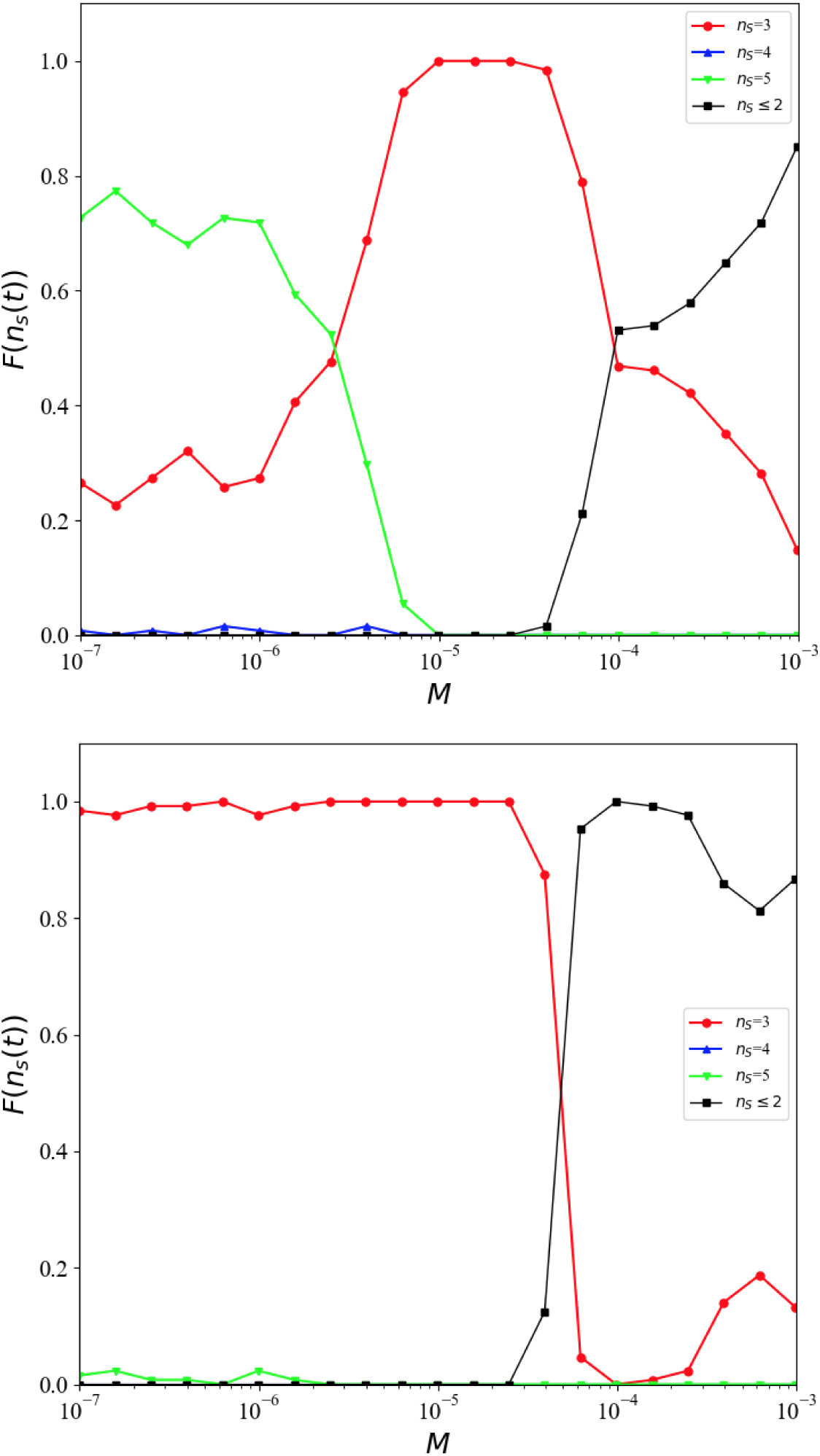
Plots of *FvsM* for the unablated (*N_a_*=0) Rpsls dominance network Z0 (illustrated in Figure 2) with *N* =200×200, for two different durations of experiment. Upper graph is from experiment duration *t*_max_=10^5^mcs as used by Zhong et al. (2022); lower graph is from experiment duration *t*_max_=10^6^mcs. Format and legend labels as for Figure 4.

To address that, the Z0 experiments were extended to *t*_max_=10^7^ for the range of *M* values where the bump in *F* (*n_s_*(*t*)=3)) values is visible in the *t*_max_=10^6^ plot of *FvsM* plot, and the results from these longer experiments are shown in Figure 17: with the longer run-time, the *FvsM* bump in the *F* (*n_s_*(*t*)= 3) data over range *M* ∈ [10*^−^*^4^, 10*^−^*^3^] has almost entirely disappeared, remaining visible here only for *M* 6.3 10*^−^*^4^. Plots of *FvsT* at *t* ≥ 10^7^ for *M* =6.3 10*^−^*^4^ and for *M* =10*^−^*^3^ are shown in Figure 18: given that for lower values of *M* the *F* (*n_s_*(*t*)=3) values decay to zero by *t*=10^7^, it seems intuitively obvious that these two long-lasting transients in *F* (*n_s_*(*t*)=3) are both monotonically approaching zero once *t>*10^5^ (with the consequence that *F* (*n_s_*(*t*)=1)→1.0) and would eventually reach zero if the durations of the experiments were sufficiently extended. That is, a reasonable projection to make here is that the true asymptotic behavior for the Z0 system with *M* ≥6.3 is *F* (*n_s_*(*t*)=3))=0 and *F* (*n_s_*(*t*)=1))=1: this projection is illustrated in the summary Z0 asymptotic *FvsM* plot of Figure 19.

**Figure 17:**
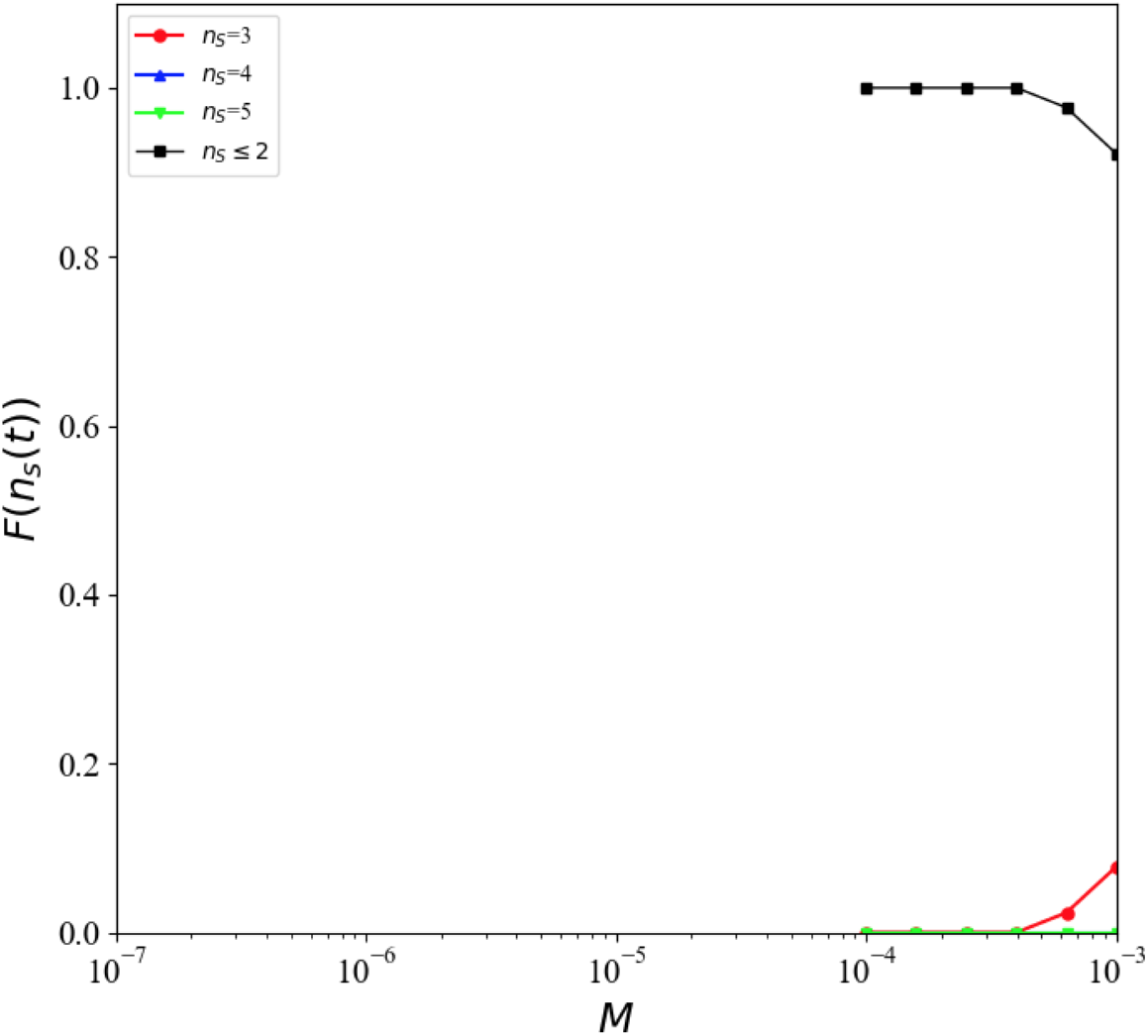
Plot of *FvsM* at *t*_max_=10^7^mcs for the unablated (*N_a_*=0) Rpsls dominance network Z0 (illustrated in Figure 2) with *N* =200 200, over the range of *M* values where the ‘bump’ in *F* (*n_s_*(*t*)=3) occurred in the *FvsM* plot for *t*_max_=10^6^mcs shown in Figure 16: here, with a tenfold longer run-time, the bump in *F* (*n_s_*(*t*)=3) has almost entirely disappeared, remaining nonzero in this plot only at *M>* 6 10*^−^*^4^. Format and legend labels as for Figure 4.

**Figure 18:**
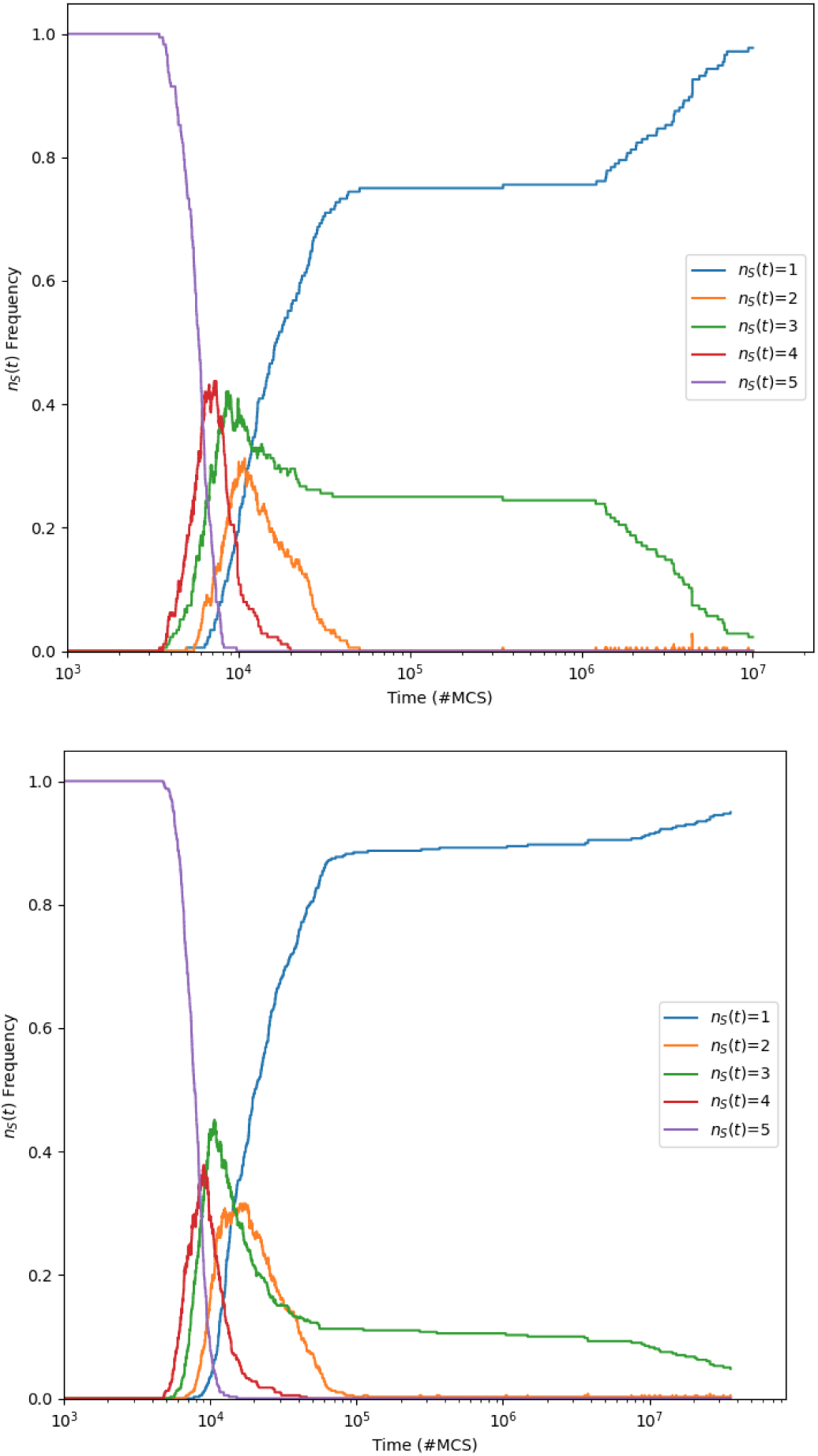
Plots of *FvsT* to *t*_max_ ≥ 10^7^ for the unablated (*N_a_*=0) original Rpsls dominance network Z0 (illustrated in Figure 2) with *N* =200×200: upper graph shows results to *t*=10^7^ for *M* =6.3 10*^−^*^4^ from *N*_iid_=175 simulations, where *F* (*n_s_*(*t*)=3) holds almost constant at ≈25% for 4 10^4^ *t* 10^6^, but which then falls steadily toward zero; lower graph shows results to *t*=3.5 × 10^7^ for *M* =10*^−^*^3^ from *N*_iid_=400 simulations, where the rate of decay of *F* (*n_s_*(*t*)=3) is shallow over 10^5^ *t* 8 10^6^ but then steepens for *t>*8 × 10^6^. Further illustrative *FvsT* plots for Z0 are given in Appendix A.

**Figure 19:**
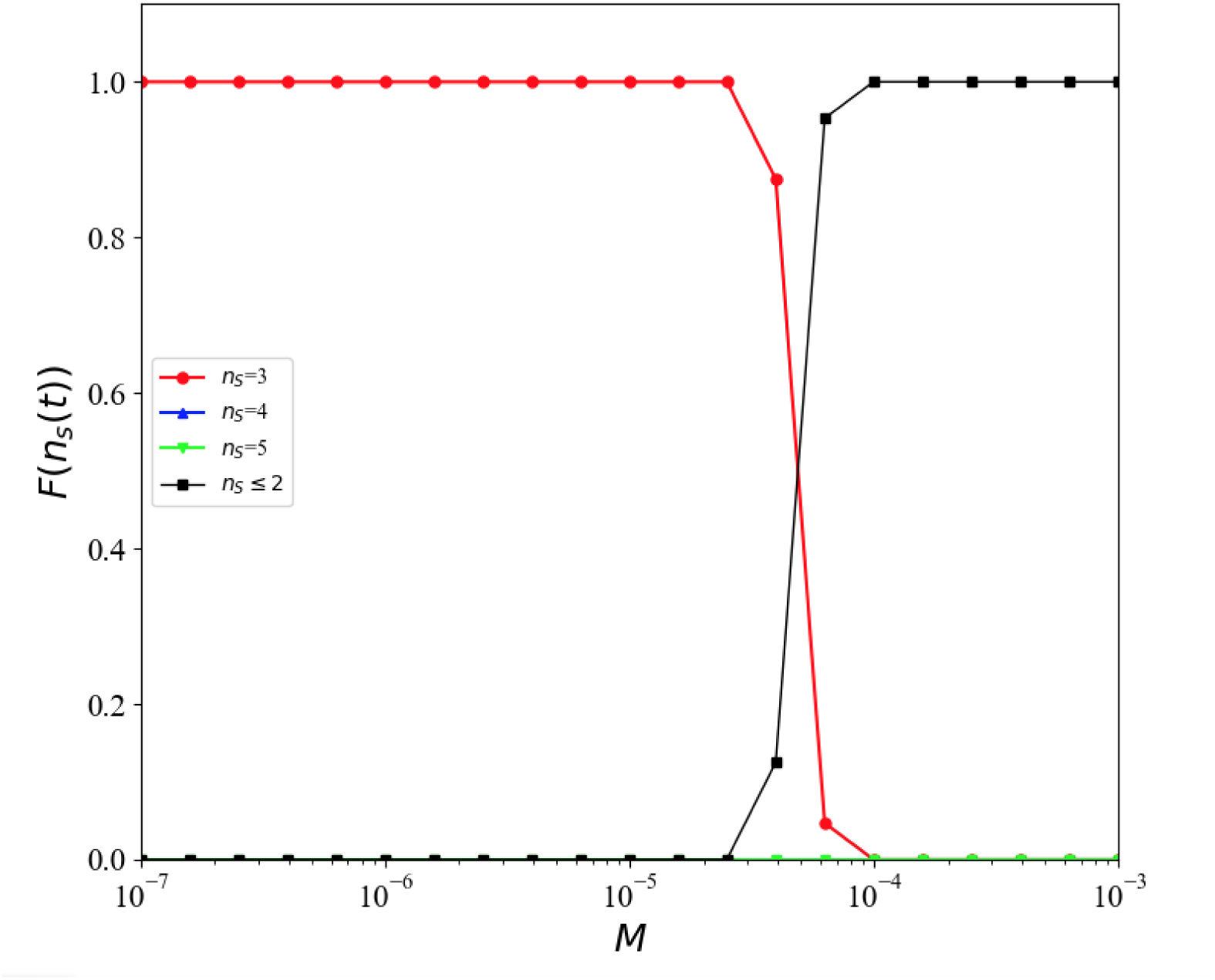
Plot of *FvsM* for the true asymptotic behavior of the Z0 unablated dominance network (i.e., the original Rpsls system) predicted/projected from the data previously shown in Figures 16, 17, and 18. The *F* values for *M<*6.31×10*^−^*^4^ are projected results for 10^6^*<t*_max_≤10^7^; and the *F* values for *M* ≥6.31×10*^−^*^4^ are projections from the results for 10^7^*<t*_max_≤3×10^7^ shown in Figure 18. Format and legend labels as for Figure 4.

To summarise the critique thus far, for ease of comparison Figure 20 shows again the true asymptotic *FvsM* plots for each of networks Z0, Z1, Z2a, and Z3b. These are all essentially the same, and so it seems that the introduction of one or more ablations to the Rpsls network does not qualitatively alter the asymptotic outcomes of the system, but instead merely changes the number of mcs time-steps needed to be simulated for the system to fully converge onto its true asymptotic state.

**Figure 20:**
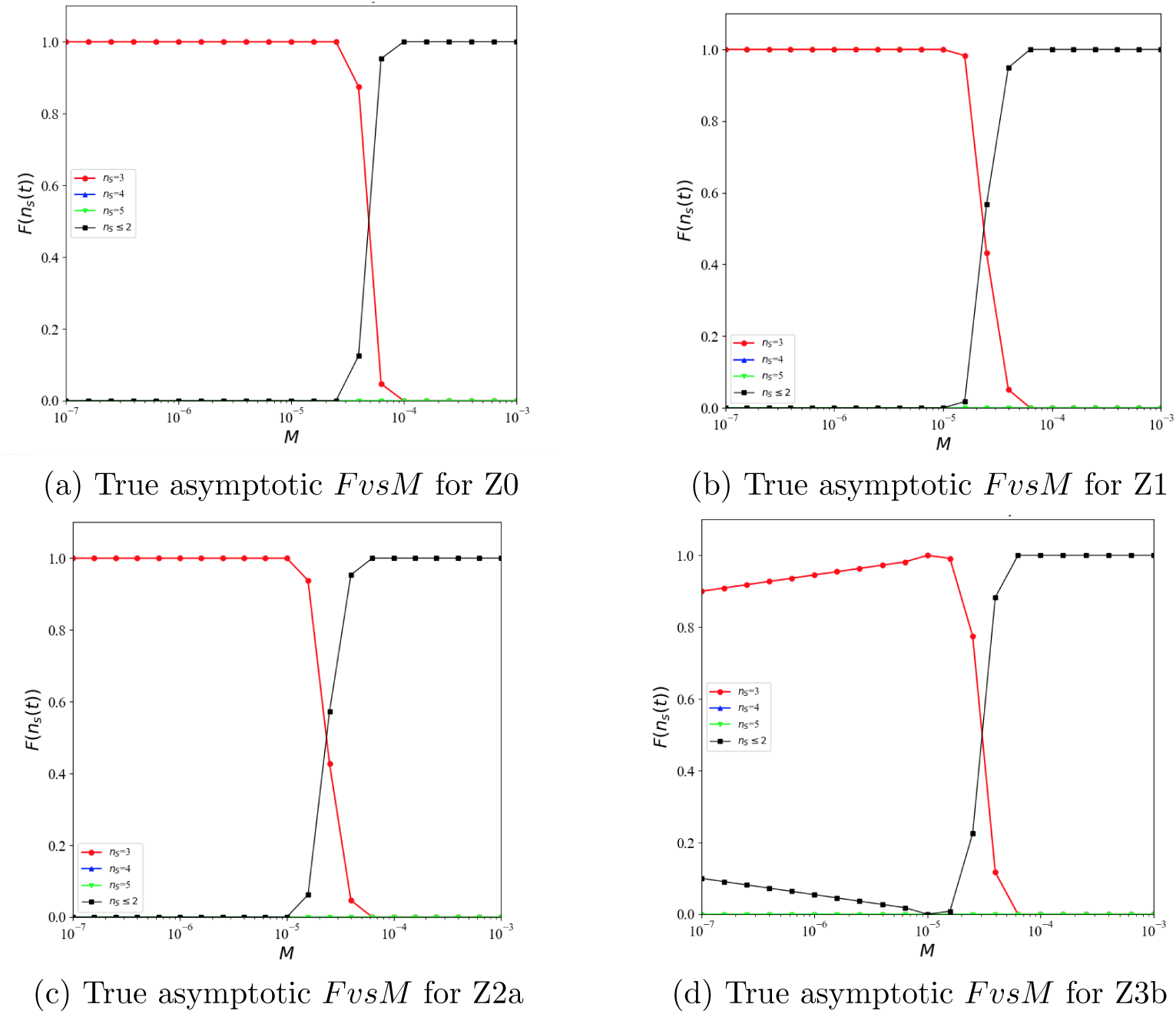
Summary *FvsM* plots of true asymptotic behaviors of the Rpsls system with ablated dominance networks: (a) Z0 (shown previously in Figure 19); (b) Z1 (at *t*_max_=5 10^5^mcs, shown previously in Figure 10); (c) Z2a (at *t*_max_=10^6^mcs, shown previously in Figure 8d); (d) Z3b (shown previously in Figure 15). As can be seen, in all four graphs, for *M* below some threshold value *M_θ_*≈10*^−^*^5^ the outcome is always *n_s_*(*t*)=3 for Z0, Z1 and Z2a, and *n_s_*(*t*)=3 in ≥90% of the Z3b outcomes; and then once *M* ≥*M_θ_* there is a sharp transition to outcomes always being *n_s_*(*t*)≤2 for all four systems. Format and legend labels as for Figure 6.

## 7. What about Z4?

Understanding why Z4 gives the results shown in Figure 7 remains something of a work in progress, and for that reason Z4 is dealt with separately here.

Zhong et al. in [1] offered no discussion of potential causal explanations for why the Z4 results are as they are, opting instead to offer only a verbal description of features manifestly evident from visual inspection of the *FvsM* plot. Discussing their Z4 results, the entirety of Zhong et al.’s narrative on the *FvsM* plot is as follows:

“In this case, only one prey-predator interaction remains in the alternative competition. We consider its representative example shown in Fig.1(e) [*reprinted here as Z4 in* Figure 3]. The results are presented in Fig.7 [*reprinted here as the upper plot in* Figure 7]. The possible asymptotic behaviors include five-species state, three-species state (Rock-Lizard-Paper), two-species states and single-species states. The latter two are the states of noninteracting species. Interestingly, the state of noninteracting species may exist at low mobility whose occurrence displays a non-monotonic behavior against the mobility. The competition between three-species state and five-species state also induces interesting behaviors. The occurrence of the five-species state displays a maximum at around *M* =2 10*^−^*^6^ In contrast, the occurrence of the three-species state shows a dip at around *M* =2 × 10*^−^*^6^ and reaches its maximum at around *M* =3 × 10*^−^*^5^ before yielding to the state of non-interacting species.” [1, Section 3.4]

Figure 21 shows plots of *FvsM* for outcomes from Z4 simulations, results from which were shown in Zhong et al.’s Figure 7, for two different durations of experiment: the upper graph is from my experiments with *t*_max_=170kmcs, which gives the best match to Zhong et al.’s results; the lower graph is from my experiments with *t*_max_=10^6^mcs.

**Figure 21:**
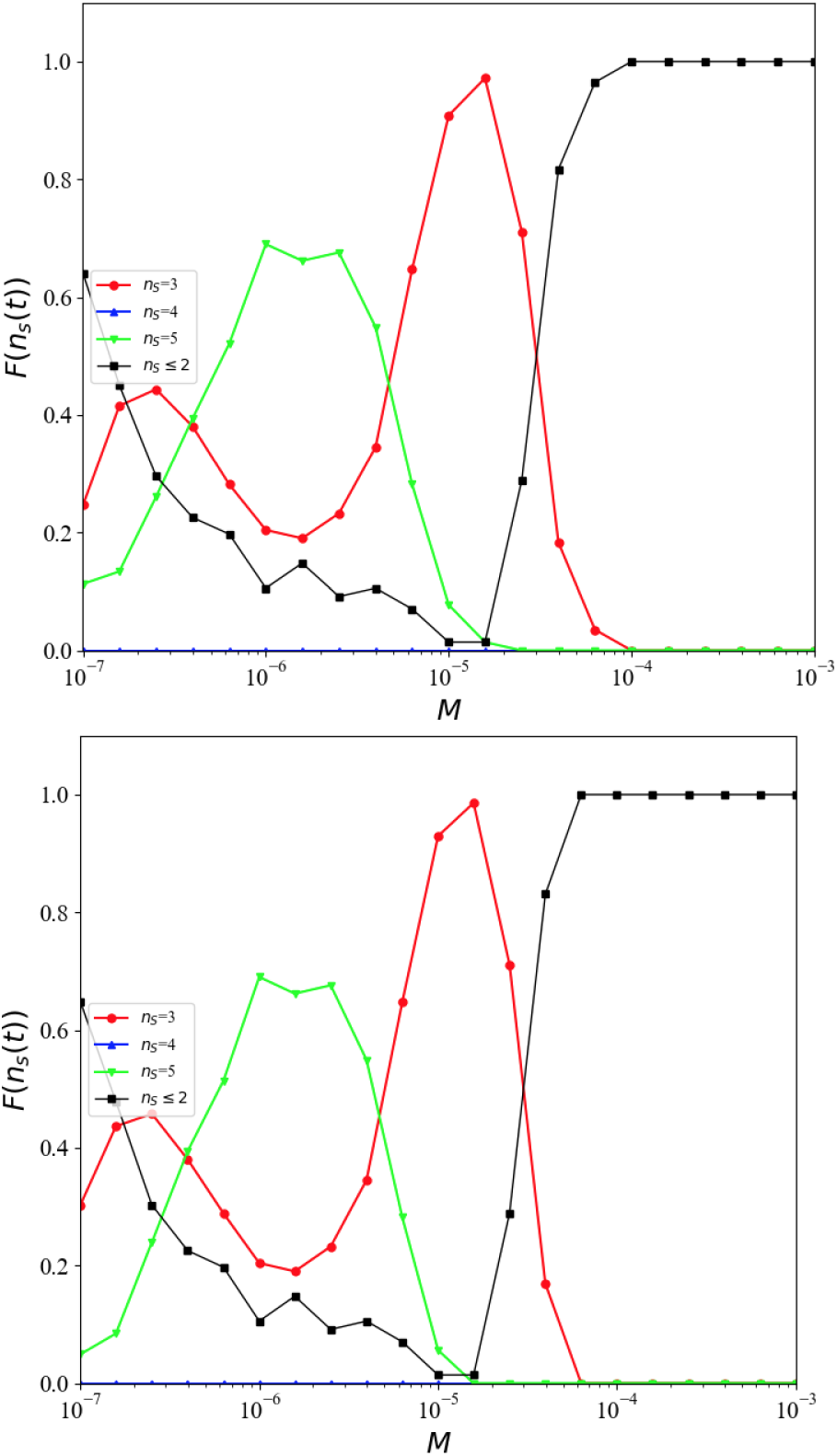
Plots of *FvsM* for the *N_a_*=4 Rpsls dominance network Z4, results from which were shown in Zhong et al.’s Figure 7; *N* =200×200, for two different durations of experiment. Upper graph is from experiment duration *t*_max_=170kmcs, which gives the best match to the results published by Zhong et al. in [1]; lower graph is from experiment duration *t*_max_=10^6^mcs. The graph for *t*_max_=10^5^mcs shows a large “bump” in *F* (*n_s_*(*t*_max_)=5) (the frequency of occurrence of the experiment ending with five-species coexistence) centred at *M* 2 10*^−^*^6^. The longer run-time brings some small reductions in *F* (*n_s_*(*t*_max_)=5) at *M* =10*^−^*^7^ and at *M* =10*^−^*^5^, but the bulk of the ‘bump’ is otherwise essentially unchanged. Format and legend labels as for Figure 4.

As with Zhong et al.’s Z4 *FvsM* plot, my replication’s *FvsM* plot for *t*_max_=10^5^mcs shows a large “bump” in *F* (*n_s_*(*t*_max_)=5) (i.e., the frequency of occurrence of five-species coexistence) centered at *M* 2 10*^−^*^6^. The longer run-time brings some small reductions in *F* (*n_s_*(*t*_max_)=5) at *M* =10*^−^*^7^ and at *M* =10*^−^*^5^, but the bulk of the bump in *F* (*n_s_*(*t*_max_)=5) is otherwise unchanged. *Prima facie*, this appear likely to be another case where, when the simulation is run for longer, the bump will disappear. Additional runs were executed with the Z4 system at *t*_max_=10^7^mcs, i.e. running the system for an additional nine million mcs but this had very little impact on the shape, the gross morphology, of the bump in *F* (*n_s_*(*t*_max_)=5) values on the *FvsM* graph.

However, Figure 22 shows *FvsT* plots for two values of *M* at *t*_max_=10^7^: in both plots, the system appears to have settled to an asymptotic equilibrium with *F* (*n_s_*(*t*)=*c*) for *c* 2, 3, 5 flatlining over a long period of apparent stasis over 3 10^4^*<t<*10^6^, but then at *t* 1.5 10^6^ in the upper plot and at *t>*6 × 10^6^ in the lower plot, we start to see a reduction in *F* (*n_s_*(*t*)=5) and commensurate rises in the frequencies of two-species and three-species co-existence. Clearly this is a situation in which running the simulation to *t*_max_=10^8^ or *t*max=10^9^ would be necessary to identify the true asymptotic behaviors, but where again it seems reasonable to conjecture that the eventual true asymptotic behavior will be for *F* (*n_s_*(*t*)=5) 0 as *t* . That would mean the ‘bump’ of five-species outcomes disappears, but it would not explain why the frequency of *n_s_*(*t*_max_) ≤ 2 outcomes for Z4 is ≈0.6 at *M* =10*^−^*^7^, falling steadily to near-zero at *M* =10*^−^*^5^, before sharply rising to unity by *M* =10*^−^*^4^: explaining this multi-phasic response in the *F* (*n_s_*(*t*_max_) ≤ 2) (which is also evident, albeit to a much lesser extent, in the Z3b asymptotic behavior *FvsM* plot of Figure 20d) remains one topic for further work. Additional topics for further work are discussed in the next section.

**Figure 22:**
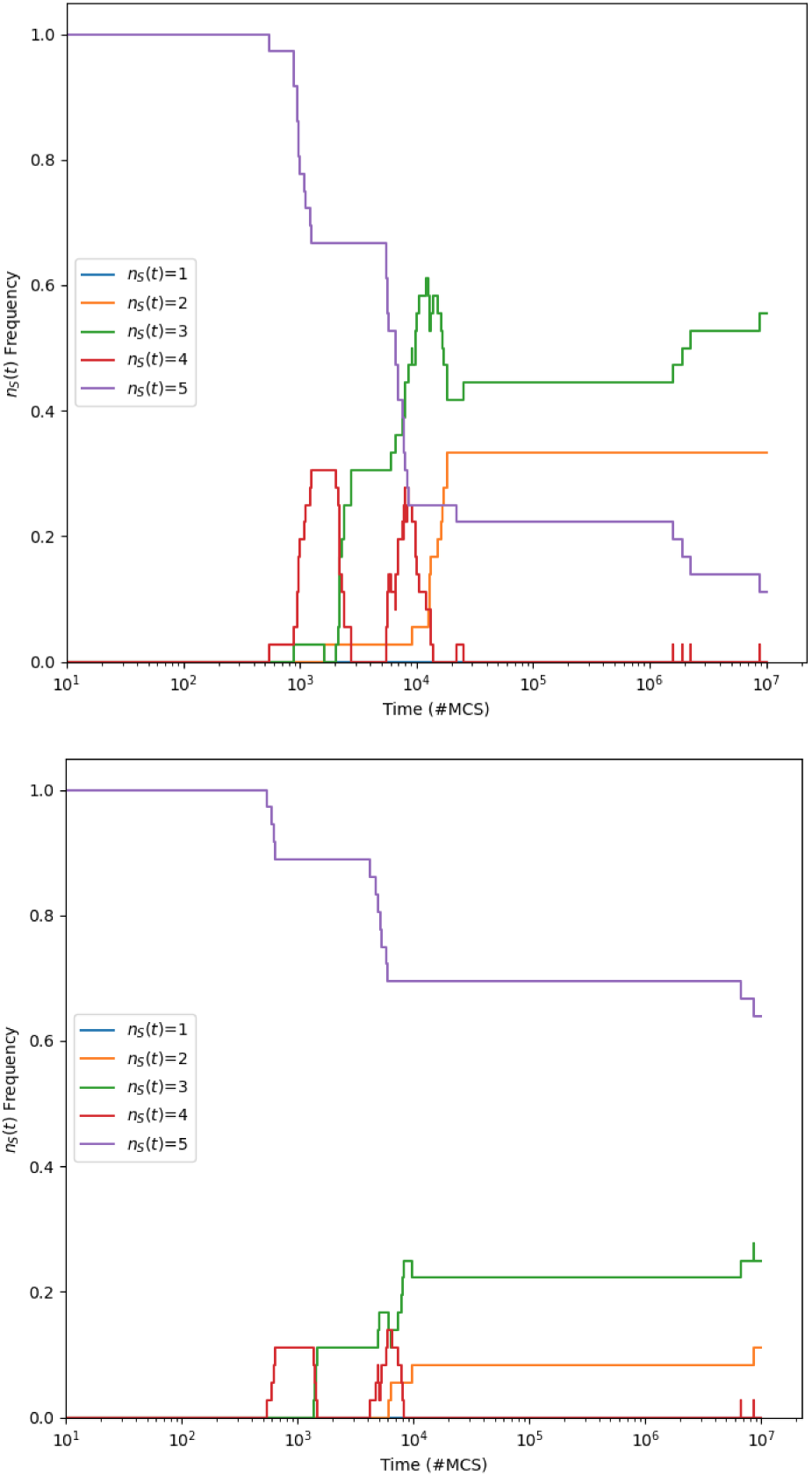
Plots of *FvsT* from the Z4 system at *t*_max_=10^7^mcs for values of *M* in the range over which the “bump” in *F* (*n_s_*(*t*)=5) was prominent in the *FvsM* plots of Figure 21. Upper graph shows *FvsT* for *M* =6.3 10*^−^*^7^; lower graph shows *FvsT* for *M* =6.3 10*^−^*^7^. Format and legend labels for both graphs are as for Figure 9. In both graphs there is a long period of equilibrium over 3×10^4^*<t<*1.5×10^6^, which could be mistaken for asymptotic behavior, but then from *t*≈1.5×10^6^ in the upper graph and *t*≈6×10^6^ in the lower graph, the value of *F* (*n_s_*(*t*)=5) starts to drop, with commensurate rises in *F* (*n_s_*(*t*)=3) and *F* (*n_s_*(*t*)=2).

## 8. Discussion and Further Work

The primary aims of this paper were to independently reproduce the results of Zhong et al. published in [1], to identify the true asymptotic behaviors for the ablated Rpsls systems, and to compare those to the true asymptotic behavior of the original unablated Z0 network. The results presented thus far have achieved all of those aims. Conclusions drawn from my results are discussed in Section 9.

Working on this replication has felt reminiscent of a notable replication from more than 30 years ago, when a paper by Nowak and May [28] was replicated by Huberman and Glance [29]. The Nowak and May paper, which was the front-cover feature article in *Nature*, explored the dynamics of another discrete-time Cartesian-lattice-based evolutionary spatial game system, based not on Rock-Paper-Scissors but instead on the iterated Prisoner’s Dilemma (see e.g. [30, 31]): Nowak and May noted that visualizations of the time-varying spatial patterns of cooperators and defectors distributed over the lattice in their experiments changed chaotically and wrote:“*… these ever-changing sequences of spatial patterns – dynamic fractals – can be extraordinarily beautiful, and have interesting mathematical properties*” [28, Abstract]. Not long later, Huberman and Glance [29] published their replication, showing that the “dynamic fractal” features highlighted by Nowak and May were simple artefacts of Nowak and May’s decision to implement their evolutionary spatial game systems using a synchronous update rule (where, as the discrete-time system transitioned from time-step *t* to *t*+1, every cell in the lattice would – in perfect synchrony – update to its new state for time *t*+1 on the basis of the states it observed in its neighbours at time *t*): when Huberman and Glance altered this aspect of the simulation to instead use asynchronous updating, updating one randomly-chosen cell at a time (as is used in the experiments reported in this paper), every cell in the lattice quickly evolved to be defector, and the intricate spatiotemporal patterns identified by Nowak and May disappeared, visualizations of the lattice becoming instead a single static solid slab of one color. In concluding their paper, Huberman and Glance wrote “*These results show that while computer experiments provide a versatile approach for studying complex systems, an understanding of their subtle characteristics is required in order to reach valid conclusions about real-world systems.”*

Those words are quite definitely just as relevant here and now as they were when first written in 1993: in this paper I have demonstrated that there are subtle characteristics in the ablated Rpsls systems introduced by Zhong et al., characteristics that only reveal themselves when a rigorously principled approach is taken to identifying the true asymptotic outcomes for these systems, and when the outcomes of those ablation treatments are compared to an appropriate control, which in this case is the result from the relavent unablated system, the Z0 network. Conclusions drawn from those results are given below in Section 9.

This paper serves to correct the publiushed record on the asympotic outcomes of the ablated Rpsls systems introduced by Zhong et al. but there is clearly more work that could be done to take this line of enquiry further. Current research is proceeding on two fronts. One strand of work is exploring the extent to which a seemingly minor revision to the Elementary Step (Algorithm 3), reported in [4], reduces volatility in the population dynamics of the ablated Rpsls systems and significantly alters the frequency distribution of observed outcome for all the ablated systems studied in [1], with extinctions being much rarer events: results from those studies will be released in [32]. A second strand of work is exploring the frequency distributions of outcomes in Rpsls-style systems with ablated dominance networks but where *N_s_>* 5, such that any one arc ablation represents a smaller percentage loss of arcs from the overall network: early results from this work have been published in [33] and a more extensive study is currently being worked on, to be released as [34].

One additional avenue for exploration in future work is to explore and identify what gives rise to the “punctuated equilibria” nature of many of the *FvsT* plots presented here, where the distribution of frequencies of species-counts over a large number of iid simulation experiments is stable for long periods and then undergoes a (relatively) brief period of sudden change, indicating that all the iid experiments underwent a species extinction at roughly the same time. For example: in Figure 9, for the first ≈150mcs all 500 iid experiments hold steady at *n_s_*(*t*) = 5 but then over the period *t* ∈ [150, 1000]mcs, every one of the simulations undergoes its first extinction such that by *t*=1000*, F* (*n_s_*(*t*)=5)=0 and *F* (*n_s_*(*t*)=4)=1; the upper plot *FvsT* plot in Figure 12 shows a similar sequence of events, with *F* (*n_s_*(*t*)=5) collapsing from unity to zero and *F* (*n_s_*(*t*)=4) rising rapidly from zero to approach 1.0 over the period running roughly *t* ∈ [500, 2500]; and to give a final example, in the upper *FvsT* plot of Figure 14 there is a long equilibrium running from *t*≈10^4^ to *t*≈10^5^ where *F* (*n_s_*(*t*)=5)≈0.35 and *F* (*n_s_*(*t*)=3)≈0.65 and then, from *t*≈10^5^ onwards, *F* (*n_s_*(*t*)=5) starts to fall steadily and hence *F* (*n_s_*(*t*)=3) starts to rise. In all three of these examples, and in every other *FvsT* plot that shows a similar long-standing equilibrium that is then “punctuated” by temporally clustered extinctions occurring across the set of iid simulations, presumably during the period of equilibrium there is a some sequence of changes in the lattice, gradually moving the state of the system toward some kind of tipping-point which, once crossed, brings the equilibrium to an end, and this happens on roughly the same timescale for all iid repetitions of the specific simulation. These observations prompt a set of related research questions: precisely *what* is it about the state of the lattice that is changing during the equilibrium periods; what is the best metric by which we may measure that which is changing during the equilibria; and what is the nature of the tipping-point or threshold-crossing that triggers these extinctions that are so strongly temporally concentrated across large numbers of iid simulations? One potentially promising line of enquiry here would be to adapt and extend the work of Dong, Li, and Yang [35] who studied extinction patterns in the three-species RPS game, finding that the maximal amplitude of density fluctuation, relative to the average density, and the average Potts energy per lattice-cell are significant factors in determining extinction patterns.

## 9. Conclusion

All research in science and engineering relies on the publication of results that are replicable, and several fields of research have struggled in recent years each with their own “replicability crisis” (see e.g. [36, 37, 38, 39, 40]). I have demonstrated here in Section 4 that, with some digging through the prior literature to establish full details of the algorithms employed, the results presented by Zhong et al. in [1] can be reproduced^5^ very well: this is, in principle, a good thing.

Zhong et al.’s paper sets out the rationale for their simulation experiments, explains the simulated Escg model in some detail, plots graphs summarising the outcomes of large numbers of experiments, and then offers brief verbal descriptions of features in the plots that are immediately obvious from visual inspection of the graphs. Their narratives on the visualizations seem superficially to be offering detailed causal explanations (such as: *“it may allow for five-species coexistence since the two three-species-cyclic interactions only share one species, Lizard. Once the coexistence of Rock-Lizard-Spock becomes possible, the coexistence of five species becomes possible . . . ”*).

However, my results demonstrate that if Zhong et al. had run their experiments for longer, and/or if they had plotted *FvsT* graphs, they would have needed to radically re-write the stories they tell about the ablated systems they studied. In light of this, Zhong et al.’s narratives run the risk of seeming to be “just-so-stories”, filling a gap in the paper where a carefully argued and rigorously evaluated causal explanation would normally be expected [41].

Furthermore, as I discussed in more detail in [4], the only dominance-network ablations that Zhong et al. chose to explore in their paper are those that led to reductions in the number of co-existing species, but other equally plausible ablated Rpsls dominance networks show no such effect, suffer no extinctions, and their species-count remains constant at five for the entire duration of the experiment: at best, [1] is selective, and tells only half the story.

But the design of experiments in [1] is more deeply questionable on two counts: Zhong et al. assert, incorrectly, and without any justification or supporting evidence, that asymptotic behaviors are observable in the ablated Rpsls system at *t*_max_=10^5^; and they also chose not to show any results from the original unablated Z0 network, which would serve as a baseline (i.e., a “pre-treatment” evaluation or *control*) against which meaningful comparisons could then be made with the results from the “post-treatment” ablated systems. The reasoning behind these omissions is not clear to me, nor why no-one involved in the peer-review of the manuscript picked up on these issues. I have demonstrated here that when the ablated systems are simulated for longer the results they then produce become, if anything, *less* interesting, because the diversity of outcomes reported and narrated on by Zhong et al. disappears once each ablated system is given long enough to converge on its true asymptotic outcome: when analysed in *FvsM* form, the true asymptotic outcomes for the ablated systems Z0-Z3b are very similar to each other, and are each in turn very similar to the *FvsM* outcome for the data from the control, the unablated Z0 system.

I have shown that the results presented in Zhong et al.’s paper, and their narrative explanations of those results, are in need of revision and are not as straightforward as Zhong et al. set them out to be. When the ablated Rpsls systems are explored in more depth, for sufficient durations, and when they are then properly compared to results from the unablated Z0 as a baseline control, the actual effects of the dominance-network ablations seems to be somewhat moot, and hence the results from the ablated Rpsls systems are probably not as interesting or relevant to studies of real-world biodiversity and species coexistence as Escg research of this type is often claimed to be. I’ve made my source-code available so that other researchers can replicate, explore, and (I hope) extend the work presented here. The appendices present additional illustrative *FvsT* plots, for reference.

**Funding:** computational resources used for the simulation experiments reported here were provided in part by Syritta Ltd, who paid for AWS cloud usage and also provided multiple 8-core Apple Mac computers. Other than that, this research did not receive any specific grant from funding agencies in the public, commercial, or not-for-profit sectors.

## Appendix A. Dynamics of *N_a_*=0 Rpsls Z0 with *µ*=*σ*=1.0, *L*=200

Figures A.23 to A.29 show time-series of the evolution the number of surviving species *n_S_*(*t*) over 10^6^mcs for the OES Escg with the unablated (*N_a_*=0) dominance network Z0 as illustrated in Figure 2, *L*=200 and *µ*=*σ*=1.0 for exponentially spaced values of *M* ∈ [10*^−^*^7^, 10*^−^*^3^], with 250 IID simulations executed for each value of *M* sampled.

Given that *N_a_*=0, the progressive reductions in *n_s_*(*t*) seen in all these time-series are each underflow extinctions, i.e. they are a consequence of the variation in population dynamics inherent in the five-species Rpsls model running up against the discretization limit of *L*=200: see [4] for further discussion of this point.

**Figure A.23:**
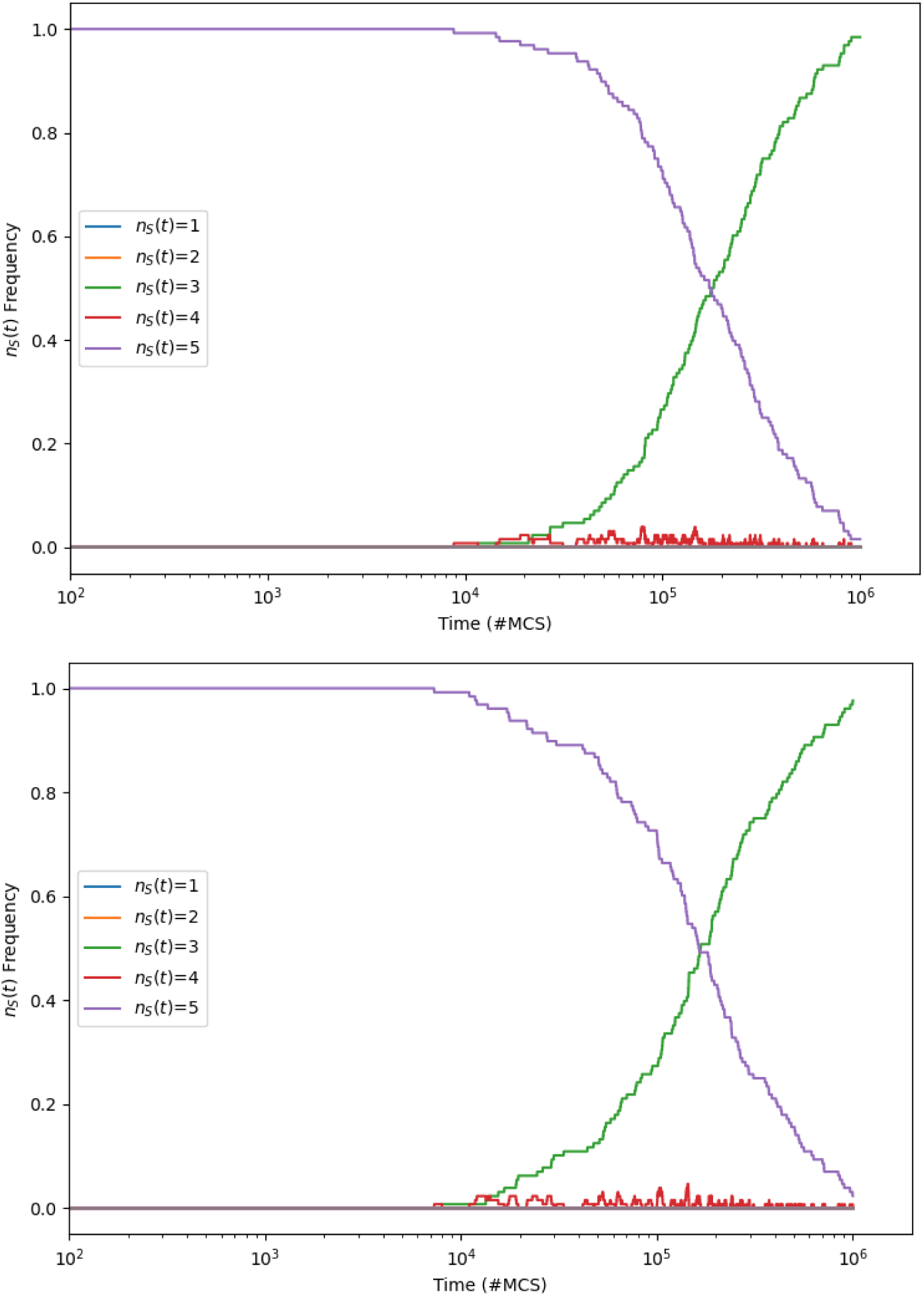
Frequency distribution of species-counts *n_s_*(*t*) : *s* ∈ {1*, . . .,* 5} against time *t* (measured in MCS) for Rpsls experiments with no ablations (*N_a_*=0): *L*=200; *µ*=*σ*=1.0; *M* =10*^−^*^7^ (upper graph), *M* =10*^−^*^6^ (lower graph). At each value of *M* sampled, 250 IID simulation experiments were performed. Horizontal axis is time, measured in mcs; vertical axis is what proportion of the simulations had *n_S_*(*t*)=*c* at each time *t* for *c*∈{1*, . . ., N_S_*}.

**Figure A.24:**
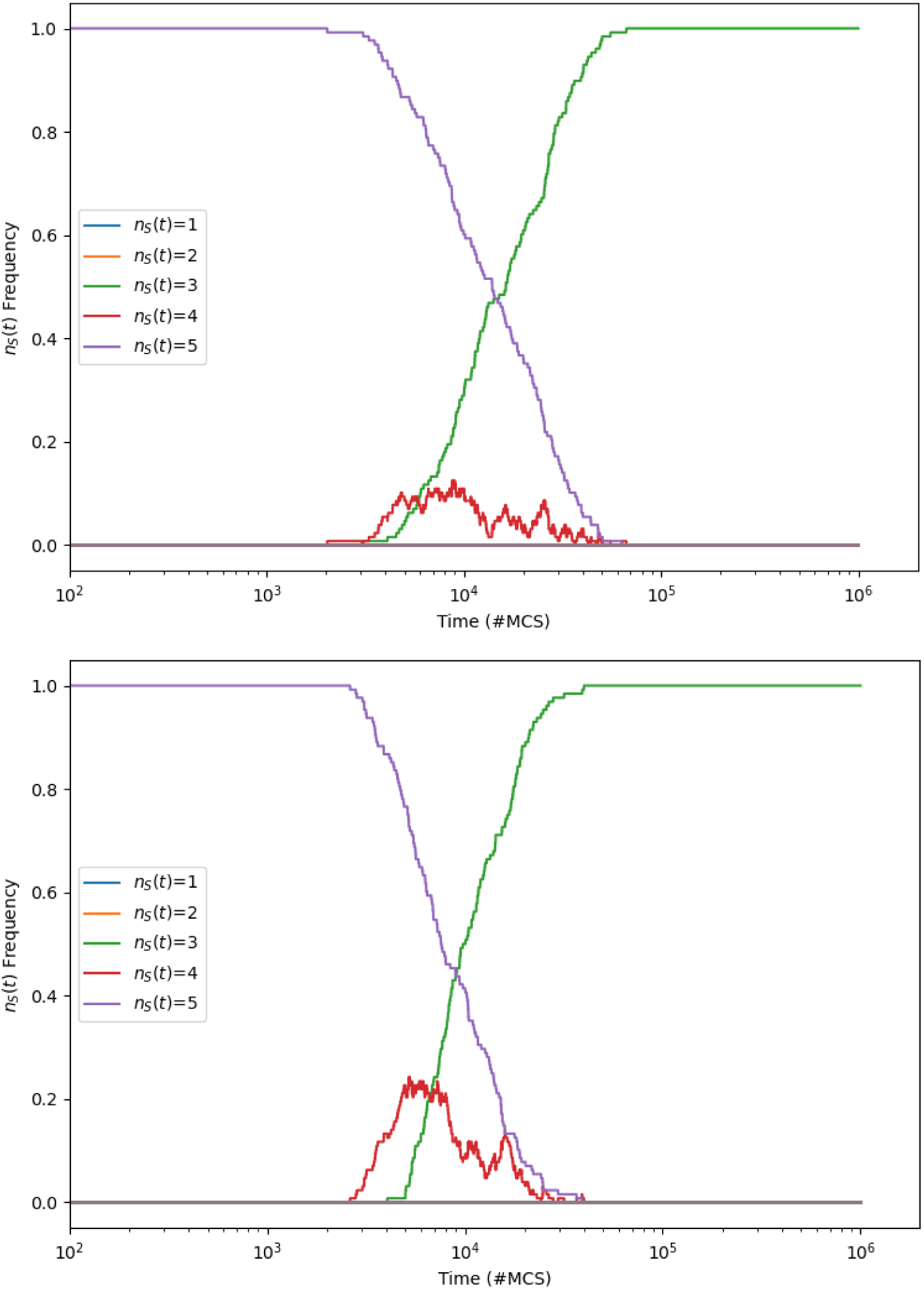
Frequency distribution of species-counts *n_s_*(*t*) against time *t* for Rpsls experiments with no ablations (*N_a_*=0): *L*=200; *µ*=*σ*=1.0; *M* =10*^−^*^5^ (upper graph), *M* =1.58×10*^−^*^5^ (lower graph). At each value of *M* sampled, 250 IID experiments were performed. Format as for Fig. A.23.

**Figure A.25:**
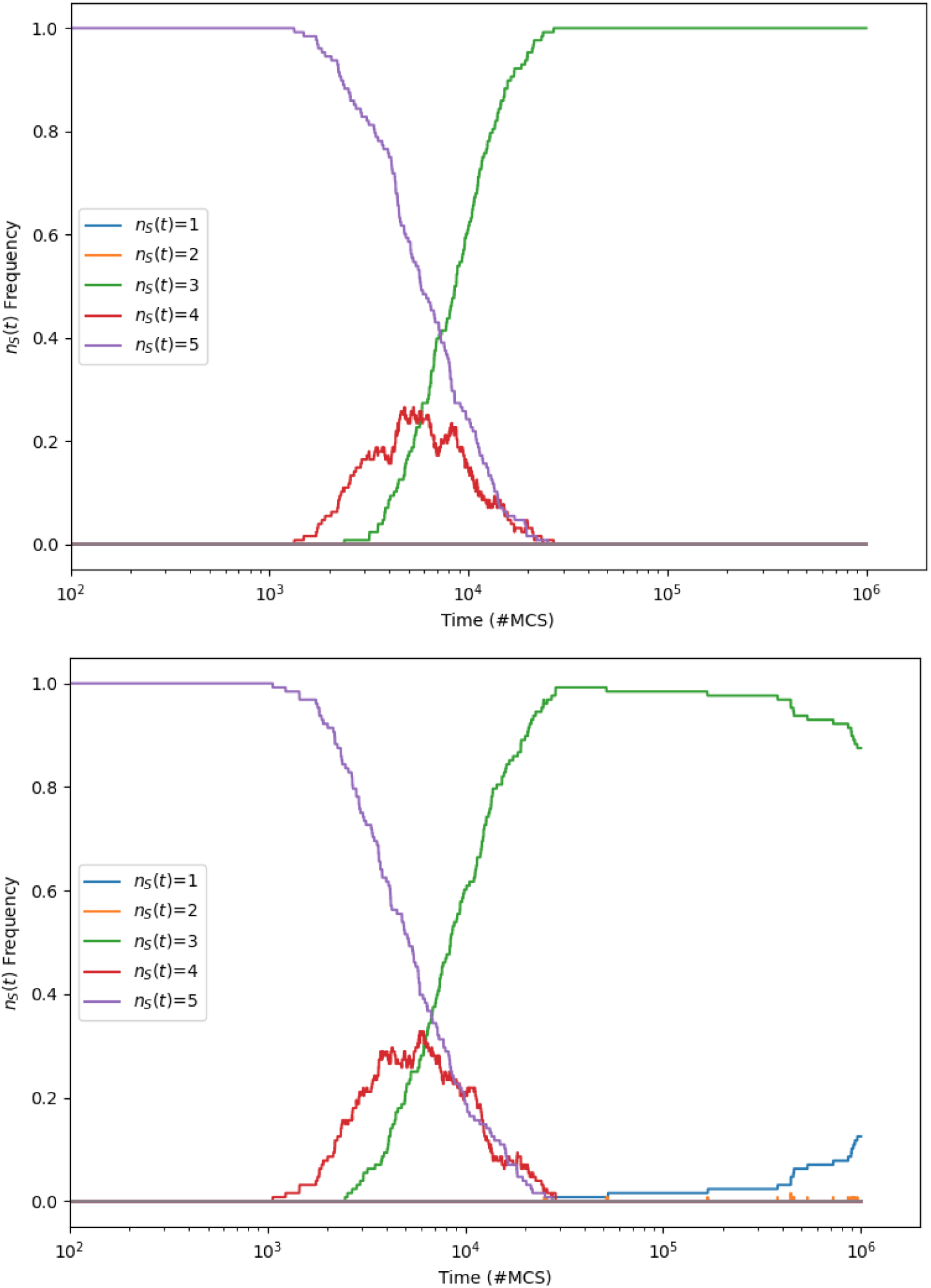
Frequency distribution of species-counts *n_s_*(*t*) against time *t* for Rpsls experiments with no ablations (*N_a_*=0): *L*=200; *µ*=*σ*=1.0; *M* =2.58×10*^−^*^5^ (upper graph), *M* =3.98×10*^−^*^5^ (lower graph). At each value of *M* sampled, 250 IID experiments were performed. Format as for Fig. A.23.

**Figure A.26:**
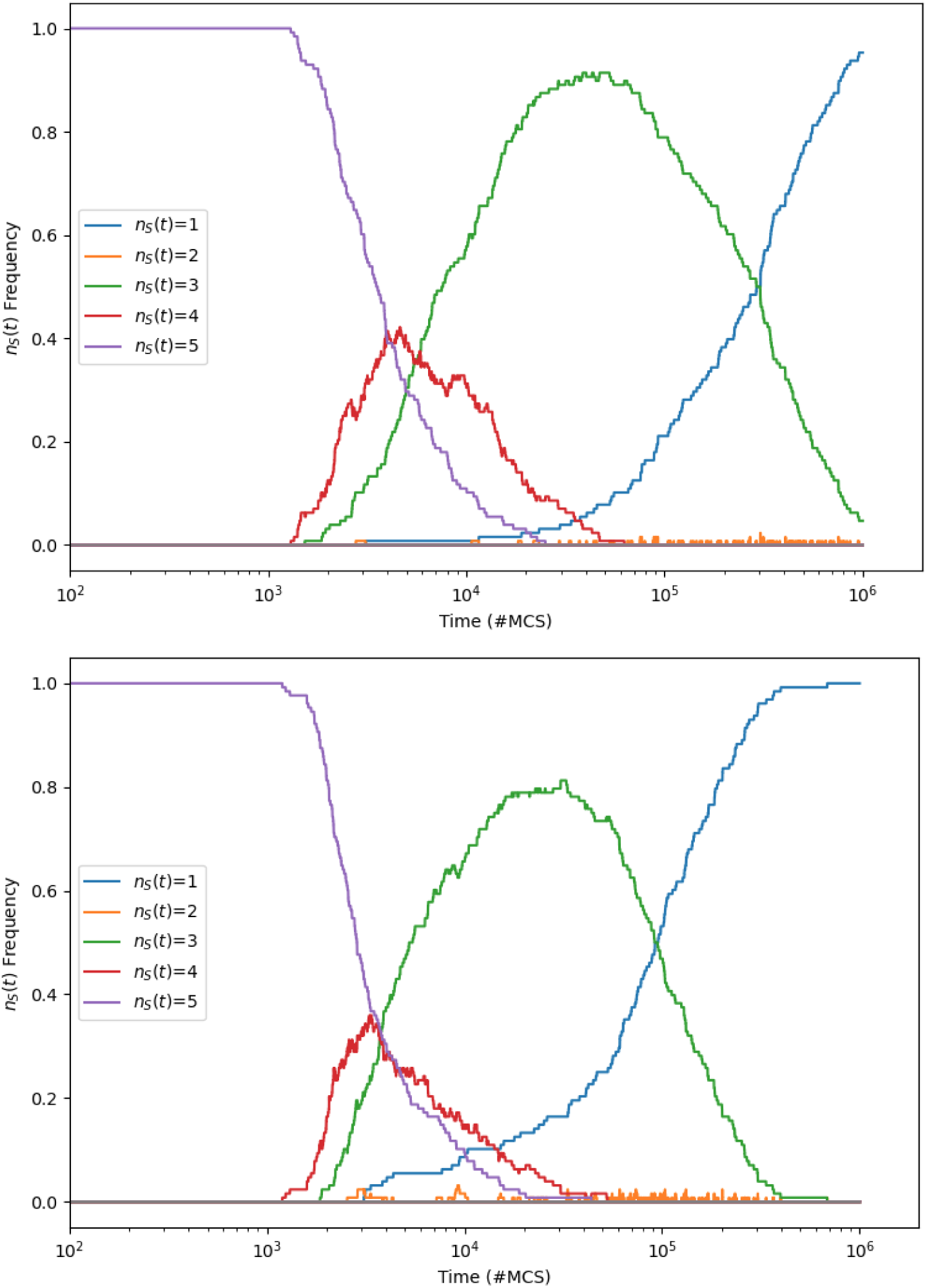
Frequency distribution of species-counts *n_s_*(*t*) against time *t* for Rpsls experiments with no ablations (*N_a_*=0): *L*=200; *µ*=*σ*=1.0; *M* =6.31 10*^−^*^5^ (upper graph), *M* =10*^−^*^4^ (lower graph). At each value of *M* sampled, 250 IID experiments were performed. Format as for Fig. A.23.

**Figure A.27:**
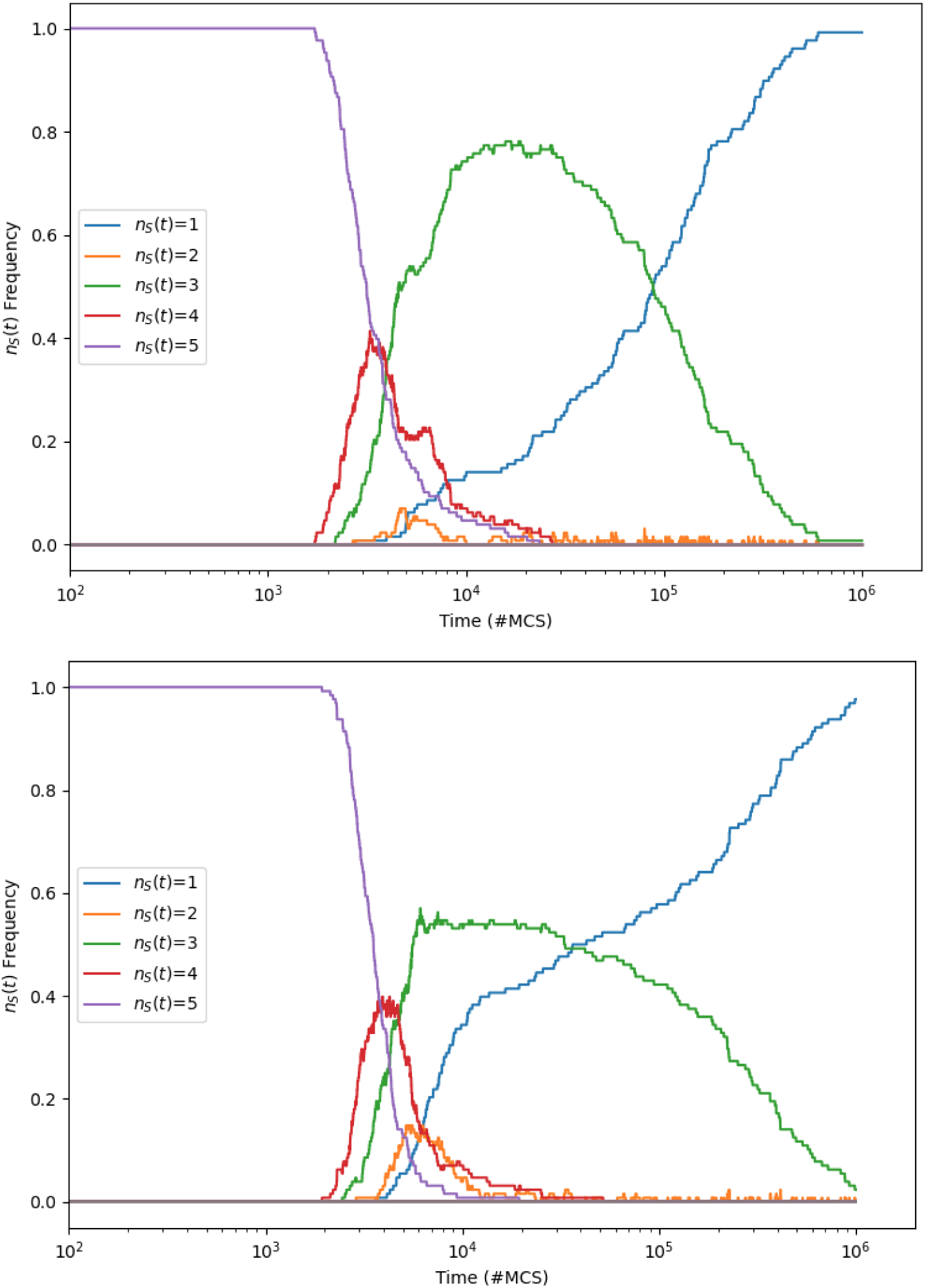
Frequency distribution of species-counts *n_s_*(*t*) against time *t* for Rpsls experiments with no ablations (*N_a_*=0): *L*=200; *µ*=*σ*=1.0; *M* =1.58×10*^−^*^4^ (upper graph), *M* =2.51×10*^−^*^4^ (lower graph). At each value of *M* sampled, 250 IID experiments were performed. Format as for Fig. A.23.

**Figure A.28:**
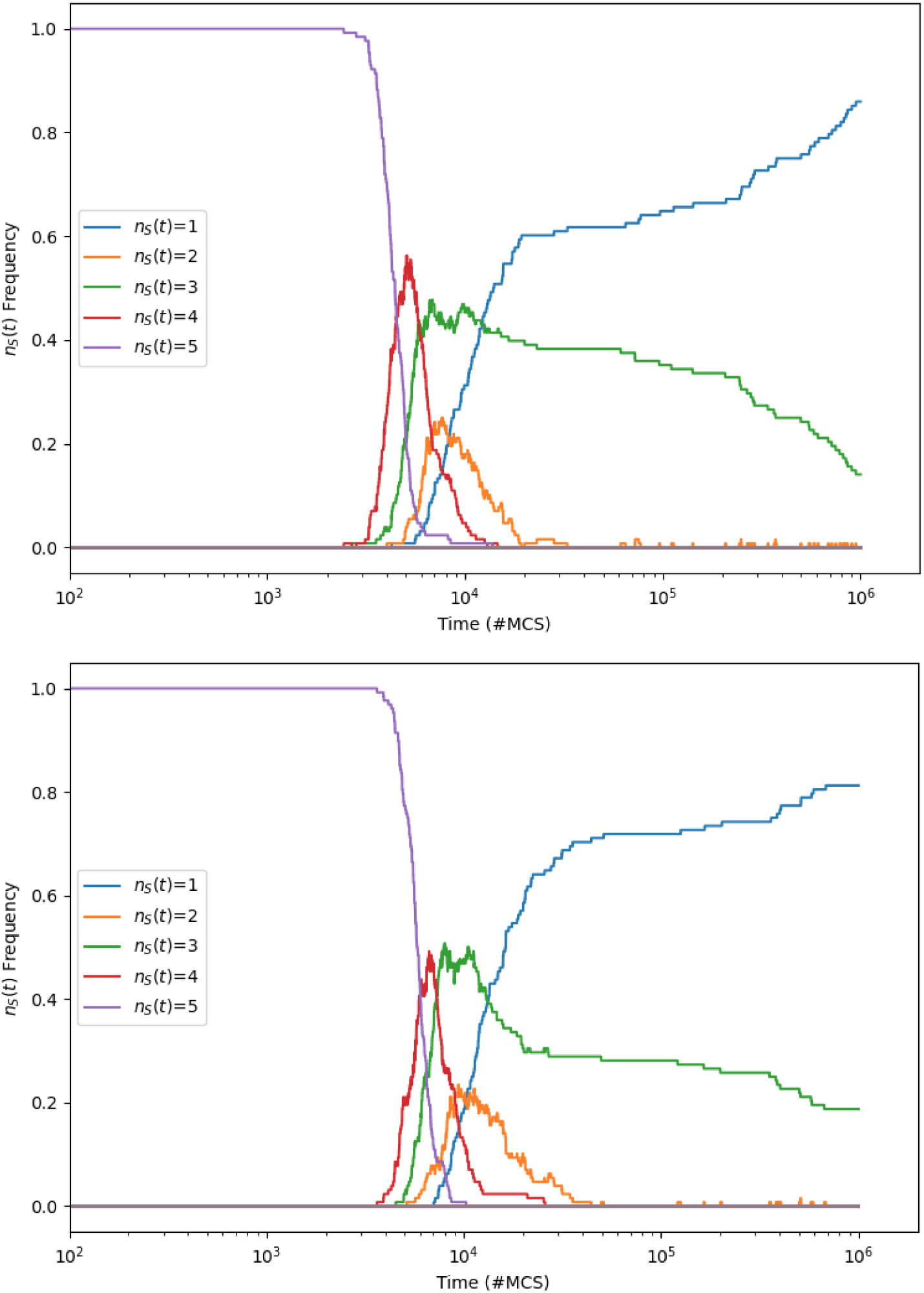
Frequency distribution of species-counts *n_s_*(*t*) against time *t* for Rpsls experiments with no ablations (*N_a_*=0): *L*=200; *µ*=*σ*=1.0; *M* =3.98×10*^−^*^4^ (upper graph), *M* =6.31×10*^−^*^4^ (lower graph). At each value of *M* sampled, 250 IID experiments were performed. Format as for Fig. A.23.

**Figure A.29:**
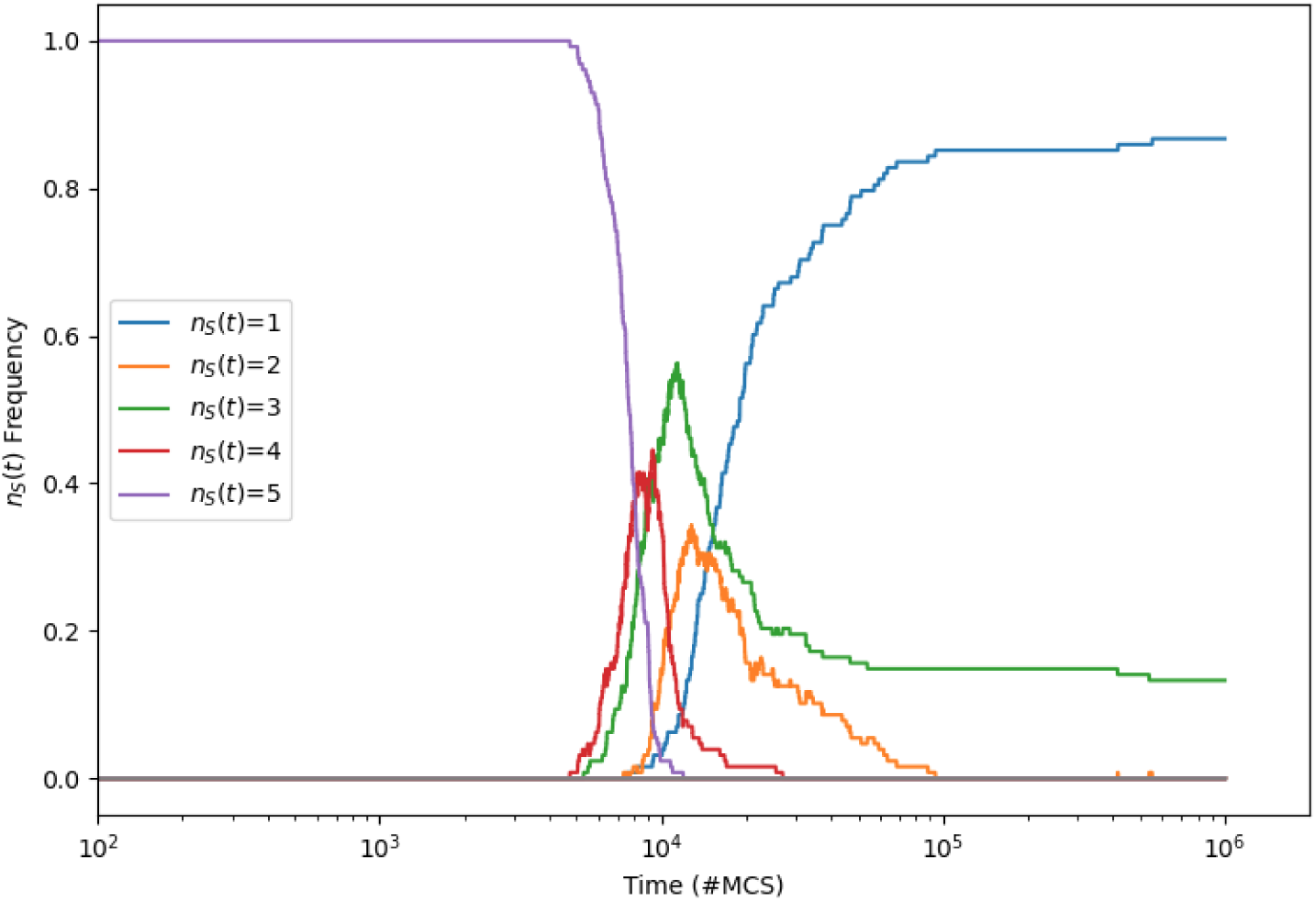
Frequency distribution of species-counts *n_s_*(*t*) against time *t* for Rpsls experiments with no ablations (*N_a_*=0): *L*=200; *µ*=*σ*=1.0; *M* =10*^−^*^3^. At each value of *M* sampled, 250 IID experiments were performed. Format as for Fig. A.23.

## Appendix B. Dynamics of *N_a_*=3 Rpsls Z3b with *µ*=*σ*=1.0, *L*=200

Figures B.30 to B.39 show time-series of the evolution of the number of surviving species *n_s_*(*t*) over 10^6^mcs for the Rpsls Escg with the three-ablation (*N_a_*=3) dominance network Z3b (as illustrated in Figure 3), *L*=200 and *µ*=*σ*=1.0 for various values of *M* ∈ [10*^−^*^7^, 10*^−^*^3^], with 250 IID simulations executed for each value of *M* sampled.

**Figure B.30:**
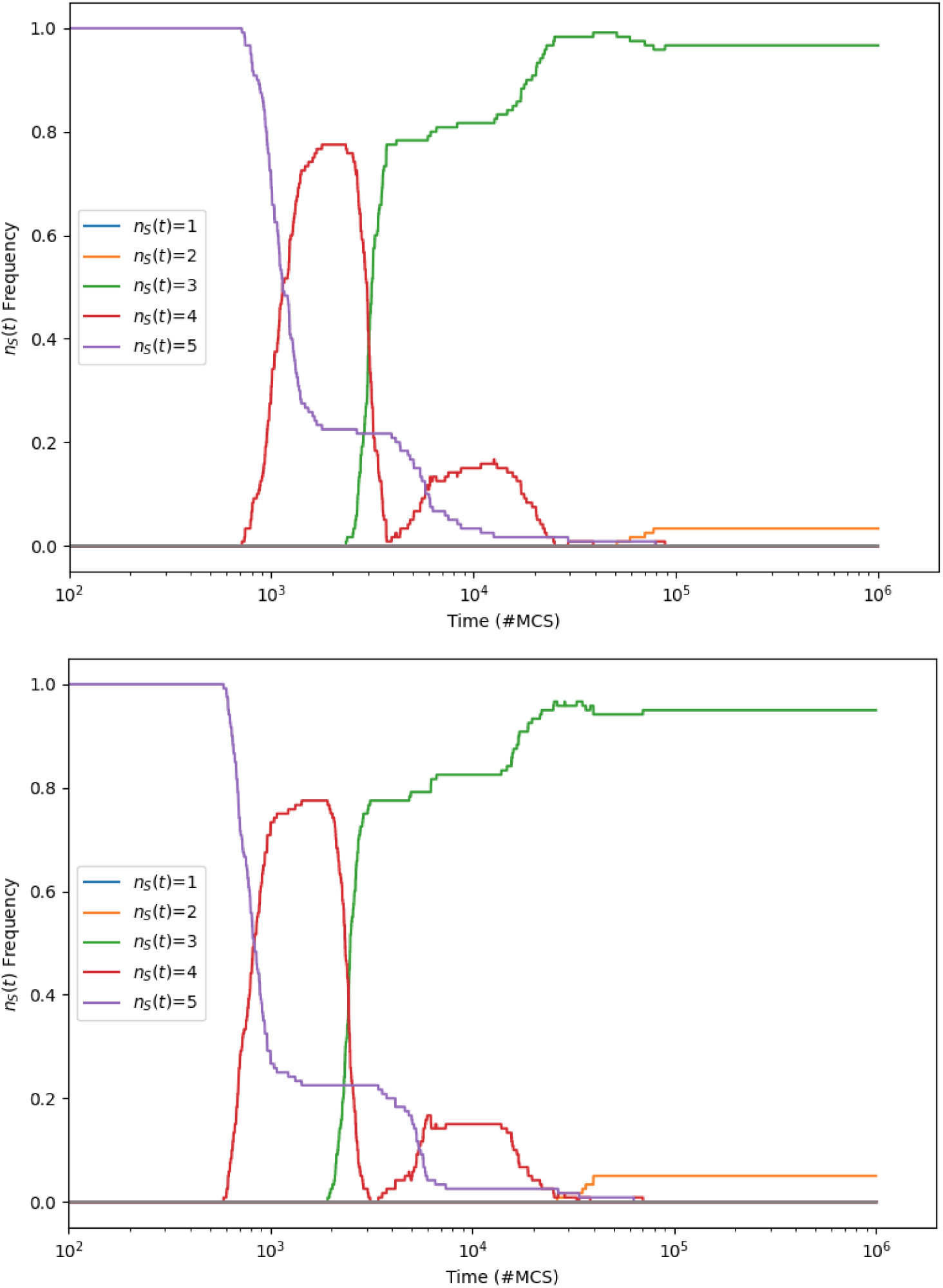
Frequency distribution of species-counts *n_s_*(*t*) : *s* ∈ {1*, . . .,* 5} against time *t* (measured in mcs) for Rpsls experiments with three ablations (*N_a_*=3): *L*=200; *µ*=*σ*=1.0; *M* =10*^−^*^7^ (upper graph), *M* =1.58 10*^−^*^7^ (lower graph). At each value of *M* sampled, 250 IID simulation experiments were performed. Format as for Fig. A.23.

**Figure B.31:**
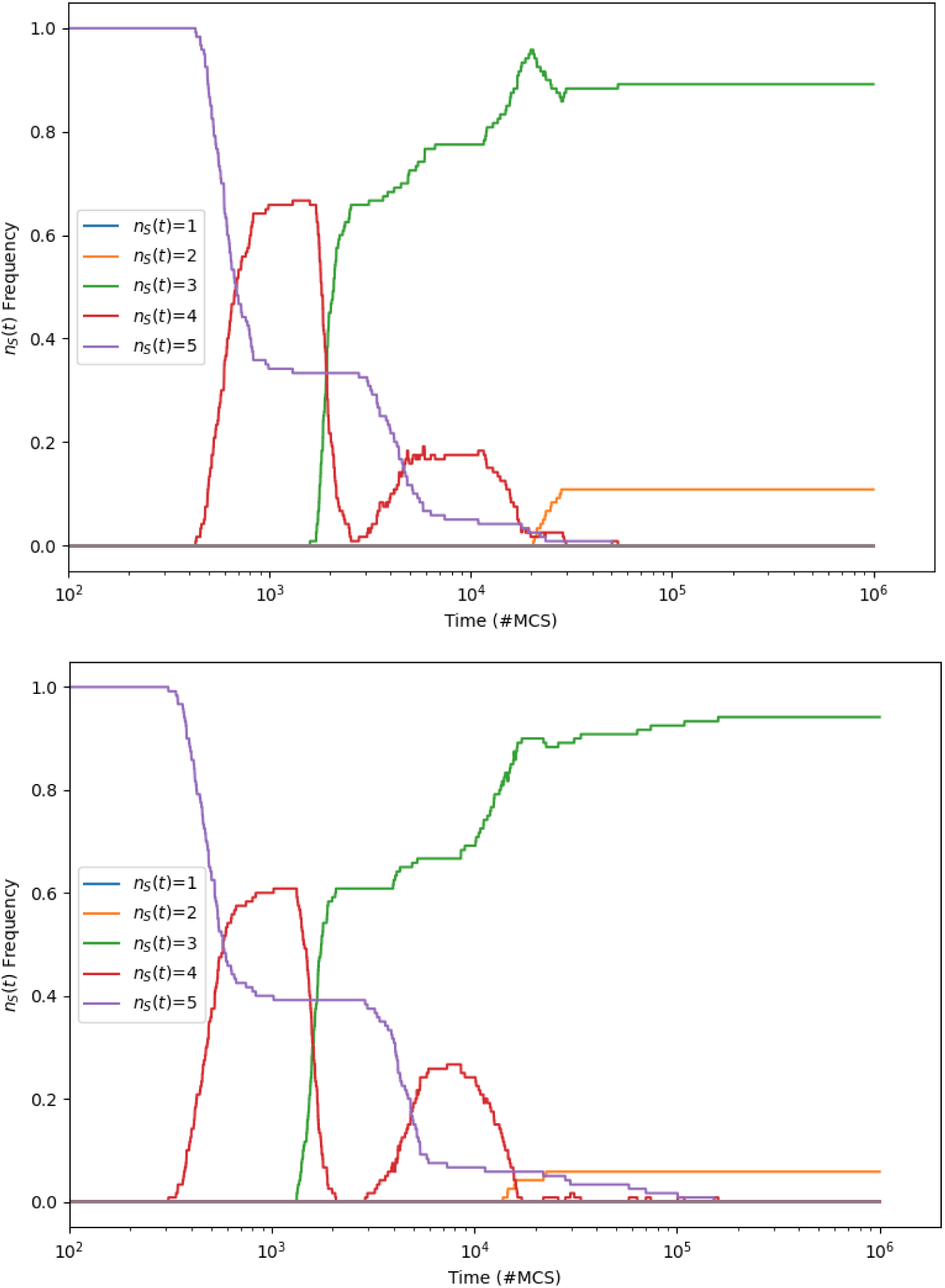
Frequency distribution of species-counts *n_s_*(*t*) against time *t* for Rpsls experiments with three ablations (*N_a_*=3): *L*=200; *µ*=*σ*=1.0; *M* =2.51 10*^−^*^7^ (upper graph), *M* =3.98 10*^−^*^7^ (lower graph). At each value of *M* sampled, 250 IID experiments were performed. Format as for Fig. A.23.

**Figure B.32:**
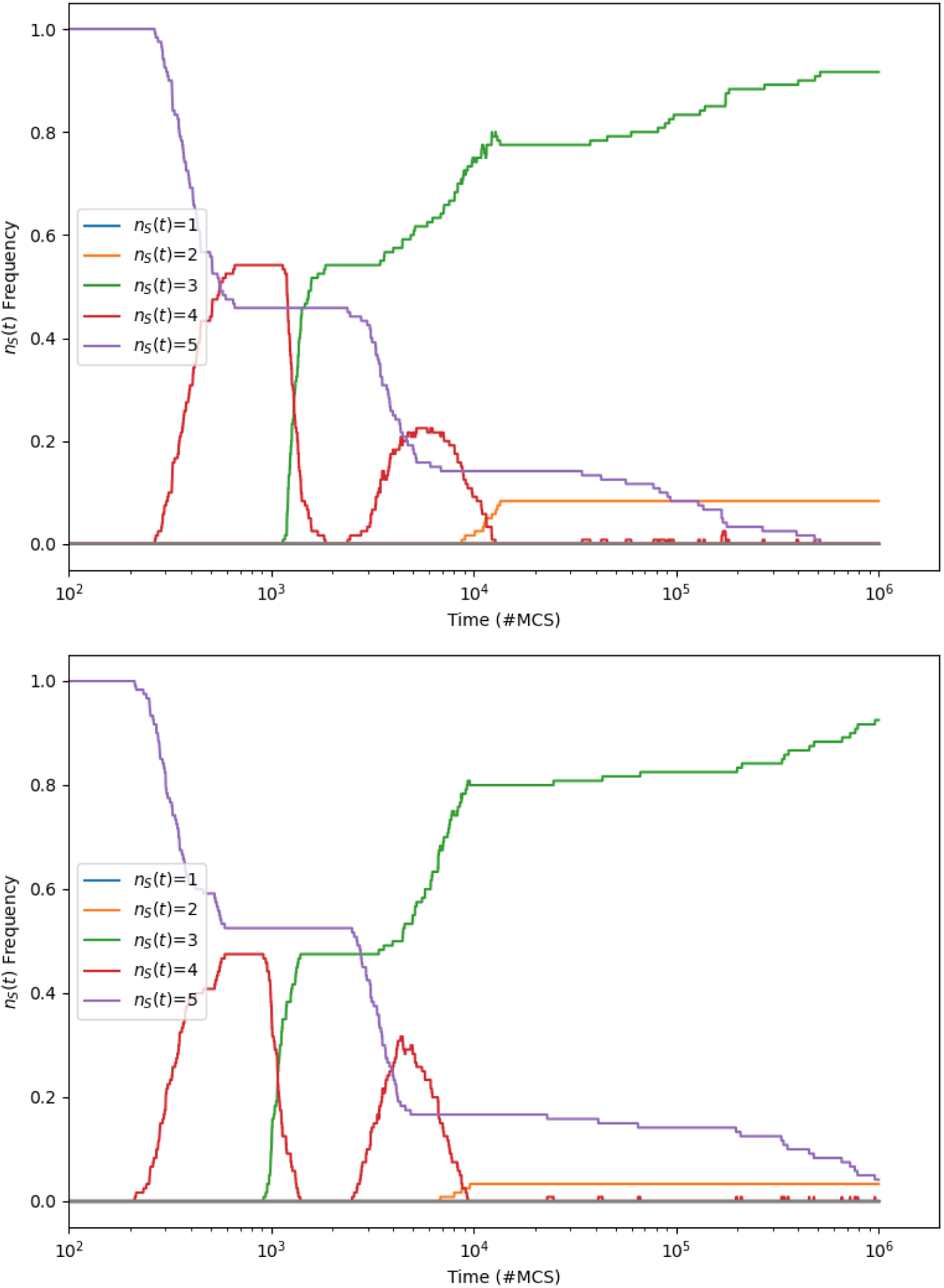
Frequency distribution of species-counts *n_s_*(*t*) against time *t* for Rpsls experiments with three ablations (*N_a_*=3): *L*=200; *µ*=*σ*=1.0; *M* =6.31 10*^−^*^7^ (upper graph), *M* =10*^−^*^6^ (lower graph). At each value of *M* sampled, 250 IID experiments were performed. Format as for Fig. A.23.

**Figure B.33:**
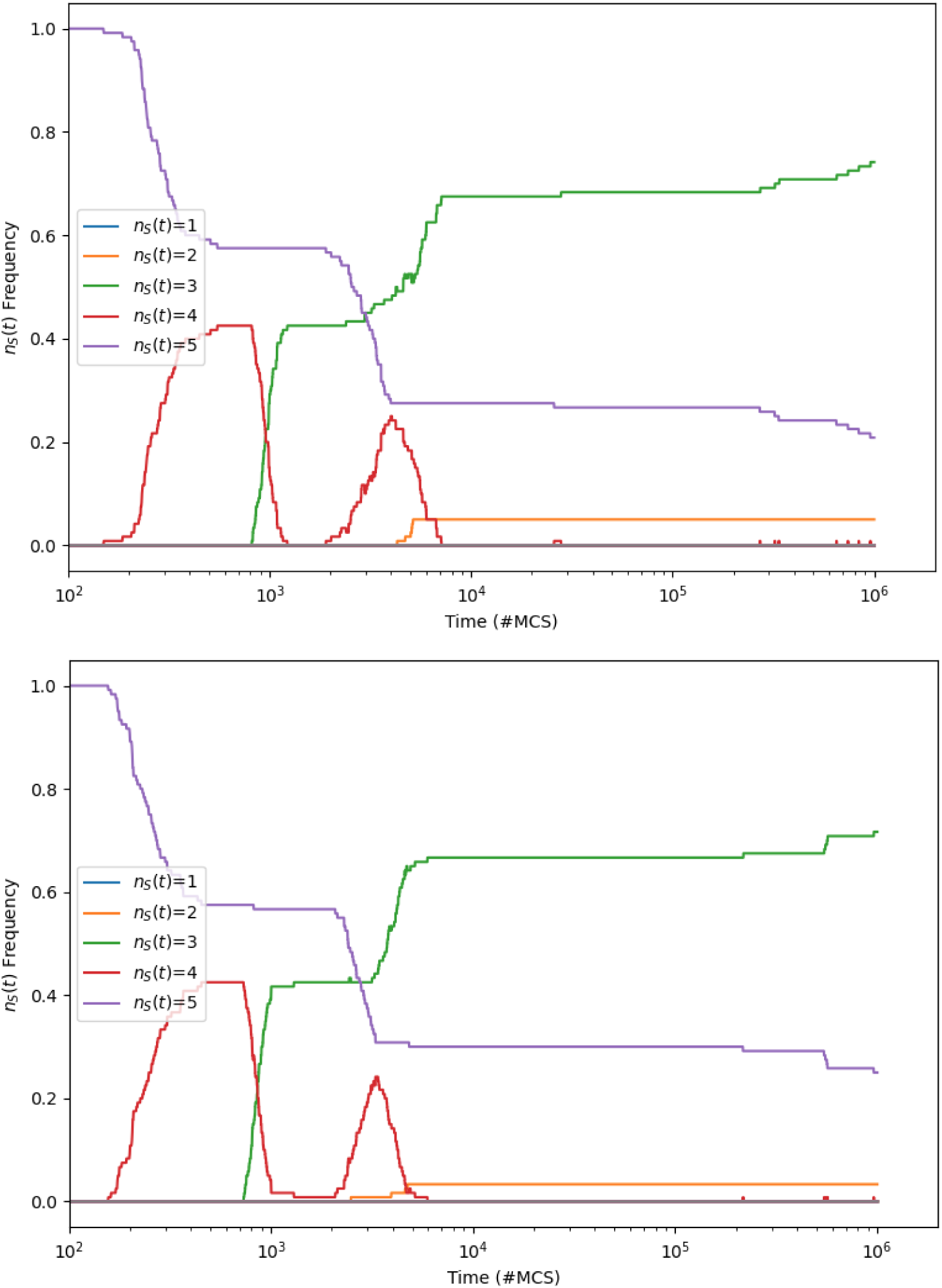
Frequency distribution of species-counts *n_s_*(*t*) against time *t* for Rpsls experiments with three ablations (*N_a_*=3): *L*=200; *µ*=*σ*=1.0; *M* =1.58×10*^−^*^6^ (upper graph), *M* =2.51×10*^−^*^6^ (lower graph). At each value of *M* sampled, 250 IID experiments were performed. Format as for Fig. A.23.

**Figure B.34:**
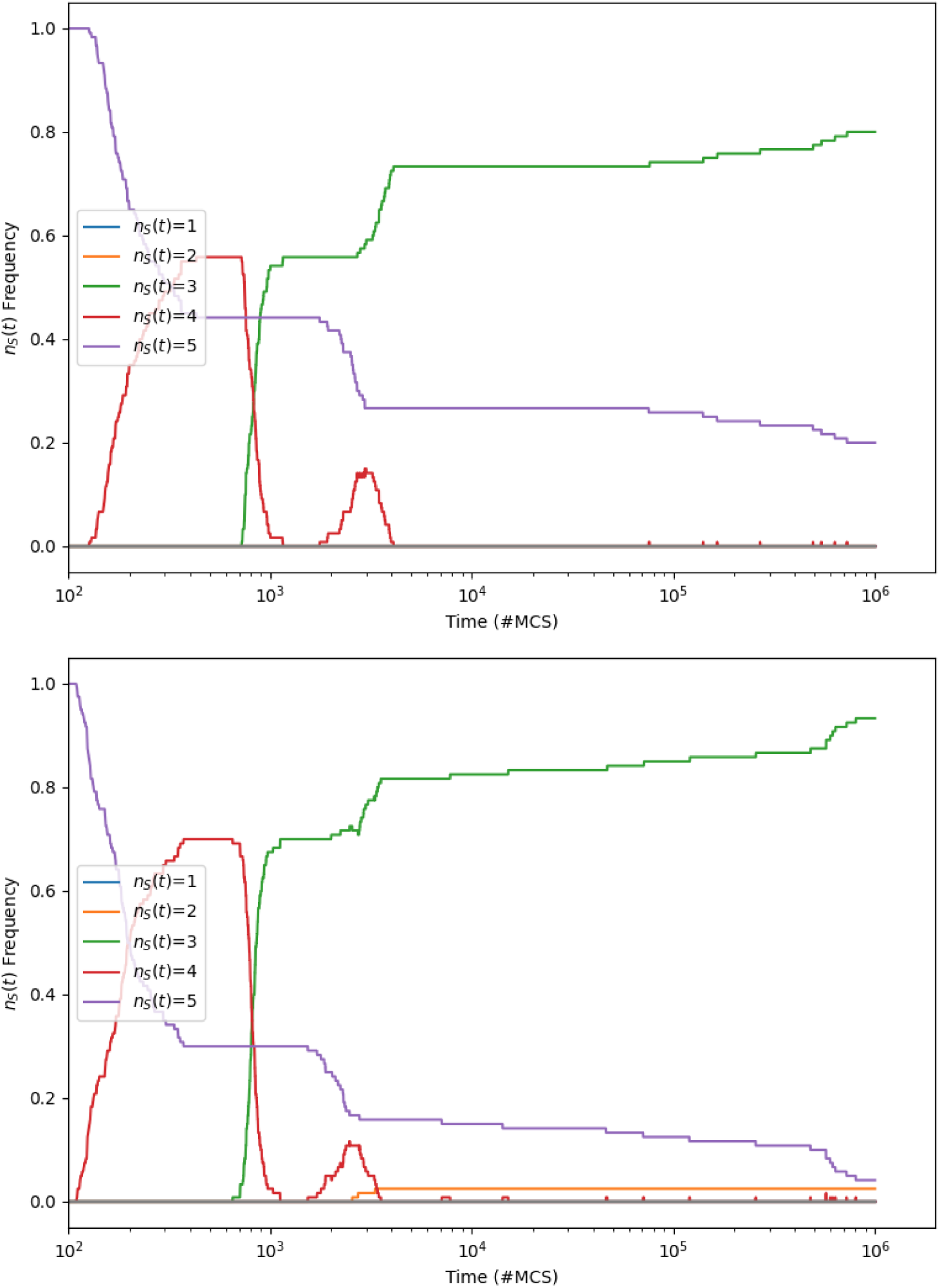
Frequency distribution of species-counts *n_s_*(*t*) against time *t* for Rpsls experiments with three ablations (*N_a_*=3): *L*=200; *µ*=*σ*=1.0; *M* =3.98×10*^−^*^6^ (upper graph), *M* =6.31×10*^−^*^6^ (lower graph). At each value of *M* sampled, 250 IID experiments were performed. Format as for Fig. A.23.

**Figure B.35:**
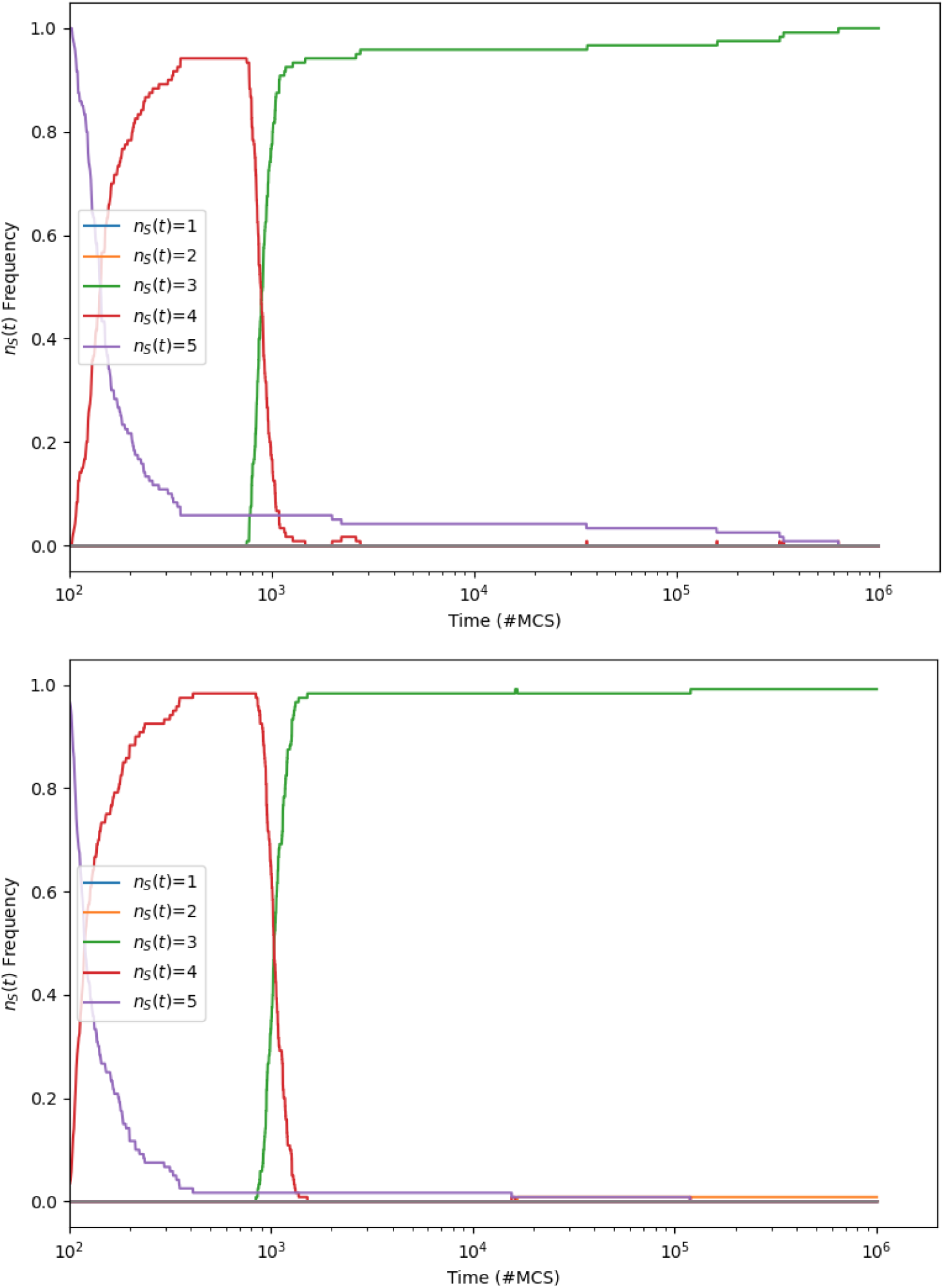
Frequency distribution of species-counts *n_s_*(*t*) against time *t* for Rpsls experiments with three ablations (*N_a_*=3): *L*=200; *µ*=*σ*=1.0; *M* =10*^−^*^5^ (upper graph), *M* =1.58×10*^−^*^5^ (lower graph). At each value of *M* sampled, 250 IID experiments were performed. Format as for Fig. A.23.

**Figure B.36:**
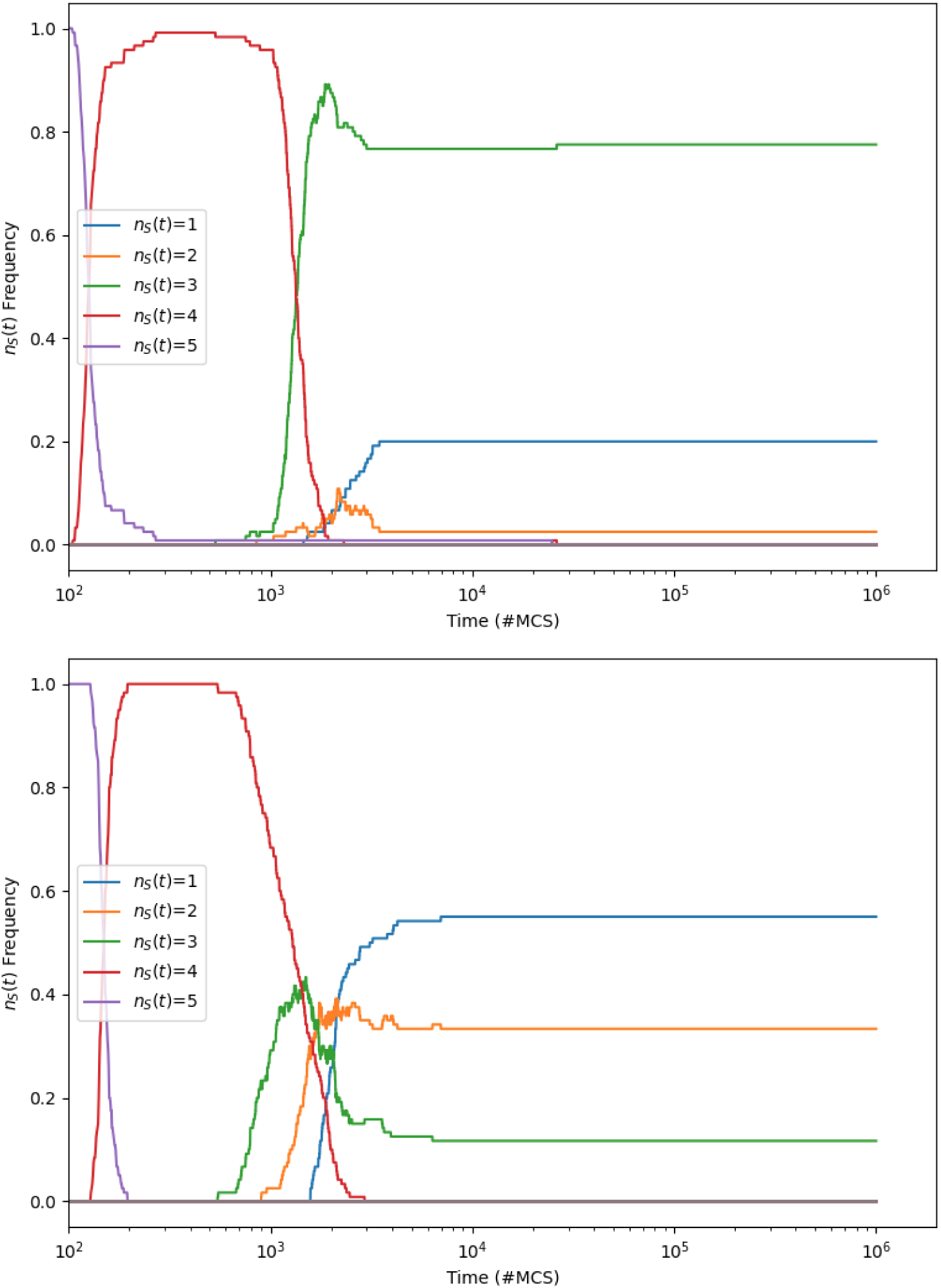
Frequency distribution of species-counts *n_s_*(*t*) against time *t* for Rpsls experiments with three ablations (*N_a_*=3): *L*=200; *µ*=*σ*=1.0; *M* =2.51×10*^−^*^5^ (upper graph), *M* =3.98×10*^−^*^5^ (lower graph). At each value of *M* sampled, 250 IID experiments were performed. Format as for Fig. A.23.

**Figure B.37:**
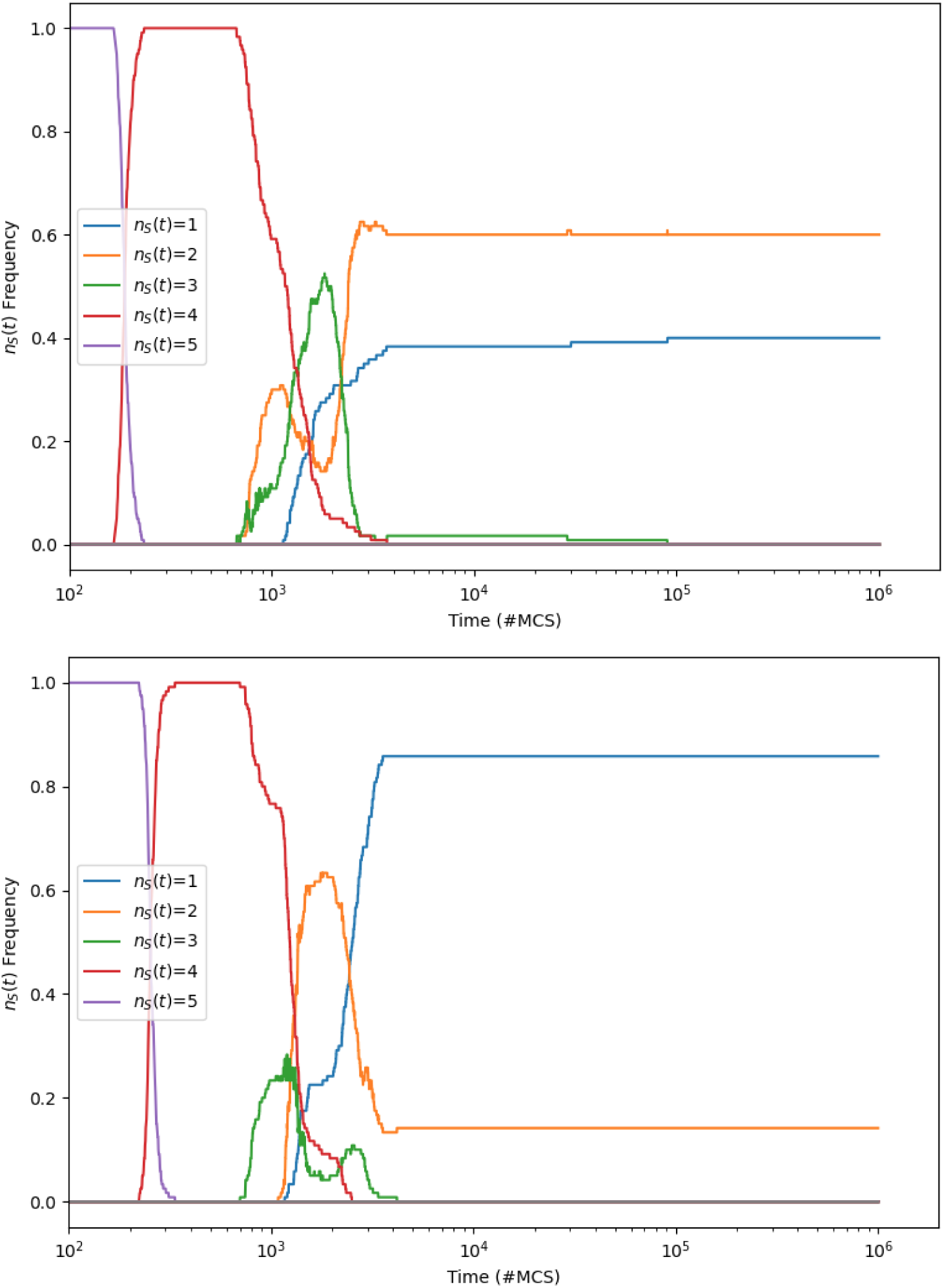
Frequency distribution of species-counts *n_s_*(*t*) against time *t* for Rpsls experiments with three ablations (*N_a_*=3): *L*=200; *µ*=*σ*=1.0; *M* =6.31 10*^−^*^5^ (upper graph), *M* =10*^−^*^4^ (lower graph). At each value of *M* sampled, 250 IID experiments were performed. Format as for Fig. A.23.

**Figure B.38:**
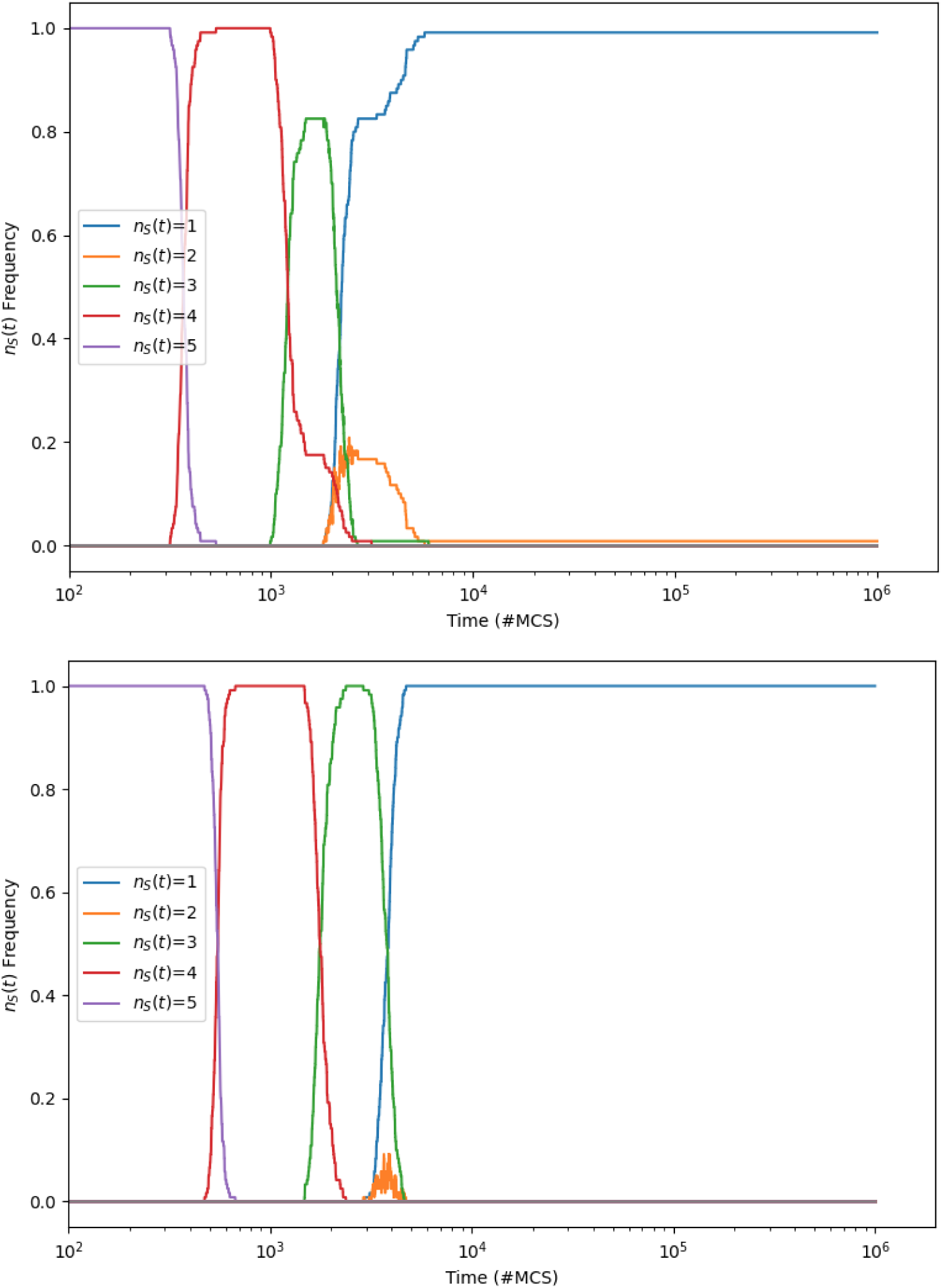
Frequency distribution of species-counts *n_s_*(*t*) against time *t* for Rpsls experiments with three ablations (*N_a_*=3): *L*=200; *µ*=*σ*=1.0; *M* =1.58 10*^−^*^4^ (upper graph), *M* =2.51 10*^−^*^4^ (lower graph). At each value of *M* sampled, 250 IID experiments were performed. Format as for Fig. A.23.

**Figure B.39:**
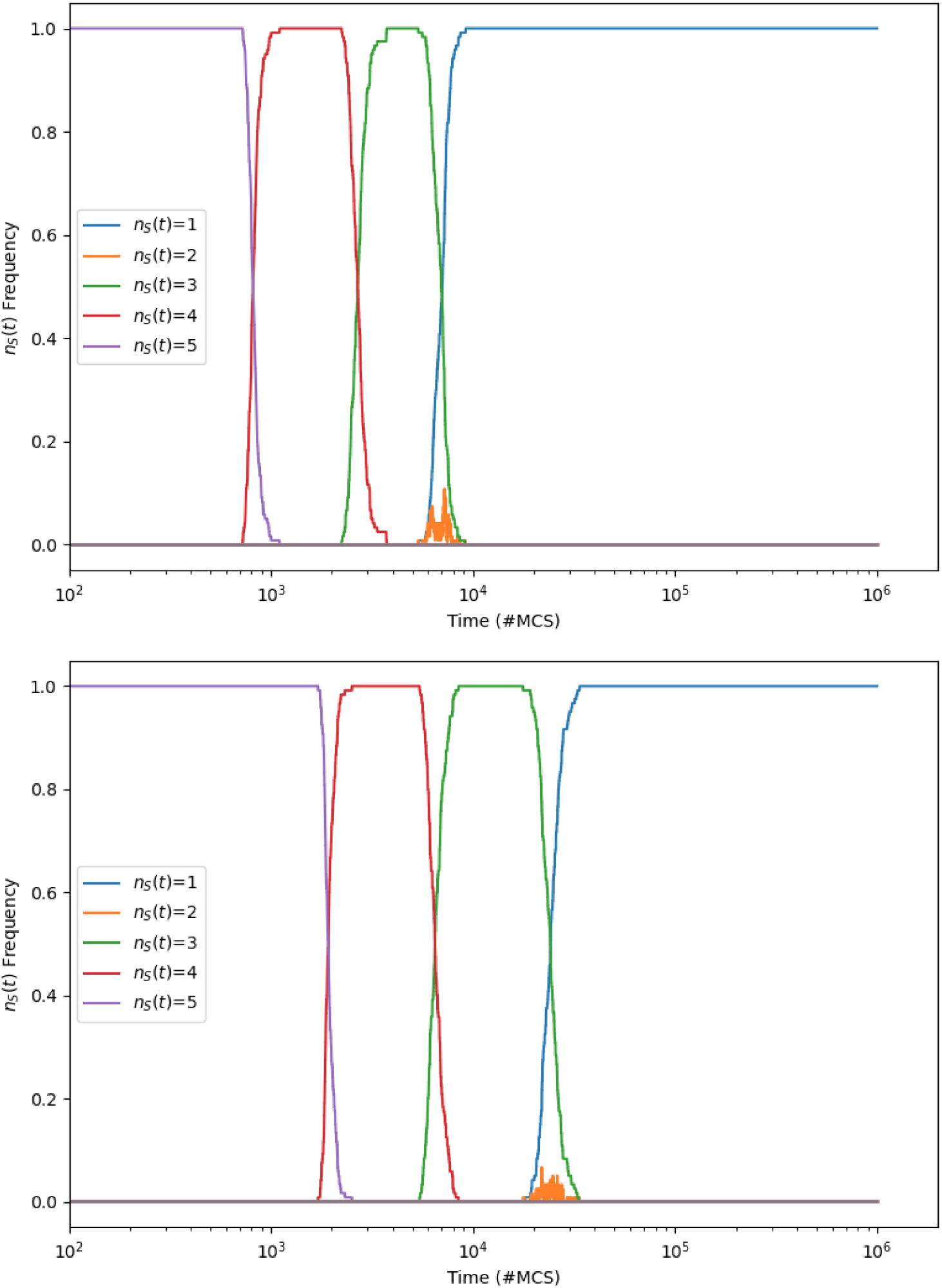
Frequency distribution of species-counts *n_s_*(*t*) against time *t* for Rpsls experiments with three ablations (*N_a_*=3): *L*=200; *µ*=*σ*=1.0; *M* =3.98 10*^−^*^4^ (upper graph), *M* =10*^−^*^3^ (lower graph). At each value of *M* sampled, 250 IID experiments were performed. Format as for Fig. A.23.

## Appendix C. Dynamics of *N_a_*=4 Rpsls Z4 with *µ*=*σ*=1.0, *L*=200

Figures C.41 to C.49 show time-series of the evolution of the number of surviving species *n_s_*(*t*) over 10^6^mcs for the Rpsls Escg with four-ablation (*N_a_*=4) dominance network Z4 (as illustrated in Figure 3), *L*=200 and *µ*=*σ*=1.0 for various values of *M* ∈ [10*^−^*^7^, 10*^−^*^3^], with 250 IID simulations executed for each value of *M* sampled.

**Figure C.40:**
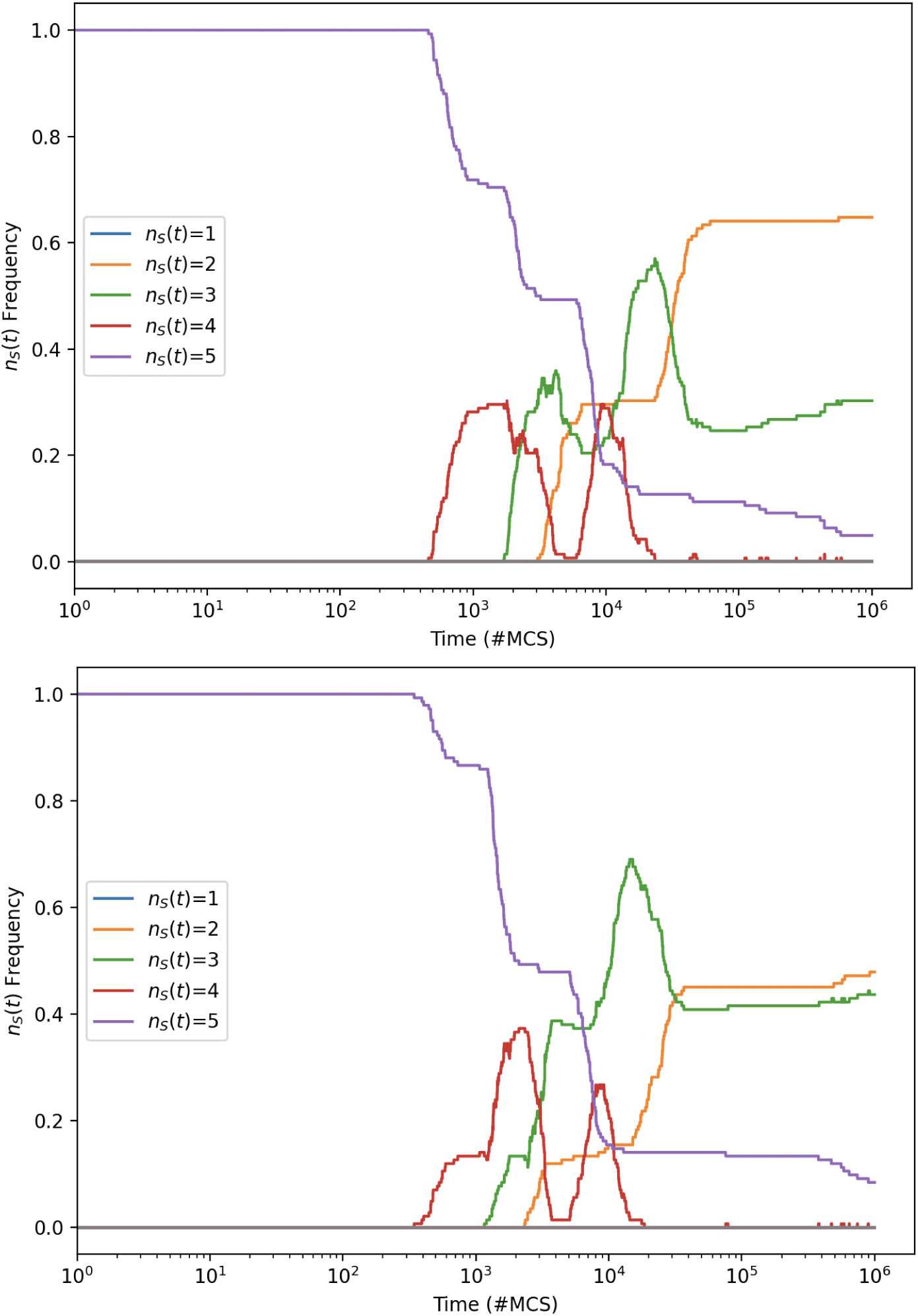
Frequency distribution of species-counts *n_s_*(*t*) : *s* ∈ {1*, . . .,* 5} against time *t* (measured in mcs) for Rpsls experiments with four ablations (*N_a_*=4): *L*=200; *µ*=*σ*=1.0; *M* =10*^−^*^7^ (upper graph), *M* =1.58 10*^−^*^7^ (lower graph). At each value of *M* sampled, 250 IID simulation experiments were performed. Format as for Fig. A.23.

**Figure C.41:**
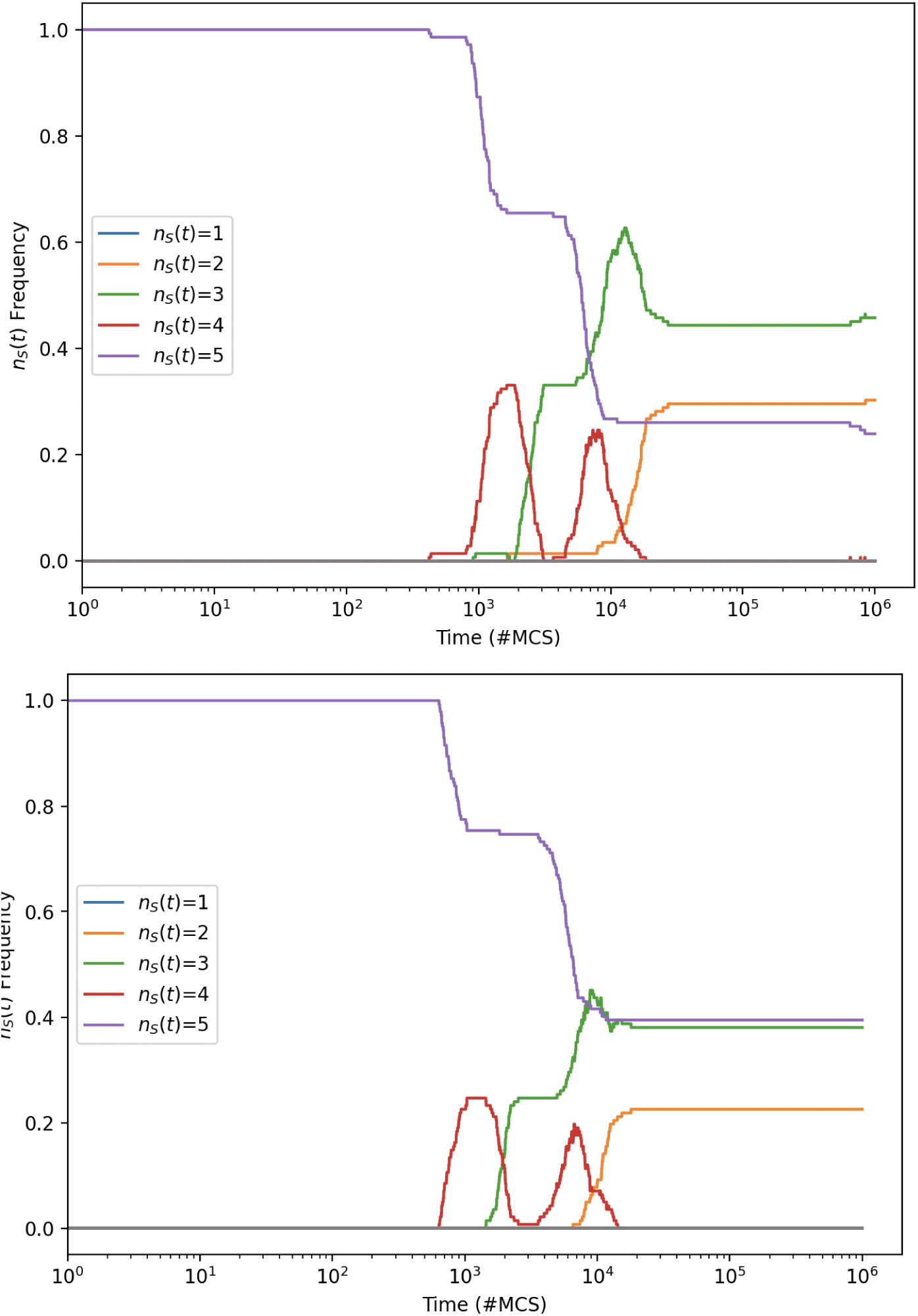
Frequency distribution of species-counts *n_s_*(*t*) against time *t* for Rpsls experiments with four ablations (*N_a_*=4): *L*=200; *µ*=*σ*=1.0; *M* =2.51 10*^−^*^7^ (upper graph), *M* =3.98 10*^−^*^7^ (lower graph). At each value of *M* sampled, 250 IID experiments were performed. Format as for Fig. A.23.

**Figure C.42:**
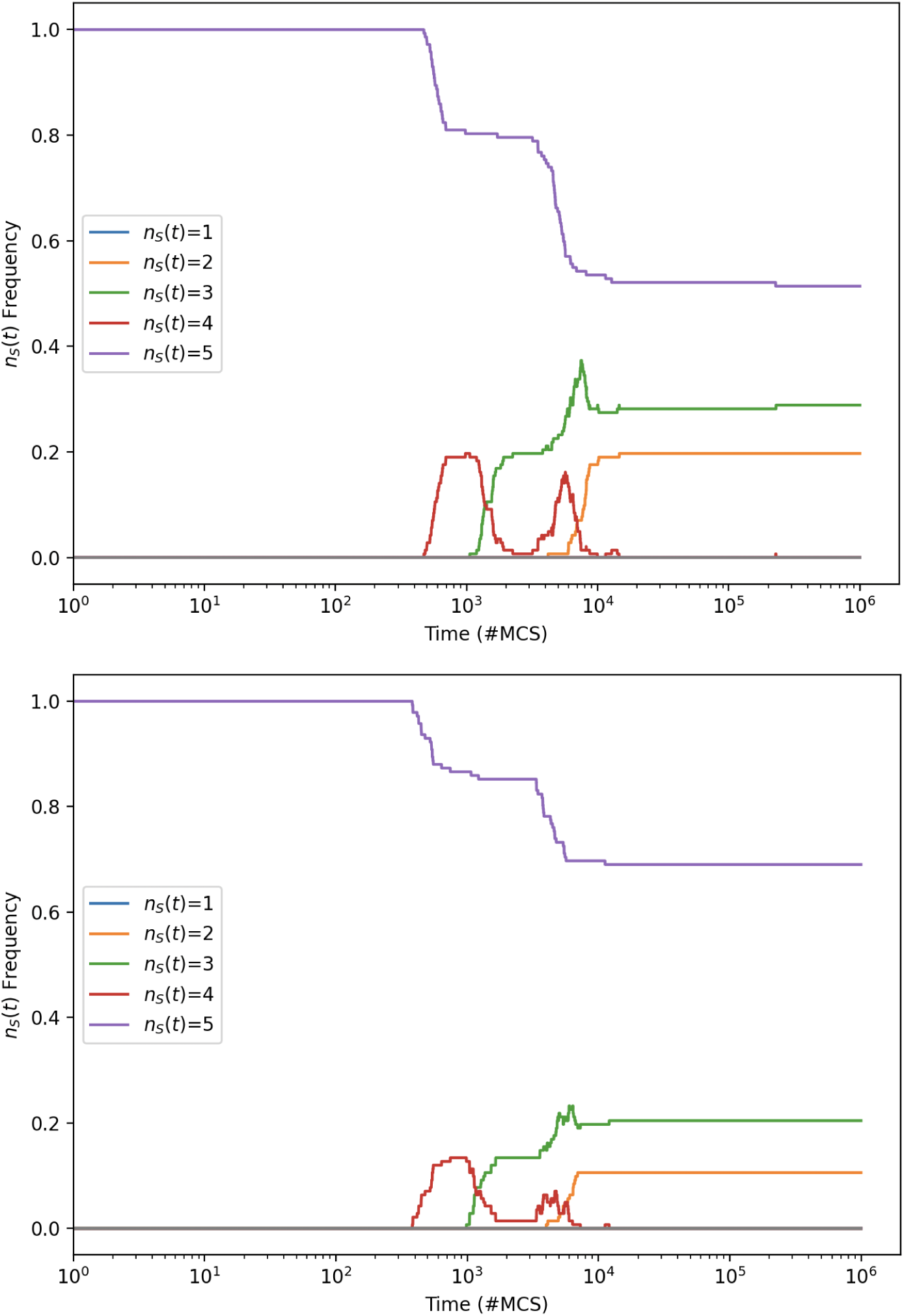
Frequency distribution of species-counts *n_s_*(*t*) against time *t* for Rpsls experiments with four ablations (*N_a_*=4): *L*=200; *µ*=*σ*=1.0; *M* =6.31 10*^−^*^7^ (upper graph), *M* =10*^−^*^6^ (lower graph). At each value of *M* sampled, 250 IID experiments were performed. Format as for Fig. A.23.

**Figure C.43:**
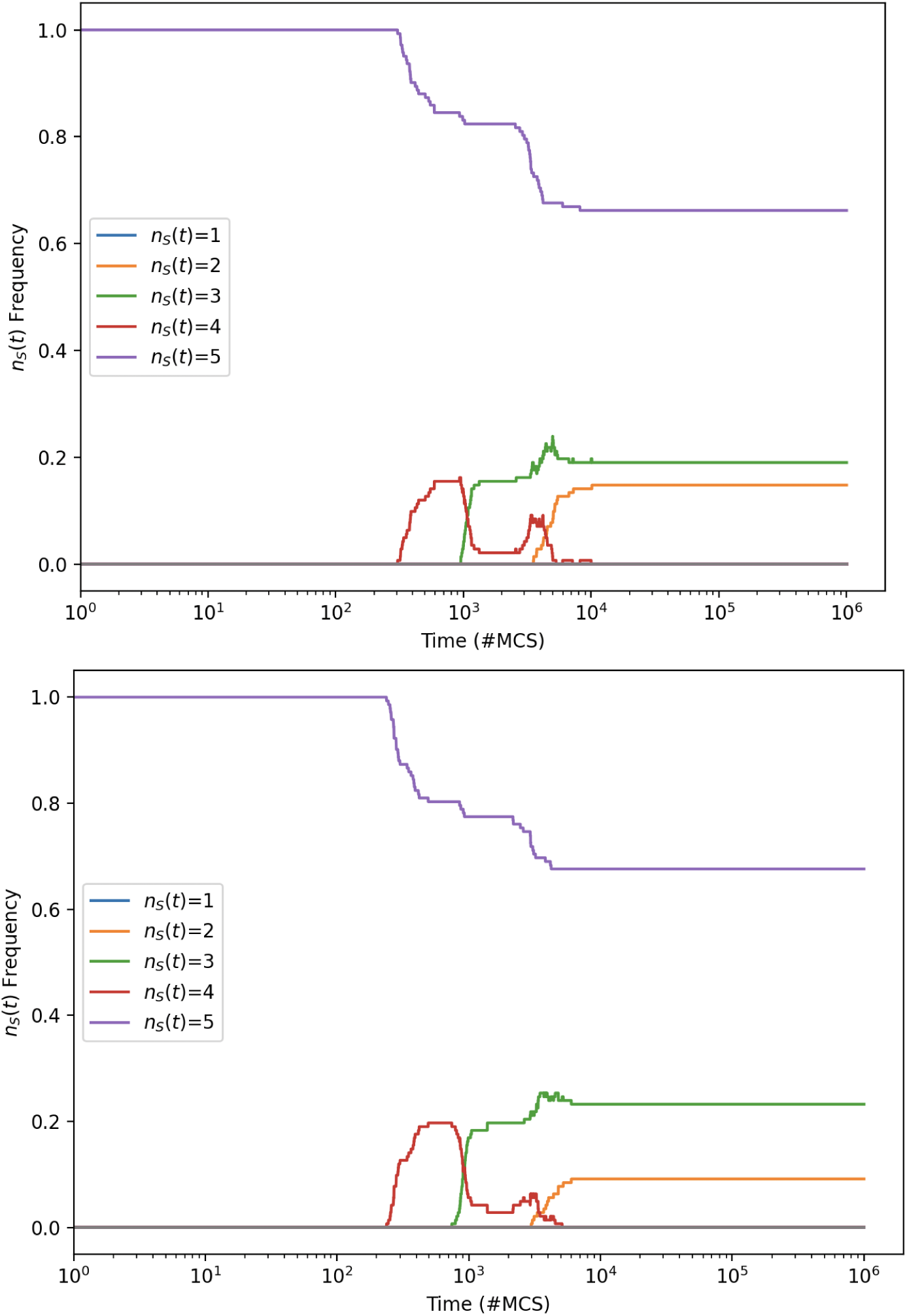
Frequency distribution of species-counts *n_s_*(*t*) against time *t* for Rpsls experiments with four ablations (*N_a_*=4): *L*=200; *µ*=*σ*=1.0; *M* =1.58×10*^−^*^6^ (upper graph), *M* =2.51×10*^−^*^6^ (lower graph). At each value of *M* sampled, 250 IID experiments were performed. Format as for Fig. A.23.

**Figure C.44:**
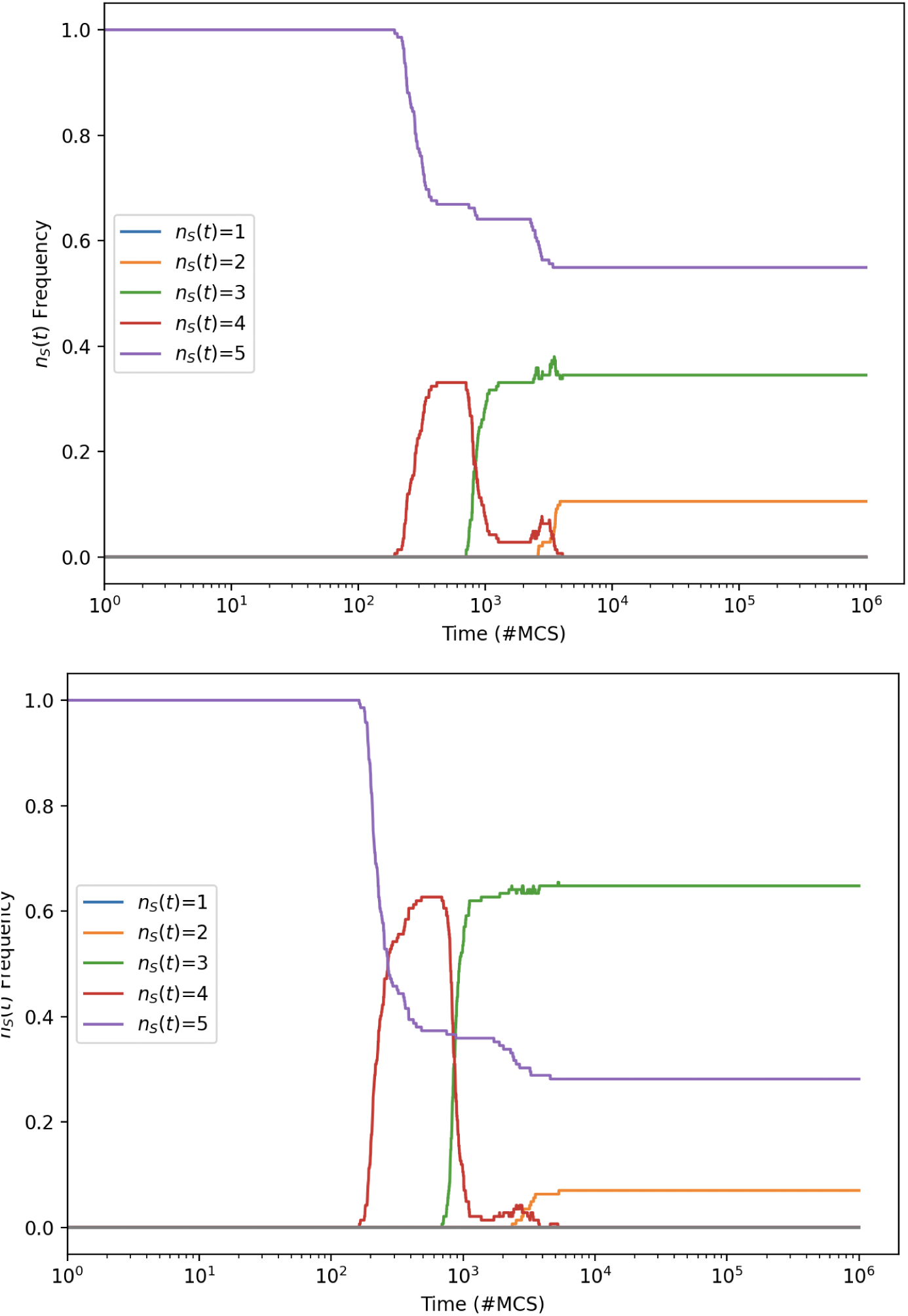
Frequency distribution of species-counts *n_s_*(*t*) against time *t* for Rpsls experiments with four ablations (*N_a_*=4): *L*=200; *µ*=*σ*=1.0; *M* =3.98×10*^−^*^6^ (upper graph), *M* =6.31×10*^−^*^6^ (lower graph). At each value of *M* sampled, 250 IID experiments were performed. Format as for Fig. A.23.

**Figure C.45:**
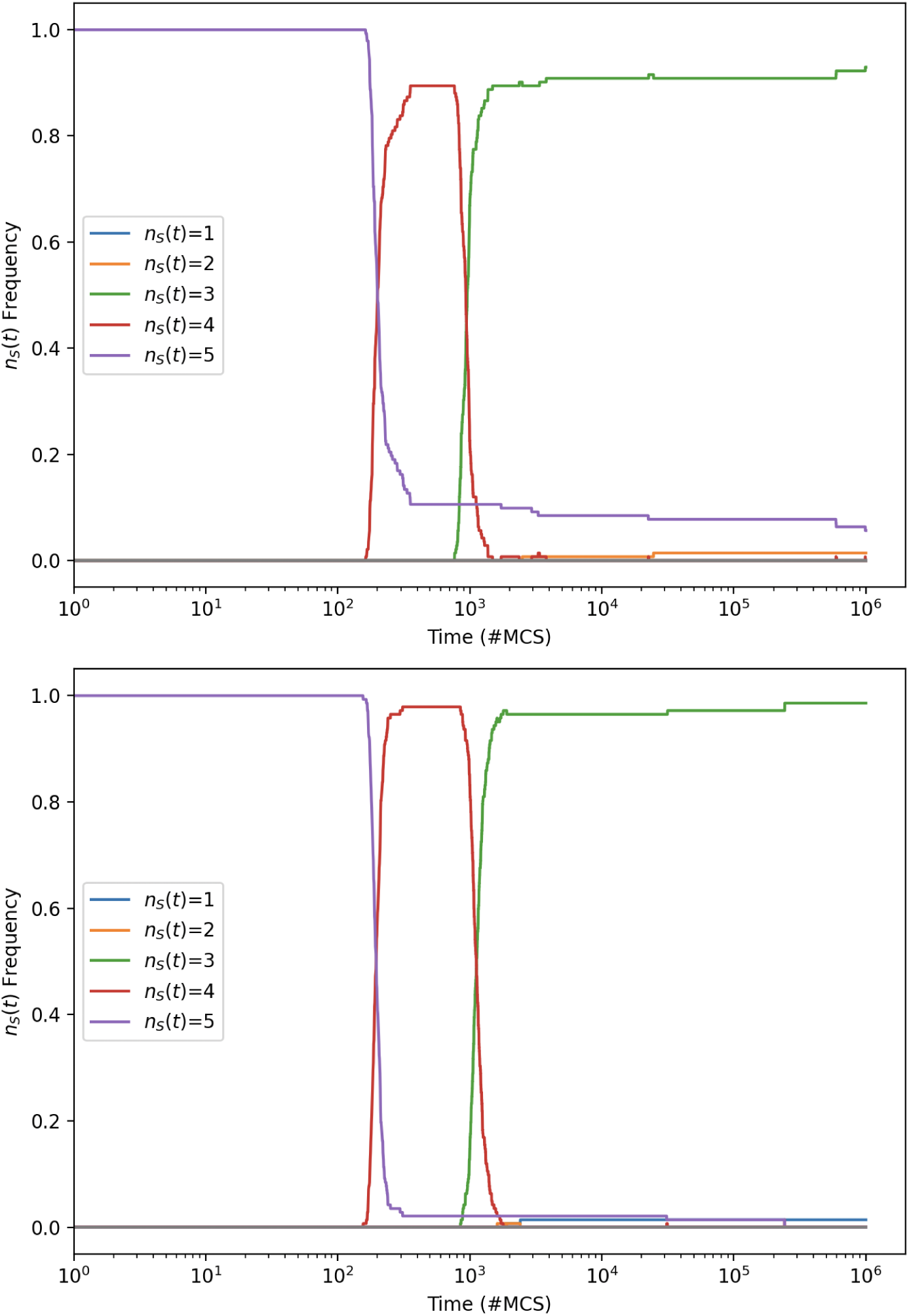
Frequency distribution of species-counts *n_s_*(*t*) against time *t* for Rpsls experiments with four ablations (*N_a_*=4): *L*=200; *µ*=*σ*=1.0; *M* =10*^−^*^5^ (upper graph), *M* =1.58×10*^−^*^5^ (lower graph). At each value of *M* sampled, 250 IID experiments were performed. Format as for Fig. A.23.

**Figure C.46:**
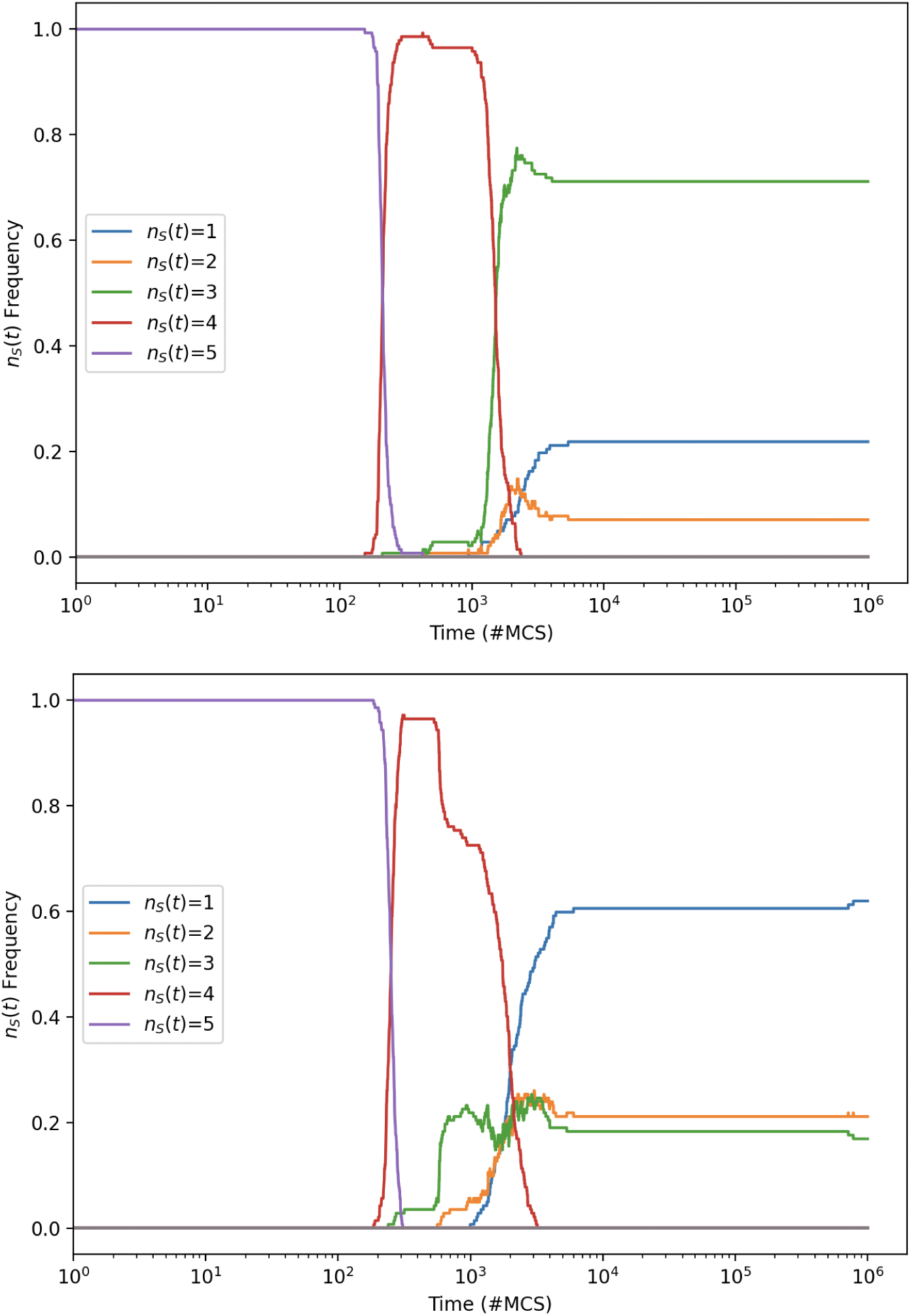
Frequency distribution of species-counts *n_s_*(*t*) against time *t* for Rpsls experiments with four ablations (*N_a_*=4): *L*=200; *µ*=*σ*=1.0; *M* =2.51×10*^−^*^5^ (upper graph), *M* =3.98×10*^−^*^5^ (lower graph). At each value of *M* sampled, 250 IID experiments were performed. Format as for Fig. A.23.

**Figure C.47:**
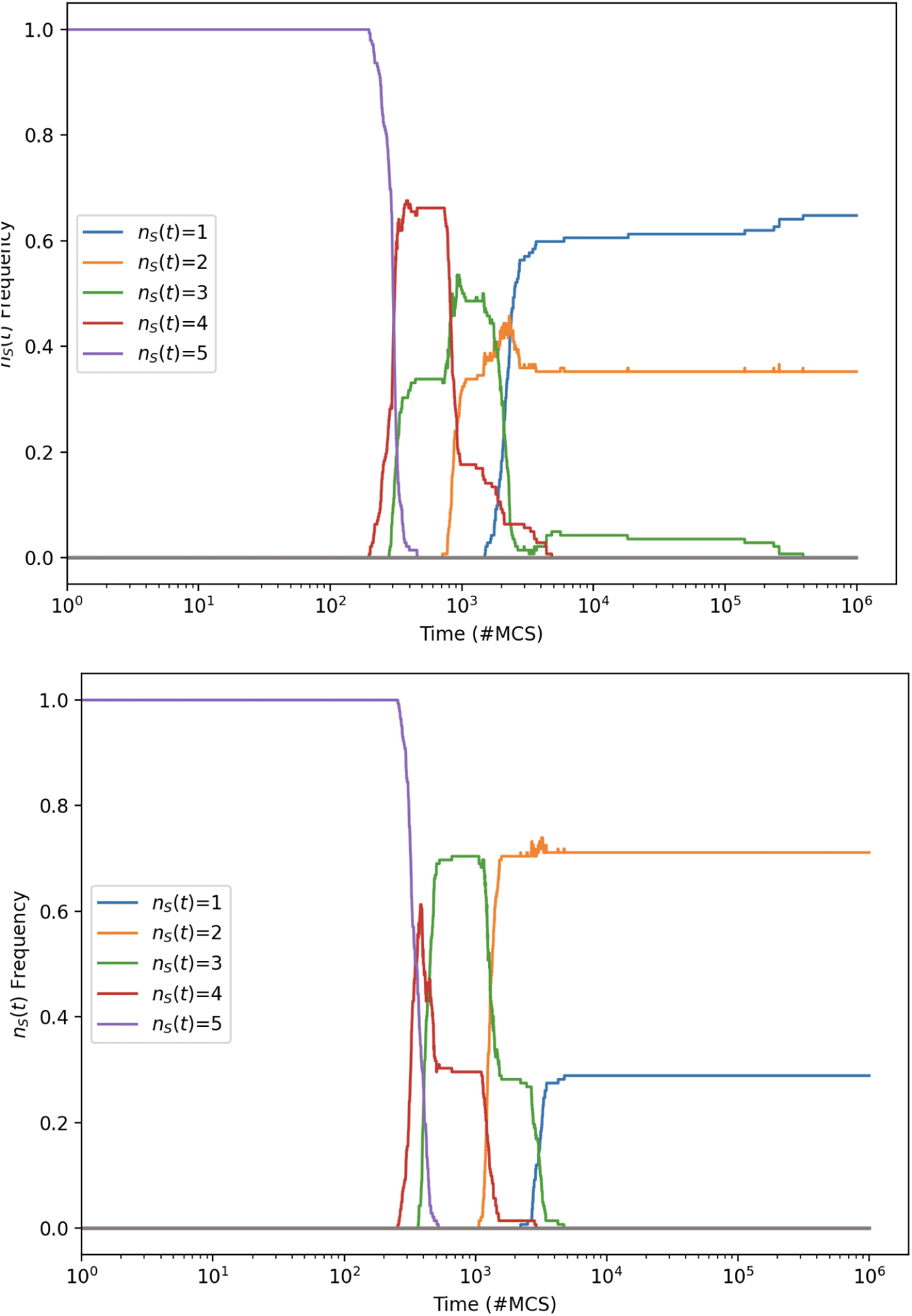
Frequency distribution of species-counts *n_s_*(*t*) against time *t* for Rpsls experiments with four ablations (*N_a_*=4): *L*=200; *µ*=*σ*=1.0; *M* =6.31 10*^−^*^5^ (upper graph), *M* =10*^−^*^4^ (lower graph). At each value of *M* sampled, 250 IID experiments were performed. Format as for Fig. A.23.

**Figure C.48:**
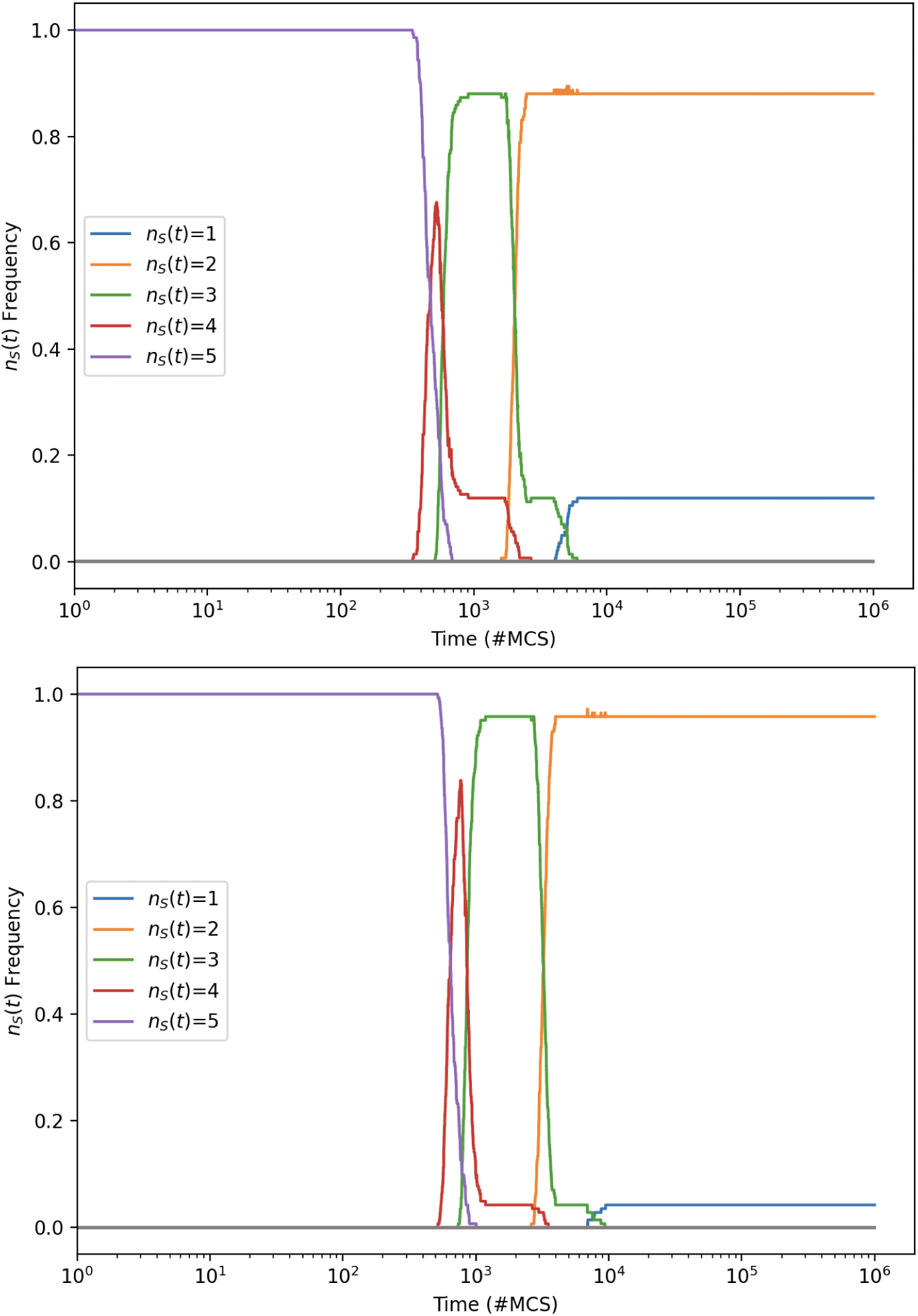
Frequency distribution of species-counts *n_s_*(*t*) against time *t* for Rpsls experiments with four ablations (*N_a_*=4): *L*=200; *µ*=*σ*=1.0; *M* =1.58 10*^−^*^4^ (upper graph), *M* =2.51 10*^−^*^4^ (lower graph). At each value of *M* sampled, 250 IID experiments were performed. Format as for Fig. A.23.

**Figure C.49:**
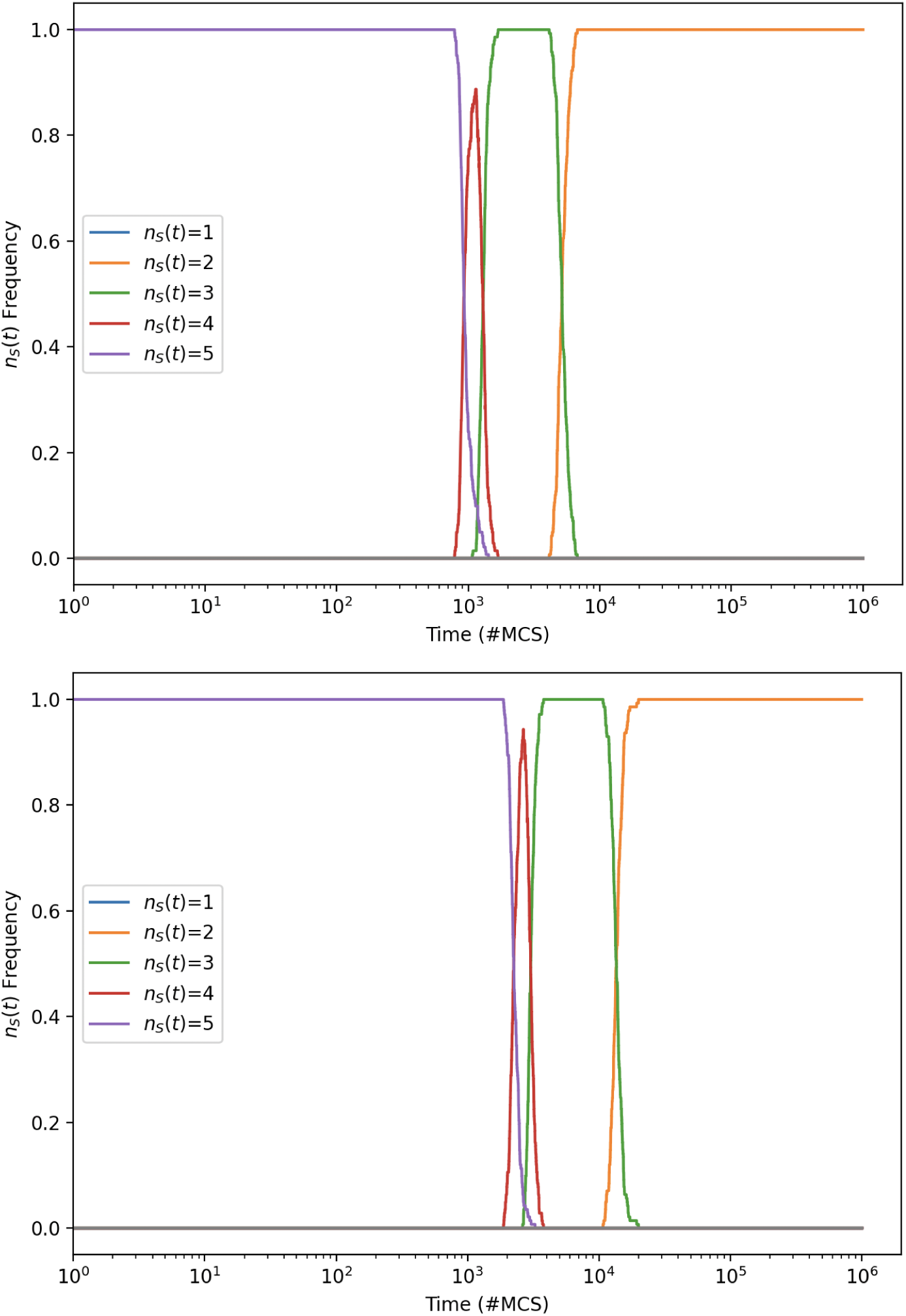
Frequency distribution of species-counts *n_s_*(*t*) against time *t* for Rpsls experiments with four ablations (*N_a_*=4): *L*=200; *µ*=*σ*=1.0; *M* =3.98 10*^−^*^4^ (upper graph), *M* =10*^−^*^3^ (lower graph). At each value of *M* sampled, 250 IID experiments were performed. Format as for Fig. A.23.

## Footnotes

1 The terminology used in the literature varies: some authors such as [2] refer to the dominance network as the *interaction graph*, others such as [3] refer to it instead as the *competitive matrix*.

2 See https://github.com/davecliff/ESCGPython.

3 Many central processor unit (CPU) chips include a no-op instruction in their assembly language: typically the no-op will consume exactly one unit of clock time while leaving everything else unchanged in the CPU and its associated memory. No-ops can be useful in parallel real-time systems where it is important to establish or maintain temporal synchronisation between the constituent sub-systems.

4 These re-printed graphs are screen-shots taken from the PDF of [1] and are reproduced here, as quotations, in the belief that this usage qualifies legally as ‘fair dealing’ for non-commercial research as defined in Section 29 of the UK Copyright, Designs and Patents Act 1988 (CDPA1988), and/or as ‘fair dealing’ for criticism, review, quotation, and reporting as defined in Section 30 of CDPA1988.

5 Writing in [37], Milkowski et al. make the distinction that *replicability* of a computational simulation or modelling experiment involves independent researchers obtaining the same output as the original experiment, using the experiment’s original code and data, whereas *reproducibility* of a computational model or experiment involves independent researchers being able to recreate the results *without* access to the original code and data. Using that distinction strictly, my work reported in this paper demonstrates the reproducibility of Zhong et al.’s results, but is not a replication of those results because I wrote all my own code from scratch.

